# A marine bacterial community that degrades poly(ethylene terephthalate) and polyethylene

**DOI:** 10.1101/2020.11.07.372490

**Authors:** Rongrong Gao, Chaomin Sun

## Abstract

Plastic wastes have become the most common form of marine debris and present a growing global pollution problem. Recently, microorganisms-mediated degradation has become a most promising way to accomplish the eventual bioremediation of plastic wastes due to their prominent degradation potentials. Here, a marine bacterial community which could efficiently colonize and degrade both poly (ethylene terephthalate) (PET) and polyethylene (PE) was discovered through a screening with hundreds of plastic waste associated samples. Using absolute quantitative 16S rRNA sequencing and cultivation methods, we obtained the abundances and pure cultures of three bacteria mediating plastic degradation. We further reconstituted a tailored bacterial community containing above three bacteria and demonstrated its efficient degradation of PET and PE through various techniques. The released products from PET and PE degraded by the reconstituted bacterial community were determined by the liquid chromatography-mass spectrometry. Finally, the plastic degradation process and potential mechanisms mediated by the reconstituted bacterial community were elucidated through transcriptomic methods. Overall, this study establishes a stable and effective marine bacterial community for PET and PE degradation and sheds light on the degradation pathways and associated mechanistic processes, which paves a way to develop a microbial inoculant against plastic wastes.

## Introduction

Plastics have been found widespread in the world’s oceans^1–4^. It has been reported that about 4.8 to 12.7 million tons of plastic debris per year enter the ocean^5^. Plastics in the marine environment are of increasing concern because of their persistence and effects on the oceans^6^, wildlife^7^, and, potentially, humans^3^. An estimated one million birds and ten thousand marine animals die each year as a result of ingestion of or trapping by plastics in the oceans^8, 9^. Moreover, after weathering, mechanical wear and ultraviolet radiation, the large plastic may be broken into fragmentation, when it is smaller than 5 mm in diameter, it was commonly defined as microplastics^10^. Of note, microplastics also negatively impact upon marine biota and can be ingested and accumulated along trophic webs until top predators^11, 12^.

Among various types of plastic wastes, poly (ethylene terephthalate) (PET) and polyethylene (PE) constitute the major 46.5% portion of the tremendous amount of plastic pollution debris^13^. PET is a type of semi-aromatic thermoplastic co-polymer resin from polyester family, which has aromatic groups heteroatoms in the main chain^14^. PE has a carbon-carbon backbone which is solely built of carbon atoms and has high resistance against various degradation processes, due to non-hydrolyzable covalent bonds^15^. Both PET and PE have properties such as lower density, long hydrocarbon chain, high molecular weight and tensile strength, low permeability to gases, durability to physical and chemical degradation, non-biodegradable compound^16–18^.

Landfilling, incineration, recycling and biodegradation are the principal strategies to solve the plastic waste problem^8^. As an environmentally friendly alternative to conventional plastic waste management methods, microorganisms-mediated degradation is the most promising way to accomplish an eventual bioremediation of plastic wastes. Microbial degradation of plastics is usually an enzymatic activity that catalyzes the cleavage of polymer bond into monomer entity^19, 20^. Thus far, PET hydrolyzing activity has been reported for members of the cutinase, lipase and esterase^20^. PE, as one of the most abundant plastics in the ocean, shows obvious signs of degradation when incubated with specific microorganisms under controlled laboratory conditions^21–23^. However, as compared to the extensive studies about PET-degrading bacteria and enzymes^14, 18, 19, 24–26^, the researches about PE degradation mediated by microorganisms lag well behind and the degradation efficiency using a single strain or enzyme is still too low to meet the industrial applications requirements. Alternatively, using microbial community to degrade PE might be a good choice given their inherent multiple robust function among synergistic effect of different species^27^. Actually, the construction of artificial microbial consortia has opened a new horizon in environment bioremediation in terms of removing hard biodegradable harmful compounds^28^.

Herein, a marine bacterial community efficiently degrading both PET and PE was obtained by a large-scale screening. Three bacteria driving plastic degradation were isolated and reconstituted to an artificial bacterial community with a similar degradation capability to that of the original bacterial flora. The degradation effects and possible products of PET and PE treated by this reconstituted bacterial community were further clarified by various techniques. Lastly, the potential degradation process and associated enzymes were disclosed through transcriptomics methods.

## Results

### Discovery of a marine bacterial community efficiently degrading both PET and PE

To obtain potential marine bacteria degrading PET or PE, we collected about 300 sediment samples contaminated by plastic debris from different locations of a bay of China. Using these samples, we initiated to screen microorganisms that could use plastic drink bottles (whose main component is PET) or commercial PE bags as major carbon sources for growth. With that, a distinct consortium derived from one of the plastic debris samples could efficiently colonize on both PET (Supplementary Fig. 1b) and PE films (Supplementary Fig. 1d). Scanning electronic microscopy (SEM) observation confirmed that the consortium could evidently colonize on PET (Supplementary Fig. 1f) and PE films (Supplementary Fig.1i). Of note, these colonizers formed an obvious biofilm layer and closely interacted each other with filament-like structures on PET (Supplementary Fig. 1g) and PE films (Supplementary Fig.1j). After removing the microbial layer from the films, significant morphological changes in both PET (Supplementary Figs. 2b-2d) and PE films were observed by SEM, especially for PE which showed large amount of heavy cracks and deep holes in the surface and even the inside of the film (Supplementary Figs. 2f-2h).

Given the fact that most commercial plastics contain various additives (such as dyes, plasticizer, antistatic agents etc.), it is necessary to make sure that the consortium indeed degraded the plastics rather than the additives. We thus repeated the degradation test by the above consortium with the PET and PE films without any additives. Consistently, both PET and PE films were evidently degraded after 4 weeks treatment by the consortium even observed by eyes (Figs. 1b, 1d), and the four corners of both films lost sharp morphology as that shown in the control (Figs. 1a, 1c). Similar to the results obtained with the additive-containing plastics, SEM observation revealed that the consortium could efficiently colonize on the surface of films (Figs. 1f, 1i) and caused obvious degradation by forming pits, cracks or holes in the surface and inside of PET (Fig. 1g) and PE film (Fig. 1j). Similar to additive-containing plastics, the consortium prefers to degrade pure PE than PET. Overall, we conclude that this consortium could efficiently degrade both PET and PE, and is worthy of further study.

**Fig. 1.**
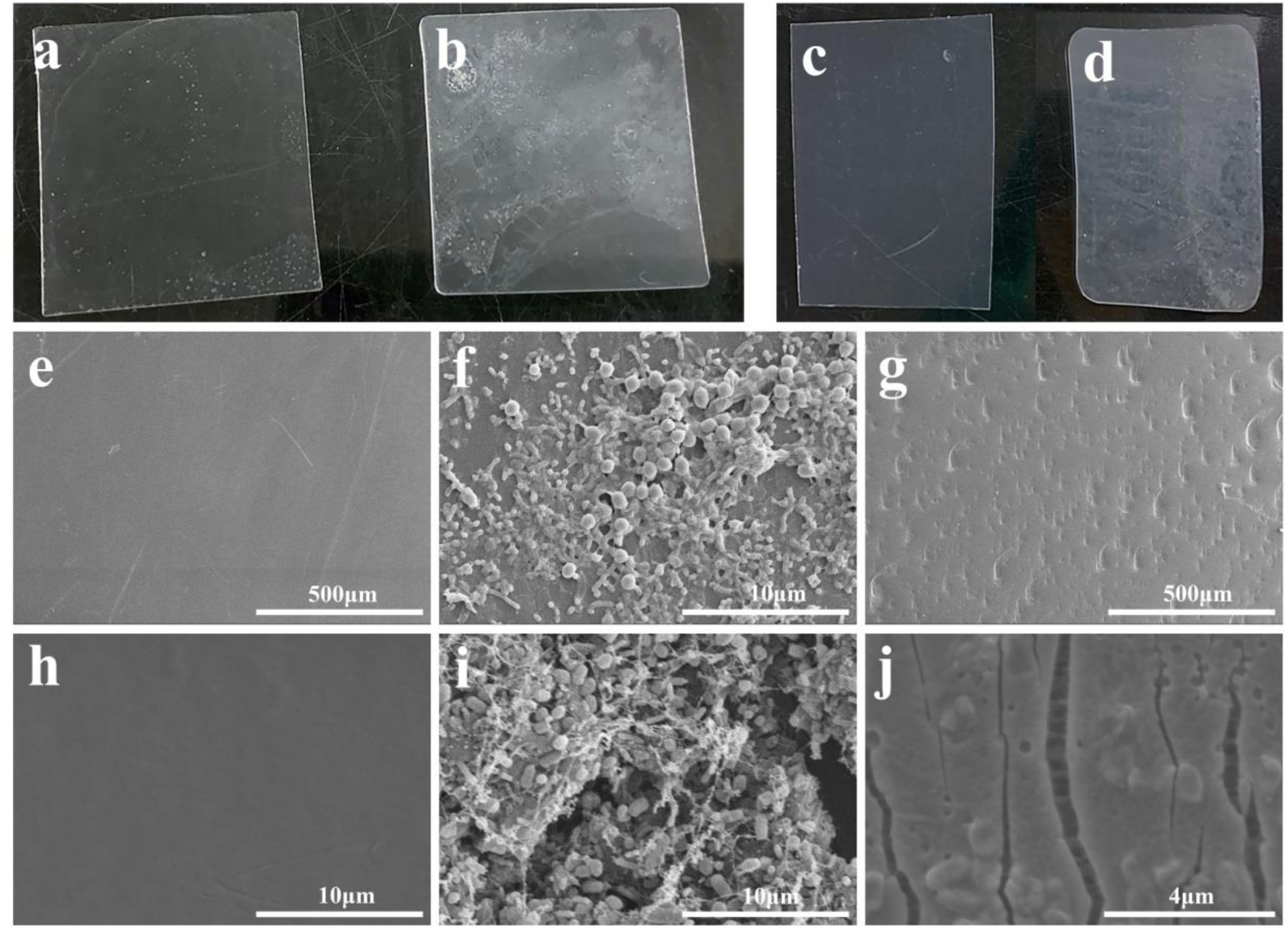
**Observation of the colonization and degradation effects of a marine bacterial community on PET and PE films**. **a**, Morphology of PET film without treatment. **b**, Morphology of PET film treated by the marine bacterial community for seven days. **c**, Morphology of PE film without treatment. **d**, Morphology of PE film treated by the marine bacterial community for seven days. **e**, SEM observation of PET film without treatment. **f**, SEM observation of the colonization of the marine bacterial community on the PET film after seven days incubation. **g**, SEM observation of the degradation effects of PET film treated by the marine bacterial community for seven days. **h**, SEM observation of PE film without treatment. **i**, SEM observation of the colonization of the marine bacterial community on the PE film after seven days incubation. **j**, SEM observation of the degradation effects of PE film treated by the marine bacterial community for seven days. The type of PET film used for this assay is ES301450 (0.25 mm in thickness). The type of PE film used for this assay is ET311350 (0.25mm in thickness).

### Isolation of the key degraders and reconstitution of the functional community capable of plastic degradation

To figure out the composition and dynamics of the above microbial community during the course of plastics degradation, we performed an absolute quantitative analysis of 16S rRNA sequences on this microbial flora. The growth curve of this microbial flora showed that it took about 10 h to enter the stationary phase and kept a stable population quantity after 7 d- or even longer incubation (Supplementary Fig. 3a). Meanwhile, we monitored the degradation effects on plastics in five different growth stages as shown in Supplementary Fig. 3a through SEM. The SEM observation showed that the thickness and density of the microbial layer on plastic films increased along with the culturing time (Supplementary Figs. 3b-3e). Obvious cracks in the films were observed after 7 days incubation (Supplementary Fig. 3f), indicating the plastics had been degraded in this stage. Therefore, we further performed an absolute quantitative analysis of 16S rRNA sequences of bacteria within the microbial community at the 7-day incubation stage. The sequencing results revealed five bacterial general ranked in the top 5 at this time phase, which were *Idiomarina* (∼50%), *Marinobacter* (∼28%), *Exiguobacterium* (∼18%), *Halomonas* (∼2%), *Ochrobactrum* (∼1%) (Fig. 2a and Supplementary Table 1). To obtain the pure cultures of these bacteria, the cells attached to the films were further isolated. As expected, bacterial strains belonging to above five genera were obtained. However, only three of them (*Exiguobacterium* sp., *Halomonas* sp.*, Ochrobactrum* sp.) showed both significant degradation capabilities against PET (Figs. 2b, 2d, 2f) and PE (Figs. 2c, 2e, 2g), indicating the other two bacteria might prefer to form biofilms on the surface but not degradation of plastics even though their high abundance within the bacterial community. Notably, SEM observations indicated that the mixture of three above bacteria had a greater degradation efficiency on both PET (Fig. 2h) and PE (Fig. 2i), suggesting that the bacterial community had a better capability than single isolate for plastics degradation. With this, we reconstituted a novel bacterial community containing these three bacteria in the proportion 1:1:1 and further tested its plastic degradation efficiency. Surprisingly, after two weeks incubation, both PET (Fig. 3b) and PE (Fig. 3d) were totally degraded to small pieces, especially for PE. SEM observations further confirmed that the reconstituted bacterial community had good colonization and degradation abilities on both PET (Figs. 3e-3g) and PE (Figs. 3h-3j).

**Fig. 2.**
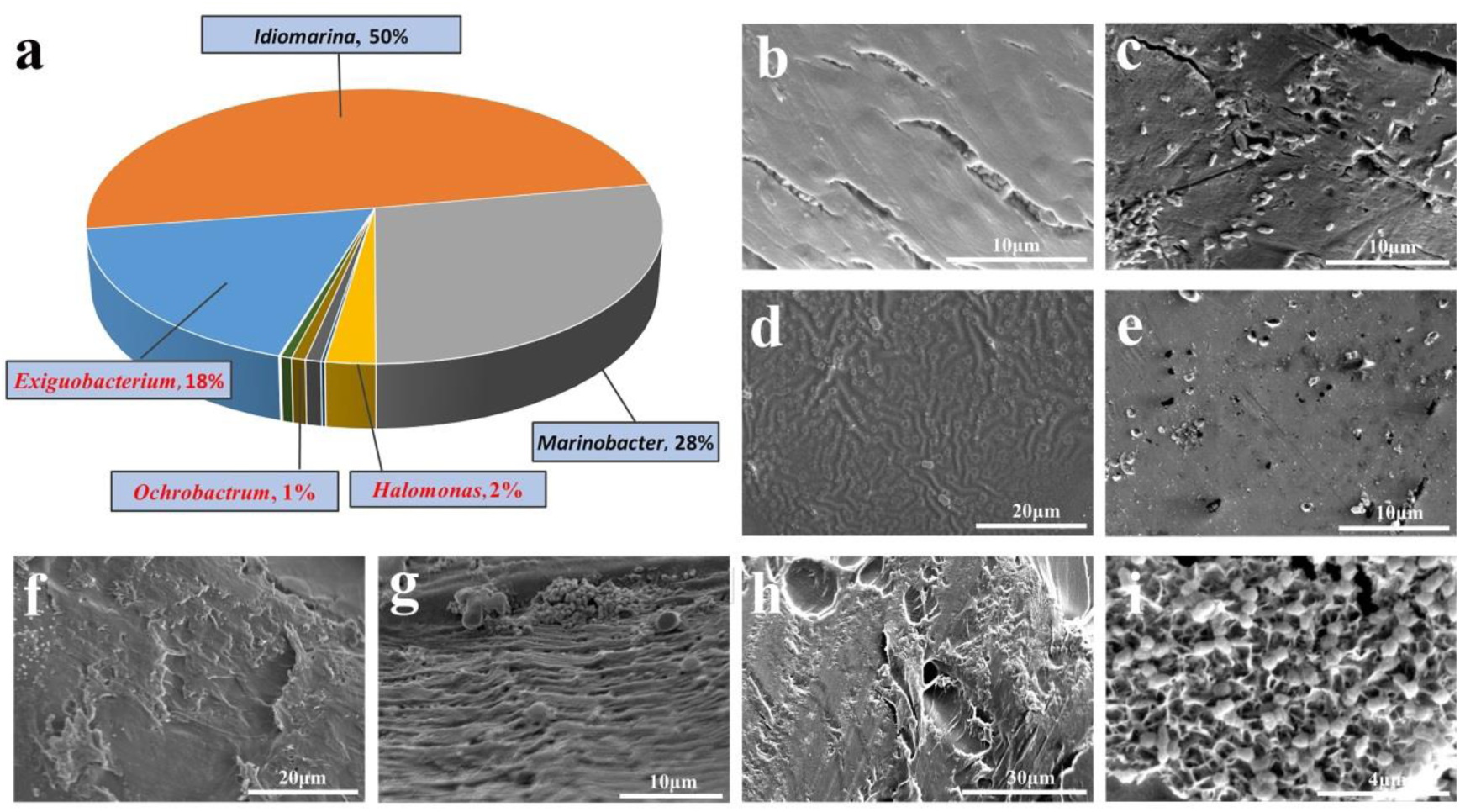
**Abundance quantification and plastic degradation of three core bacteria derived from the original marine bacterial community**. **a**, Absolute abundance quantification of bacteria within the original community with plastic-degrading capability through 16S rRNA sequencing method after 5-day incubation with plastics. The top 5 high abundance general names were shown. The abundance of each genus was indicated after corresponding name. The general names for three core bacteria are highlighted with red color. SEM observation of PET (**b**) and PE (**c**) treated by *Exiguobacterium* sp. for 14 days. SEM observation of PET (**d**) and PE (**e**) treated by *Halomonas* sp. for 14 days. SEM observation of PET (**f**) and PE (**g**) treated by *Ochrobactrum* sp. for 14 days. SEM observation of PET (**h**) and PE (**i**) treated by the mixture of *Exiguobacterium* sp., *Halomonas* sp. and *Ochrobactrum* sp. for 7 days.

**Fig. 3.**
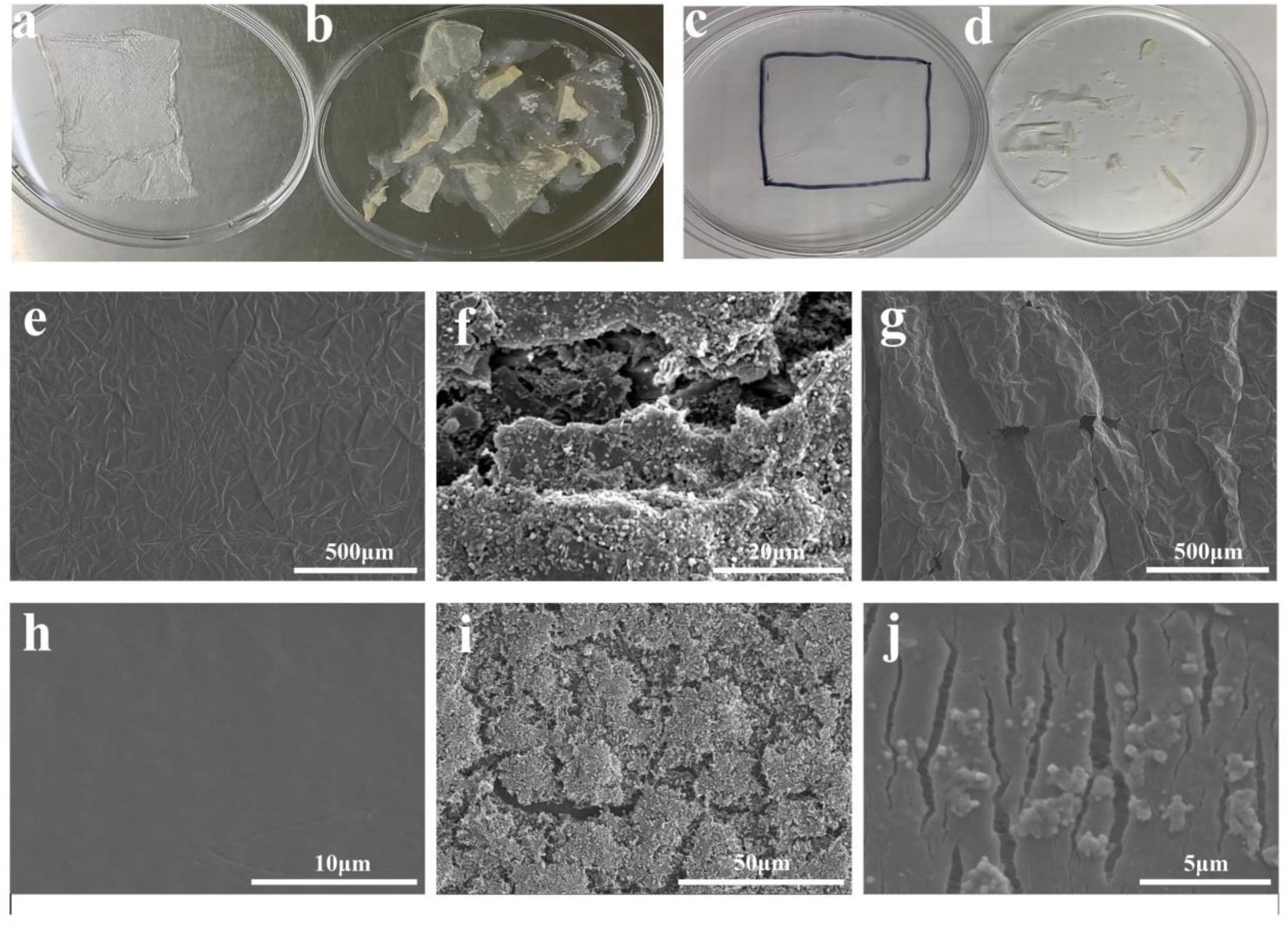
**Observation of the colonization and degradation effects of the reconstituted bacterial community on PET and PE films. a**, Morphology of PET film without treatment. **b**, Morphology of PET film treated by the reconstituted bacterial community for 14 days. **c**, Morphology of PE film without treatment. **d**, Morphology of PE film treated by the reconstituted bacterial community for 14 days. **e**, SEM observation of PET film without treatment. **f**, SEM observation of the colonization of the reconstituted bacterial community on the PET film after 14 days incubation. **g**, SEM observation of the degradation effects of PET film treated by the reconstituted bacterial community for 14 days. **h**, SEM observation of PE film without treatment. **i**, SEM observation of the colonization of the reconstituted bacterial community on the PE film after 14 days incubation. **j**, SEM observation of the degradation effects of PE film treated by the reconstituted bacterial community for 14 days. The type of PET film used for this assay is ES301005 (0.0005 mm in thickness). The type of PE film used for this assay is ET311350 ET311126 (0.025mm in thickness).

### Verification of the degrading effects of the reconstituted bacterial community on PET and PE films

To further verify the degradation effects of the reconstituted bacterial community on PET and PE, we performed multiple techniques to clarify the degradation efficiency and products led by this bacterial community. First, Fourier Transform Infrared (FTIR) imaging was used to analyze the changes of the surface chemical components and function groups of PET and PE treated by this reconstituted bacterial community for four weeks. The FTIR spectra showed that the treated PET had a distinct peak observed at a wave number at 3318 cm^-1^ (Fig. 4a), which was attributed to the carboxylic acid and alcohol functional groups (R-OH stretching, 3000-3500 cm ^-1^)^29^. On the other hand, FTIR spectra of treated PE showed two different peaks compared with the control group (Fig. 4e), one was observed in the vicinity of 1715 cm^-1^ and assigned to the carbonyl band (-C=O-), the other was observed at a wave number at 3318 cm^-1^ and was attributed to the hydroxyl group^15^ (Fig. 4e). Thus, it is reasonable to see that the reconstituted bacterial community has a better degradation efficiency on PE than PET due to the dysfunction of more key bonds of PE. Overall, according to the FTIR spectra, the formation of carboxylic acid end groups and carbonyl bands suggested the hydrolysis reaction of PET and PE, which led to a reduction in molecular weight of the main polymer and cleavage of the ester bond of PET polymer and carbon-carbon bond of PE.

**Fig. 4.**
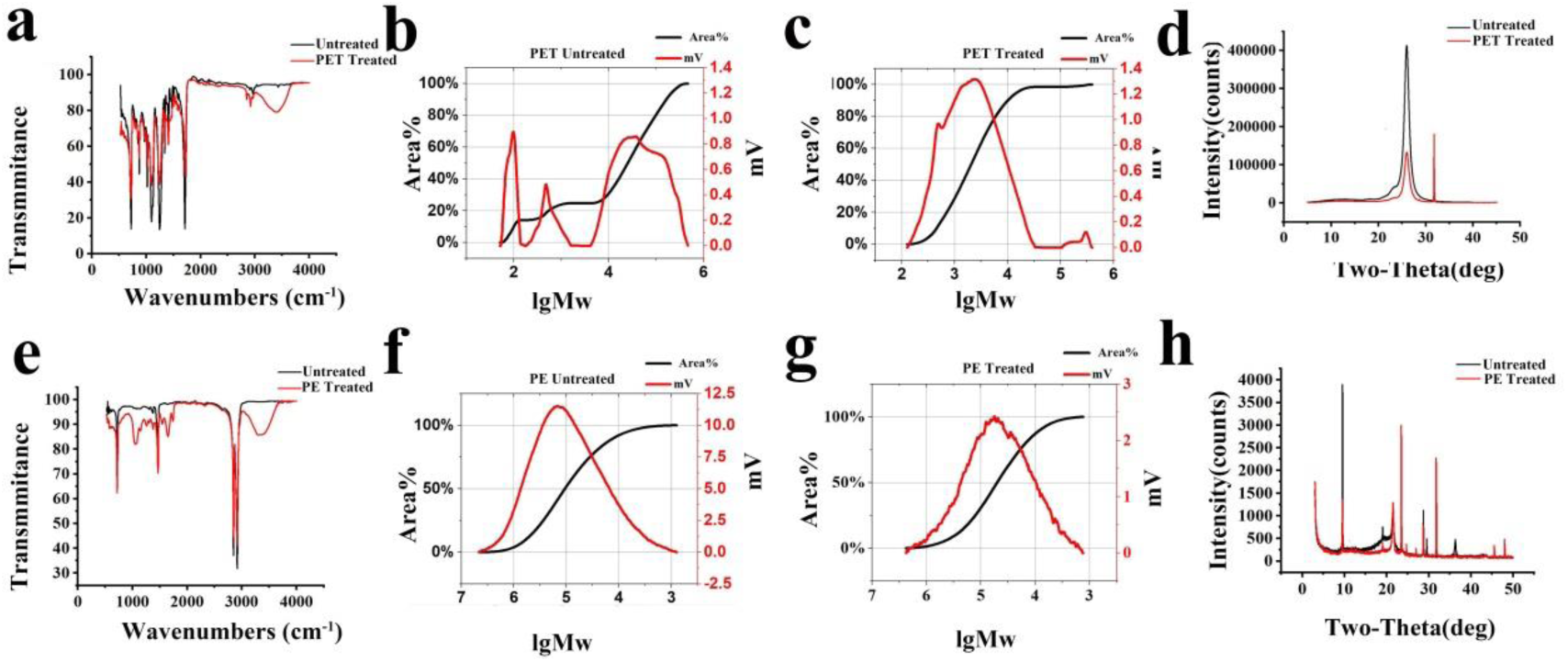
**Validation of PET and PE degradation by the reconstituted bacterial community. a**, FTIR analysis of untreated and treated PET film by the reconstituted bacterial community for 28 days. **b**, GPC analysis of untreated PET film. **c**, GPC analysis of PET film treated by the reconstituted bacterial community for 28 days. **d**, XRD analysis of untreated and treated PET film by the reconstituted bacterial community for 28 days. **e**, FTIR analysis of untreated and treated PE film by the reconstituted bacterial community for 28 days. **f**, GPC analysis of untreated PE film. **g**, GPC analysis of PE film treated by the reconstituted bacterial community for 28 days. **h**, XRD analysis of untreated and treated PET film by the reconstituted bacterial community for 28 days. The type of PET film used for this assay is ES301005 (0.0005 mm in thickness). The type of PE film used for this assay is ET311350 ET311126 (0.025mm in thickness).

Furthermore, the molecular weight distribution (MWD) changes for PET and PE treated by the reconstituted bacterial community for four weeks were analyzed by Gel Permeation Chromatography (GPC). The MWD of the treated PET showed an obvious depolymerization trend (Fig. 4c), and two peaks appeared in the curve, one representing the range of molecular weight of 98,451-399,162 Da (accounting for 1.55%), the other representing the range of molecular weight of 126-33,575 Da (accounting for 98.44%). By comparison, the MWD curve of control had three peaks (Fig. 4b), representing the range of molecular weight of 2,824-472,417 Da (accounting for 75%), 170-1,617 Da (accounting for 10.64%) and 51-140 Da (accounting for 14.17%), respectively. Clearly, the GPC results indicated the treated PET had a significant reduction in the proportion of large molecules, and an obvious increase in the scale of small molecules. Similarly, the MWD of the treated PE decreased from 231,017Da to 122,388Da (Figs. 4f, 4g). Consistently, according to a peak-differentiating and imitating calculation analyzed by X-Ray Diffraction (XRD), the relative value of crystallinity degree reduced from 92.55% to 89.85% for treated PET for four weeks (Fig. 4d), and decreased from 49.10% to 29.50% for treated PE (Fig. 4h) for four weeks. Together, in combination of the results of SEM observation, FTIR, GPC and XRD analyses, we conclude that the reconstituted bacterial community indeed possesses a strong capability of degrading both PET and PE.

Next, the degraded products by the reconstituted bacterial community were analyzed by High-performance liquid chromatography-Mass spectrometry (HPLC-MS). For PET, terephthalic acid (TPA), ethylene glycol (EG), mono-(2-hydroxyethyl) terephthalate (MHET) and bis (2-hydroxyethyl) terephthalate (BHET) have been identified as the main enzymatic degradation products^19^. Consistently, TPA (Figs. 5a, 5b) and MHET (Figs. 5c, 5d) were identified as hydrolysis products after PET was treated by the reconstituted bacterial community for four weeks. The degradation products of PE after a 14-day treatment were also analyzed by HPLC-MS. The results showed significant differences in the abundance and categories of the eluted compounds as compared with the control group (Fig. 5e), especially in the end of the mass spectrum graph, strongly suggesting PE was decomposed to novel substances. Unfortunately, the chemical identity of these degradation compounds was not confirmed due to lack of standard samples but their presence supports the hypothesis of PE degradation by the reconstituted bacterial community.

**Fig. 5.**
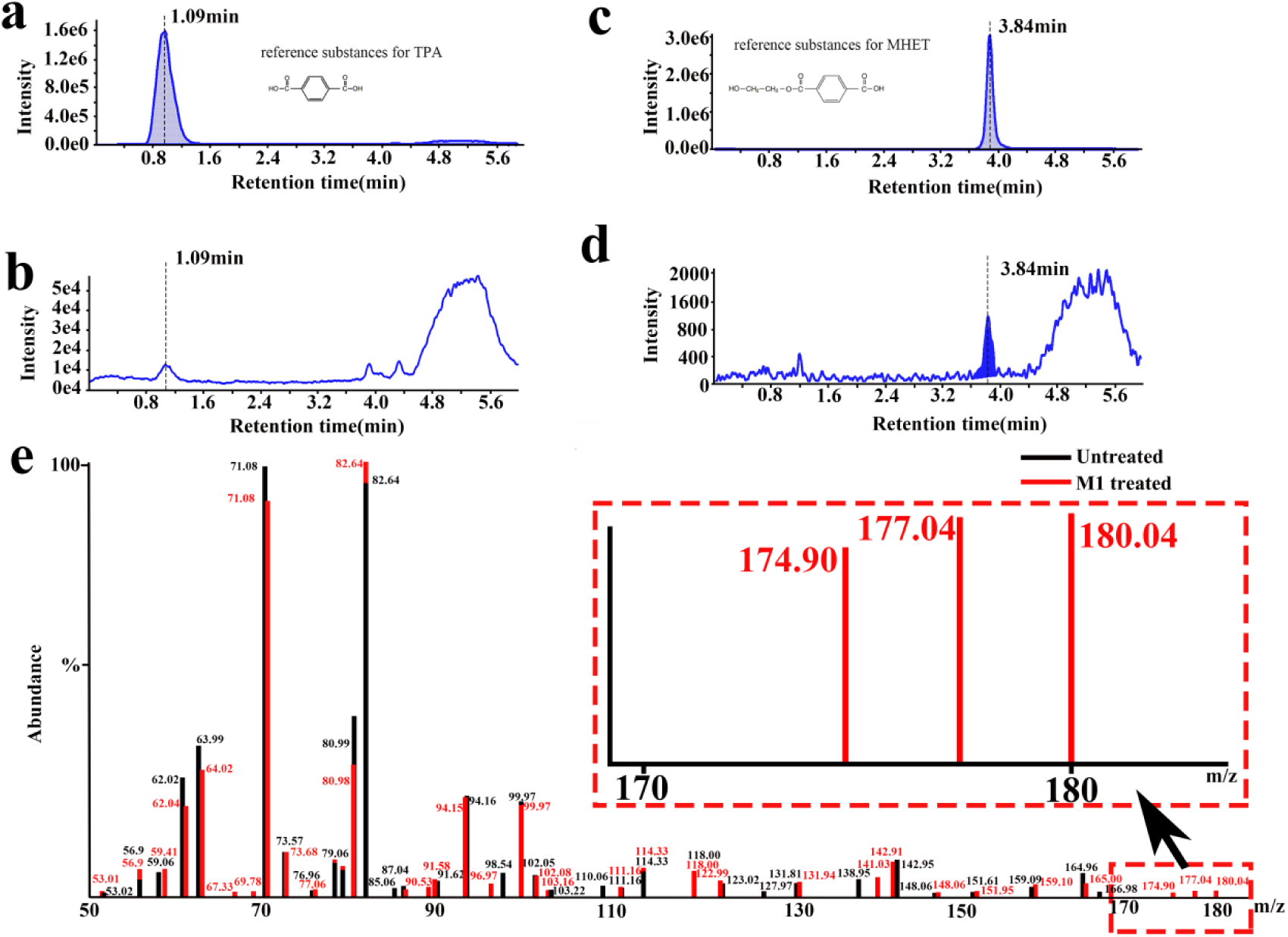
**Analysis of the released products from PET and PE films treated by the reconstituted bacterial community. a**, HPLC spectrum of the standard terephthalic acid (TPA). **b**, HPLC spectrum of the product (prediction as TPA) released from the PET film treated by the reconstituted bacterial community for 28 days. **c**, HPLC spectrum of the standard mono-(2-hydroxyethyl) terephthalate (MHET). **d**, HPLC spectrum of the product (prediction as MHET) released from the PET film treated by the reconstituted bacterial community for 28 days. **e**, HPLC-MS spectrum of the products released from the PE film treated by the reconstituted bacterial community for 14 days. Magnification of the area is indicated by a dashed rectangle. The type of PET film used for this assay is ES301005 (0.0005 mm in thickness). The type of PE film used for this assay is ET311126 (0.025mm in thickness).

### Transcriptomic profiling of the plastic degradation process and mechanism led by the reconstituted bacterial community

To explore the plastic degradation process and potential mechanisms mediated by the reconstituted bacterial community, we performed a macro transcriptome analysis of this flora in the presence of PET or PE. Based on our above results, we chose three time points (8 h, 7 d and 14 d) for further transcriptome analysis. According to the analysis of genes differential expression after 8 h incubation with PET or PE, the significantly up-regulated genes in the three bacteria (*Exiguobacterium* sp., *Halomonas* sp.*, Ochrobactrum* sp.) were mainly associated with energy production and cell growth regardless of the plastic type, such as citrate cycle and ribosomal biosynthesis (Fig. 6a). Clearly, in the first 8 h incubation time, bacteria mainly utilized the easily available nutrient in the original minimal medium to quickly multiply. When extending the incubation time to 7 d or 14 d, most markedly up-regulated genes were closely related to biofilm formation (such as quorum sensing, bacterial chemotaxis, flagellar assembling and two-component system), bacterial secretion system, and cell growth and reproduction (such as citrate cycle, carbon metabolism, fatty acid degradation and ribosomal biosynthesis), regardless of plastic type and incubation time (Fig. 6b). Based on the growth assay of the bacterial community shown in this study, with 7 d- or longer incubation time, the population quantity of the community is still stable in the presence of plastics (Supplementary Fig. 3a), suggesting the bacteria could still obtain enough nutrient to support cell growth. Given very little carbon source provided in the original minimal medium, it is reasonable to propose that bacteria are forced to colonize and thereby degrading and utilizing the plastics-the only available nutrient source in the environment. To better occupy the surface of plastics, the best choice is to form biofilm, which could explain why the expression of large amount genes associated with biofilm formation were significantly up-regulated, and this result also consists well with the observation of huge biofilm formation in the plastic surface (Figs. 1-3). Once the bacterial community successfully colonizes on the targets, they might secret diverse enzymes to degrade the plastics. As shown in Figs. 6c and 6d, the expression of many genes encoding potential plastics degrading enzymes was evidently up-regulated, such as lipase, esterase, cutinase and hydrolase. With this, the bacteria could obtain energy for growth and reproduction through degradation and utilization of plastics, as revealed by the transcriptomic results that the expressions of many genes associated with energy production and carbon metabolism were significantly increased (Fig. 6b).

**Fig. 6.**
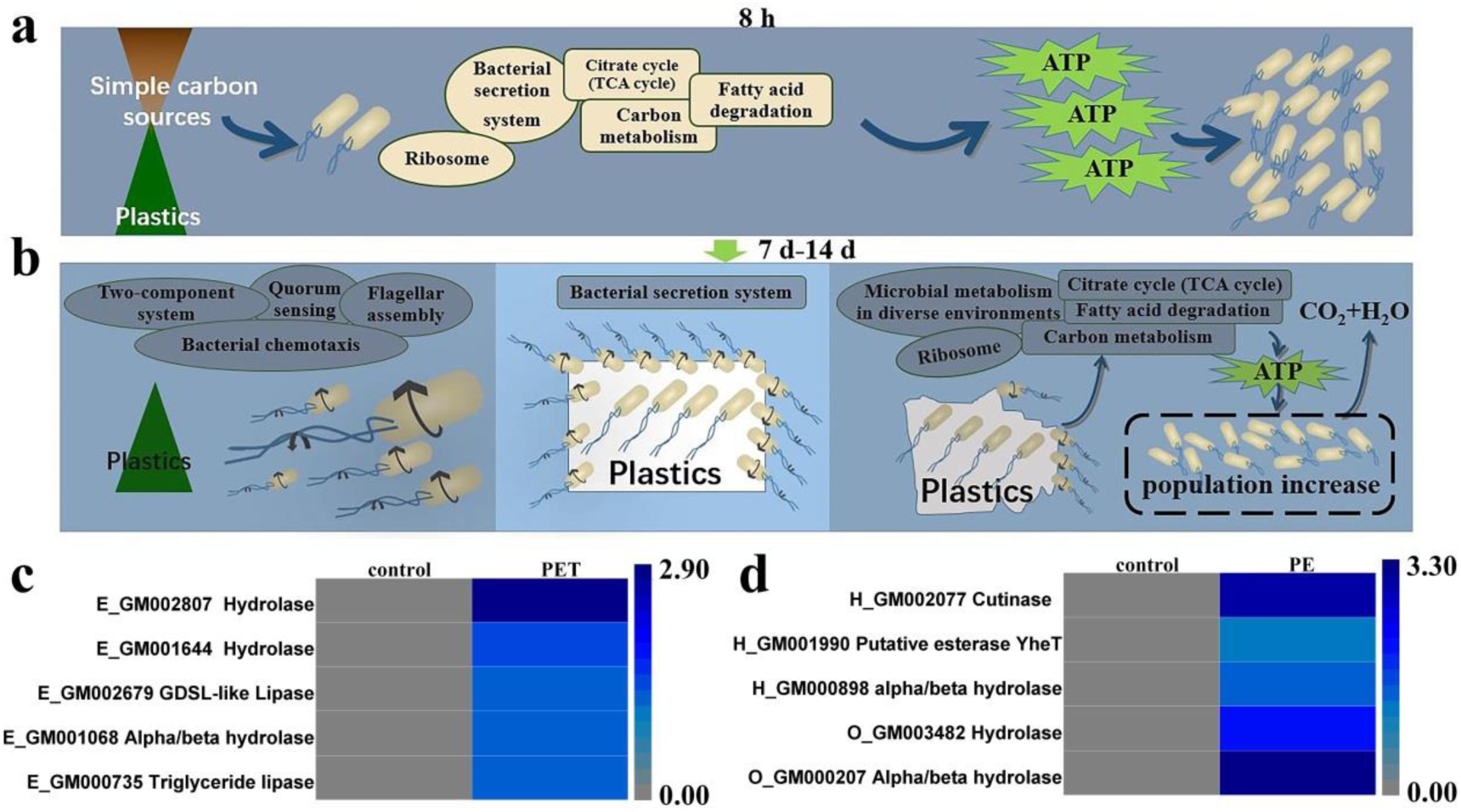
**Transcriptomic analysis of the plastic degradation process and mechanisms mediated by the reconstituted bacterial community. a**, Growth status of the reconstituted bacterial community cultured in the minimal medium for 8 hours supplemented with PET or PE film. **b**, The proposed model of plastic degradation and utilization mediated by the reconstituted bacterial community cultured in the minimal medium for 7 or 14 days supplemented with PET or PE film. The proposed growth and metabolic pathways were referred to the transcriptomic results. The type of PET film used for this assay is ES301005 (0.0005 mm in thickness). The type of PE film used for this assay is ET311350 ET311126 (0.025mm in thickness). **c**, Heat map showing the predicted PET-degradation enzymes derived from the reconstituted bacterial community. **d**, Heat map showing the predicted PE-degradation enzymes derived from the reconstituted bacterial community. All the transcriptomic data associated with this figure are shown in the Supplementary information (Supplemental Figs. 4-63) and uploaded in the public database.

## Discussion

Plastics have become a global concern as the accumulation in the world’s oceans and their impacts on marine organisms and human health^2–4, 30^. Therefore, much effort has been exerted to reduce plastic wastes. To remove plastic wastes and recycle plastic-based materials, biocatalytic degradation might be applied as an ecofriendly method^18, 19^. Microbes have potentials to degrade plastics with ester bond via enzymatic hydrolysis through colonization onto the surfaces of materials^18, 19, 21^. Because some marine bacteria have the ability to degrade hydrocarbons which possessing similar chemical structure with plastics, it has been suggested that a certain fraction of the microbial community colonizing plastics might have capability of degrading plastics^31, 32^ and thereby using as a carbon matrix. However, it remains unclear whether and to what extent marine prokaryotic communities are capable of degrading plastic in the ocean. For the first time, this study investigated the community structures of marine microbial biofilms attached to two different plastics (PET and PE) (Fig. 2 and Supplementary Table 1), successfully reconstituted a tailored bacterial community possessing significant plastic degrading capabilities (Figs. 3, 4), and clarified the degradation process and products eventually (Figs. 5, 6). Therefore, our results answer the question that whether oceanic bacteria are capable to degrade plastics, and clearly show that plastic waste associated bacteria in the marine environment have great potentials to develop plastic degradation bio-products.

Indeed, we obtained the functional bacterial community efficiently degrading plastics from a bay heavily contaminated by plastic waste, while we failed to find any potential plastic-degrading bacterial flora from the deep-sea sediment samples, indicating that it is advisable to screen potential degraders in the plastic polluted locations in the future. Similarly, several researchers have obtained microplastic-associated bacterial community having plastic-degrading potentials in the field investigation. For example, the plastisphere bacterial communities on PET surfaces in the North Sea^33^, bacterial biofilms associated with PE, PP and glass during an *in-situ* incubation experiment taking place in the northern Adriatic Sea^27^. However, these studies mainly focus on the community structures and dynamics of bacteria on the surfaces of plastics, the metabolic pathways and associated mechanistic processes involved in the biodegradation of plastics are yet to be characterized. In the present study, we disclosed the degradation details through transcriptomics methods, revealing three potential steps including biofilm formation, degradation and utilization involved in the degradation process exerting by bacterial esterase, cutinase and hydrolase (Fig. 6). Overall, our study paves a way to obtain plastic degraders in marine environments and a good candidate to explore plastic degradation mechanisms in the future.

Of note, in the present study, we adopted community but not individual bacterium for plastic degradation given the better degradation effect of three bacteria over single bacterium against plastic waste (Fig. 2). Indeed, the efficiency for biodegradation and bioremediation of environmental pollutants using single strains is still very low and restricted^28^. In contrast, mixed flora has stronger environmental adaptability, higher degradation efficiency and more ample scope for the greater use of biotechnologies in biodegradation compared with single pure bacterium due to the synergistic effect of different microorganisms among them^28^. Therefore, more and more attention has been shifted towards microbial consortia, since their inherent multiple function, robust and adaptable characteristics. Actually, great achievements have obtained in the bioremediation fields with microbial consortia, such as treatment of wastewater eutrophication^34^, removal of heavy metals^35^, degradation of dyes^36^. By comparison, there is rarely report on plastic degradation led by bacterial community, especially the reconstituted functional consortium. Our study pioneered a way to advance the development of high-efficient, stable and controllable synthetic microbial consortia against different plastic waste. Notably, three bacteria within the reconstituted bacterial community belong to the genera of *Exiguobacterium*, *Halomonas* and *Ochrobactrum*, which are all easily cultured and fast grown bacteria, providing a great advantage for the future application.

In addition, the present bacterial community prefers to act on PE and enables to degrade the intact PE film to debris (Fig. 3d), showing a great capability to solve the PE degradation problem. It is known that PE is largely utilized in packaging, representing ∼40% of total demand for plastic products (www.plasticseurope.org) with over a trillion plastic bags used every year^15^. PE comprises a linear backbone of carbon atoms and holds the highly stable carbon-carbon (C-C) bonds, which is resistant to degradation and has become a big challenge for plastic waste biodegradation. PE biodegradation has been observed with an extreme long incubation time (up to couples of months), given appropriate conditions. For example, modest degradation of PE was observed after a combination of nitric acid treatment and 3-month incubation with the fungus *Penicillium simplicissimum*^37^. Even slower PE degradation was also recorded after 4 to 7 months exposure to the bacterium *Nocardia asteroids*^38^. Excitedly, our reconstituted bacterial community could lead a significant degradation towards PE (e.g. ubiquitous cracks in the surface and damages in the four corners of PE film) within several days, and totally degrade the PE film to very small pieces within 2 weeks. Undoubtedly, given the fast rate of biodegradation reported here, these findings have potential for significant biotechnological applications. Further investigation is also required to explore the detailed nature of the products derived from degraded PE exerted by this reconstituted bacterial community, and determine what enzymes responsible for PE/PET degradation predicted in this study.

## Methods

### Screening of microbial community degrading PET and PE

About 300 sediment samples contaminated by plastic debris from different locations of Huiquan Bay (Qingdao, China) were collected and kept in the plastics container until using for screening. Three kinds of PET plastic were used for degradation assays in the present study, including plastic drink bottle, type ES301450 (0.25 mm in thickness) and type ES301005 (0.0005 mm in thickness), the latter two were purchased from the Good Fellow Company (UK). Similarly, three kinds of PE plastic were also used in the present study, including commercial PE bags, type ET311350 (0.25mm in thickness) and type ET311126 (0.025mm in thickness), the latter two were also purchased from the Good Fellow Company (UK). None additives were contained in the films purchased from the Good Fellow Company according to the manufacturer’s standard (Q/SH3180014). All PET and PE films used in this study were treated in 75% ethanol, and then air-dried in a laminar-flow clean bench prior to use. For screening microbial community degrading plastics, PET or PE films were all cut into 30 mm × 20 mm square sheets and incubated in minimal medium (0.5 g yeast extract, 1 g peptone, in 1 L filtered sea water, and PH is adjusted to 7.0.) containing different plastics-contaminated samples for couples of weeks to months before checking the degradation effects.

### Absolute quantification of individual bacterial abundance in the community

The original bacterial community cultivated in the minimal medium without the supplement of any plastics was set as a control group. Correspondingly, the culture supplement with PET (type ES301450) or PE (type ET311350) was set as experimental groups. The samples was collected in five different growth stages (group A: OD_600_=0.3, cultivated for 4.5 h; group B: OD_600_=0.58, cultivated for 7.5 h, group C: OD_600_=0.85, cultivated for 8.5 h; group D: OD_600_=1.05, cultivated for 9.5 h; group E: OD_600_=1.13, cultivated for 7 d). Three parallel groups were set for absolute quantification of 16S rRNA sequences. The analyses were performed by Genesky Biotechnologies Inc. (Shanghai, China). Briefly, total genomic DNAs of different samples were extracted by using the QIAamp Rapid DNA Kit (Qiagen, Germany). The DNA concentration and integrity were measured by a NanoDrop2000 spectrophotometer (Thermo Fisher Scientific, USA) and agarose gel electrophoresis, respectively. The spike-in sequences contained conserved regions identical to those of natural 16S rRNA genes and artificial variable regions were distinct from those found in nucleotide sequences in public databases. These served as internal standards and facilitated absolute quantification across samples. Appropriate mixtures with known copy numbers of spike-in sequences were added to the sample DNAs. The V4-V5 regions of the 16S rRNA genes and spike-in sequences were amplified and sequenced using Illumina HiSeq.

### Scanning Electron Microscopy (SEM) observation

PET and PE films were collected from each sample and observed through SEM to examine the degradation effects by different bacterial community. After treated by bacteria, the PET or PE films were soaked in 5% glutaraldehyde for cell fixation, then were dehydrated in 30-100% graded ethanol for 15 min each and critical-point-dried with CO_2_^39^. Dried specimens were sputter coated for 5 min with gold and platinum (10 nm) using a Hitachi MC1000 Ion Sputter (Japan), and were examined using a field emission scanning electron microscope (Hitachi S-3400N) operating at an accelerating voltage of 5 kV. The PET and PE films treated with or without bacterial community were washed in ultrasonic cleaner with 1% SDS, distilled water, and then ethanol^19^. The film was air-dried, coated with gold and platinum (10 nm) using a Hitachi MC1000 Ion Sputter, and subjected to SEM observation.

### Fourier Transform Infrared (FTIR) analysis

The PET and PE plastics exposed to the microbial consortia were recovered after an incubation period of four weeks, then were carefully rinsed in ultrasonic cleaner with 1% SDS, distilled water, and then ethanol^29, 40^. After air dried, the plastic films treated with or without bacterial community were recorded over the wavelength range 450-4000 cm^−1^ at a resolution of 1 cm^-1^ using a Nicolet-360 FTIR (Waltham, USA) spectrometer operating in ATR mode^37, 38^. Thirty two scans were taken for each spectrum.

### Gel Permeation Chromatography (GPC) Analysis

The molecular weight of PET films treated with or without bacterial community was determined by GPC, which were carried out on an instrument of model Shimadzu GPC-20A (Japan) equipped with LC20 columns and operating at 35 °C^41^. Trichlorobenzene was used as mobile phase (1 mL/min) after calibration with polystyrene standards of known molecular mass. A sample concentration of 1 mg/mL was employed^42^. The molecular weight of the PE films treated with or without bacterial community was determined by GPC on an Agilent PL-GPC220 equipped with Agilent PLgel Olexis 300 × 7.5 mm columns and operating at 150 °C^43, 44^. Trichlorobenzene was used as mobile phase (1 mL/min) after calibration with polystyrene standards of known molecular mass. A sample concentration of 1 mg/mL was employed^38, 45^.

### X-Ray Diffraction (XRD) analysis

XRD was carried out by using the Bruker D8 Advance instrument with a wavelength of 1.5406 angstrom of CuKα ray. The XRD tube current was set as 40 mA, and the tube voltage was set as 40 kV. The measurements for PET were set in the angle range from 2θ = 5° to 2θ = 45° at a rate of 1°/min^46, 47^. The measurements for PE were set in the angle range from 2θ = 3° to 2θ = 50° at a rate of 1°/min^48^.

### High-performance liquid chromatography-Mass spectrometry (HPLC-MS) analysis

HPLC-MS for PET was performed on an API QTRAP 5500 LCMS system equipped with a XB-C18 100A analytical column (2.1× 250 mm, 2.6 μm). The mobile phase was methanol at a flow rate of 0.5 mL/min, and the effluent was monitored at a wavelength of 240 nm. The typical elution condition was followed as: 0 to 1 min, 10% (v/v) methanol; 1 to 2 min, 10-50% methanol linear gradient; 2 to 2.5 min, 50-95% (v/v) methanol; 4.5 to 4.6 min, 95-10% methanol linear gradient. The reaction mixture supernatant was diluted with the mobile phase toward to the calibration range, acidified with concentrated HCl (37%) and centrifuged to remove any precipitation^19, 49^. Standard curves of TPA and BHET were prepared in a concentration range from 0.17 to 1 mM and showed no significant difference.

HPLC-MS samples for PE biodegradation were submerged in acetonitrile and sonicated for around 1 minute. The soluble products were then dissolved in 1 mL fresh acetonitrile, which was then transferred to a microcentrifuge tube and spun down for 2 minutes. Then, 2 µL of supernatant was used for LC-MS (Waters ACQUITY SQD, USA) analysis. The degradation products were detected by LC-MS equipped with a ACQUITY UPLC BEH C18 analytical column (2.1 × 50 mm, 1.7 µm) at 50-1500 m/z^15^. The column temperature was 30 °C, and the flow rate was 0.300 mL/min during operation. The typical elution condition was set as: 0 to 3 min, 90% water, 10% acetonitrile; 3 to 4 min, 10-90% acetonitrile linear gradient; 4 to 8 min, 90% (v/v) acetonitrile; 8 to 8.5min, 90-10% acetonitrile linear gradient; 8.5 to 10 min, 10% (v/v) acetonitrile^50^. The difference between the traces, untreated and treated is the MS peak observed at 0.581 minutes.

### Isolation and genome sequencing of three bacteria leading plastics degradation

To isolate the bacteria in the community, the biofilm attached to the films were collected and plated on the 2216E solid medium (containing 5 g/L peptone, 1 g/L yeast extract, 1 L filtered seawater, 15 g agar, pH adjusted to 7.4-7.6). The visible colony was further purified several times until it was considered to be axenic. The purity of bacterial strains was confirmed by repeated partial sequencing of the 16S rRNA gene. Using this method, five pure cultures were obtained and three of them could degrade plastic, which were *Exiguobacterium* sp., *Halomonas* sp., and *Ochrobactrum* sp, which were further incubated and performed genomic sequencing as described previously^51^. Sequencing libraries were generated using NEBNext® Ultra™ DNA Library Prep Kit for Illumina (NEB, USA) following manufacturer’s recommendations and index codes were added to attribute sequences to each sample^52^. Seven databases were used to predict gene functions, including GO^53^, KEGG^54^, COG^55^, NR^56^ and Swiss-Prot^57^.

### Transcriptomics analysis

The reconstituted bacterial community cultivated in the minimal medium without supplement of any plastics was set as a control group. Correspondingly, the culture supplement with PET (type ES301450) or PE (type ET311350) was set as experimental groups. With this, we collected the control or experimental samples grown for 8 h, 7 d, 14 d, respectively. Transcriptomics analysis was performed by Novogene (Tianjin, China). Briefly, total RNAs of all samples were extracted using TRIzol reagent (Invitrogen, USA) and the DNA contamination was ruled out with MEGA clear Kit (Life technologies, USA). Sequencing libraries were generated using NEBNext^®^ Ultra™ Directional RNA Library Prep Kit for Illumina^®^ (NEB, USA) following manufacturer’s recommendations and index codes were added to attribute sequences to each sample. rRNA is removed using a specialized kit that leaves the mRNA. Fragmentation was carried out using divalent cations under elevated temperature in NEBNext First Strand Synthesis Reaction Buffer (5×). The clustering of the index-coded samples was performed on a cBot Cluster Generation System using TruSeq PE Cluster Kit v3-cBot-HS (Illumia) according to the manufacturer’s instructions^58^. After cluster generation, the library preparations were sequenced on an Illumina Hiseq platform and paired-end reads were generated. Raw data of fastq format were firstly processed through in-house perl scripts. In this step, clean data were obtained by removing reads containing adapter, reads containing ploy-N and low quality reads from raw data. At the same time, Q20, Q30 and GC content the clean data were calculated. All the downstream analyses were based on the clean data with high quality. Differential expression analysis of two conditions/groups was performed using the DESeq R package (1.18.0). Then, KOBAS software was used to test the statistical enrichment of differential expression genes in KEGG pathways. GO enrichment analysis of differentially expressed genes was implemented by the GOseq R package, in which gene length bias was corrected. GO terms with corrected *P* value less than 0.05 were considered significantly enriched by differential expressed genes^59, 60^.

### Data availability

The complete genome sequences of *Halomonas* sp., *Exiguobacterium* sp. and *Ochrobactrum* sp. have been deposited at GenBank under the accession numbers PRJNA673668, PRJNA673665 and PRJNA673670, respectively.

## Acknowledgements

This work is funded by the China Ocean Mineral Resources R&D Association Grant (Grant No. DY135-B2-14), National Key R and D Program of China (Grant No. 2018YFC0310800), Strategic Priority Research Program of the Chinese Academy of Sciences (Grant No. XDA22050301), the Taishan Young Scholar Program of Shandong Province (tsqn20161051), and Qingdao Innovation Leadership Program (Grant No. 18-1-2-7-zhc) for Chaomin Sun.

## Author Contributions

RG and CS conceived and designed the study. RG performed all the experiments. RG and CS analyzed the data. RG wrote the manuscript. CS revised the manuscript. All authors read and approved the final manuscript.

## Supplemental Figures

**Supplementary Fig. 1.**
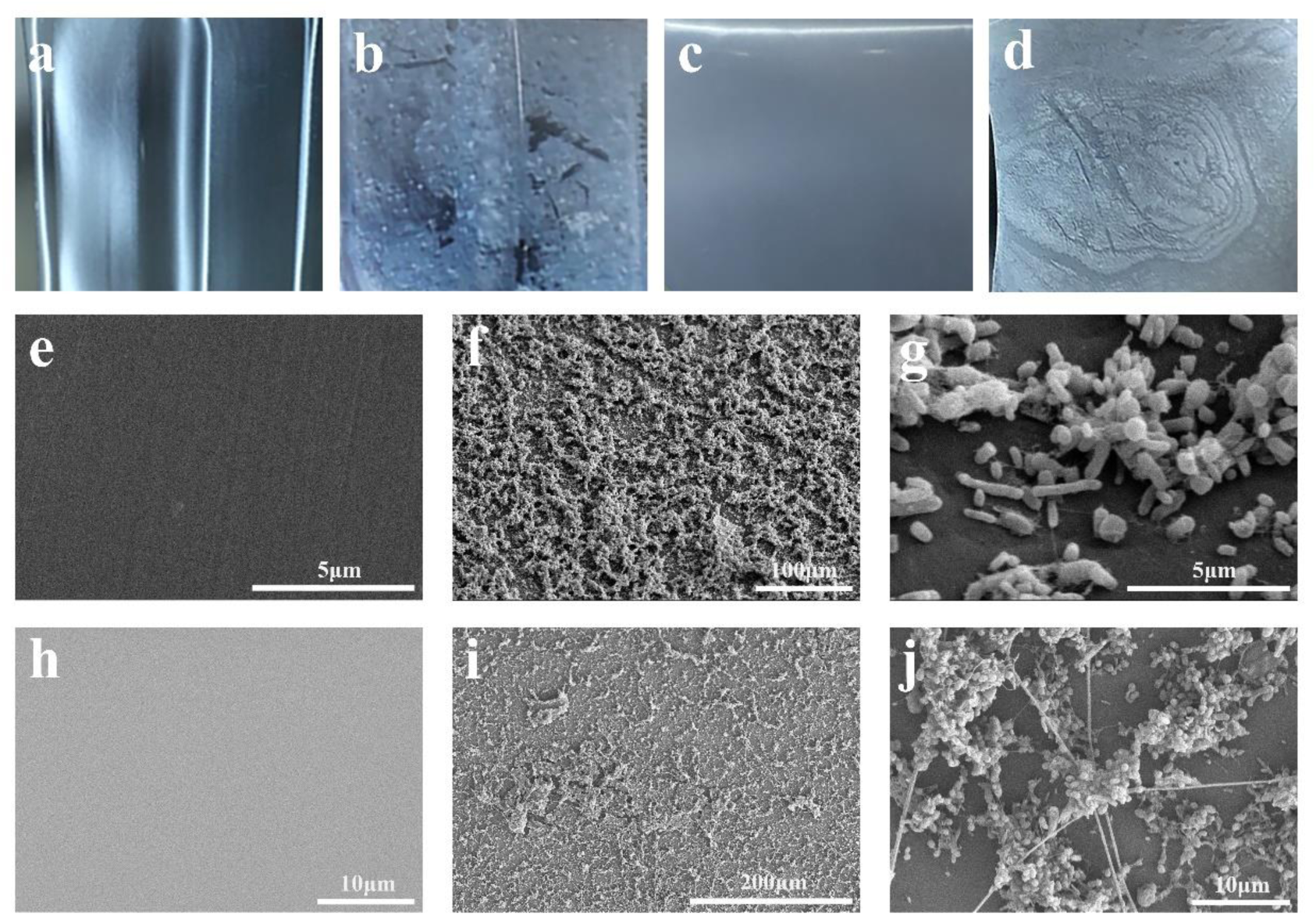
**Observation of the colonization and degradation effects of a marine bacterial community on PET and PE films**. **a**, Morphology of PET film without treatment. **b**, Morphology of PET film treated by the marine bacterial community for seven days. **c**, Morphology of PE film without treatment. **d**, Morphology of PE film treated by the marine bacterial community for seven days. **e**, SEM observation of PET film without treatment. **f**, **g**, SEM observation of the colonization of the bacterial community on the PET film after seven days incubation. **h**, SEM observation of PE film without treatment. **i**, **j**, SEM observation of the colonization of the bacterial community on the PE film after seven days incubation. The type of PET film used for this assay is derived from the drink bottle containing additives. The type of PE film used for this assay is derived from the commercial PE bags containing additives.

**Supplementary Fig. 2.**
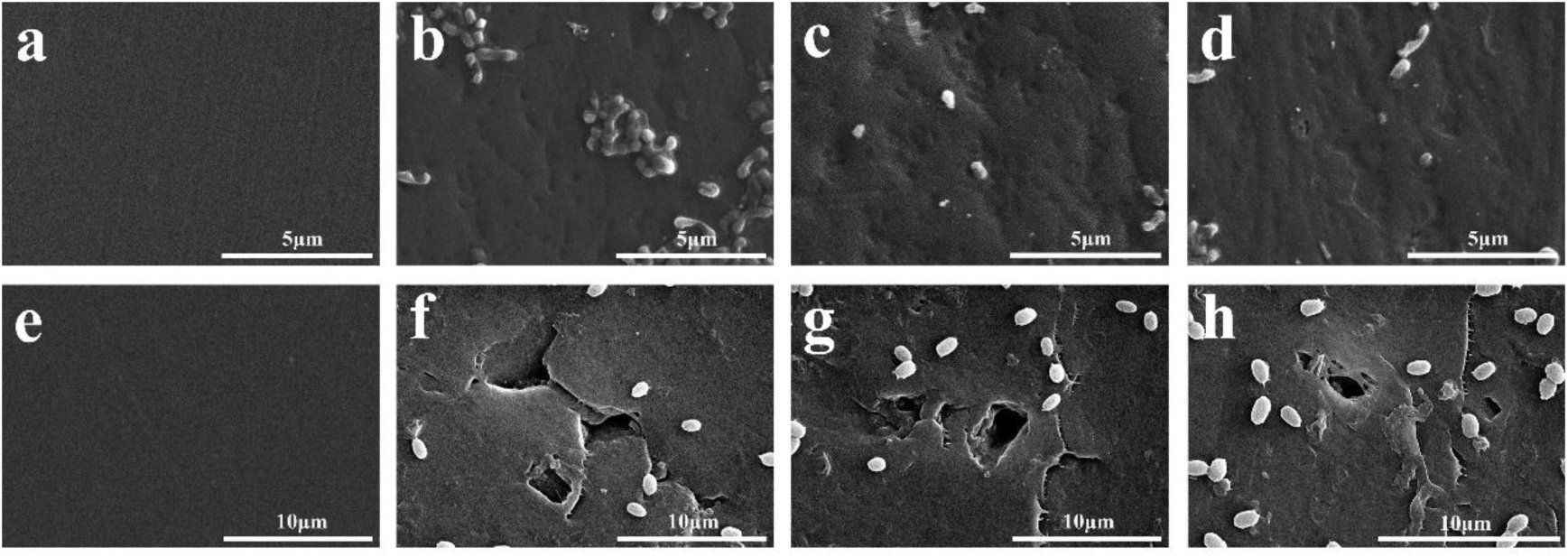
**SEM observation of degradation effects on PET and PE films by the marine bacterial community**. **a**, SEM observation of PET film without treatment. **b-d**, SEM observation of the degradation effects on the PET film by the marine bacterial community after seven days incubation. **e**, SEM observation of PE film without treatment. **f-h**, SEM observation of the degradation effects on the PE film by the marine bacterial community after seven days incubation. The type of PET film used for this assay is derived from the drink bottle containing additives. The type of PE film used for this assay is derived from the commercial PE bags containing additives.

**Supplementary Fig. 3.**
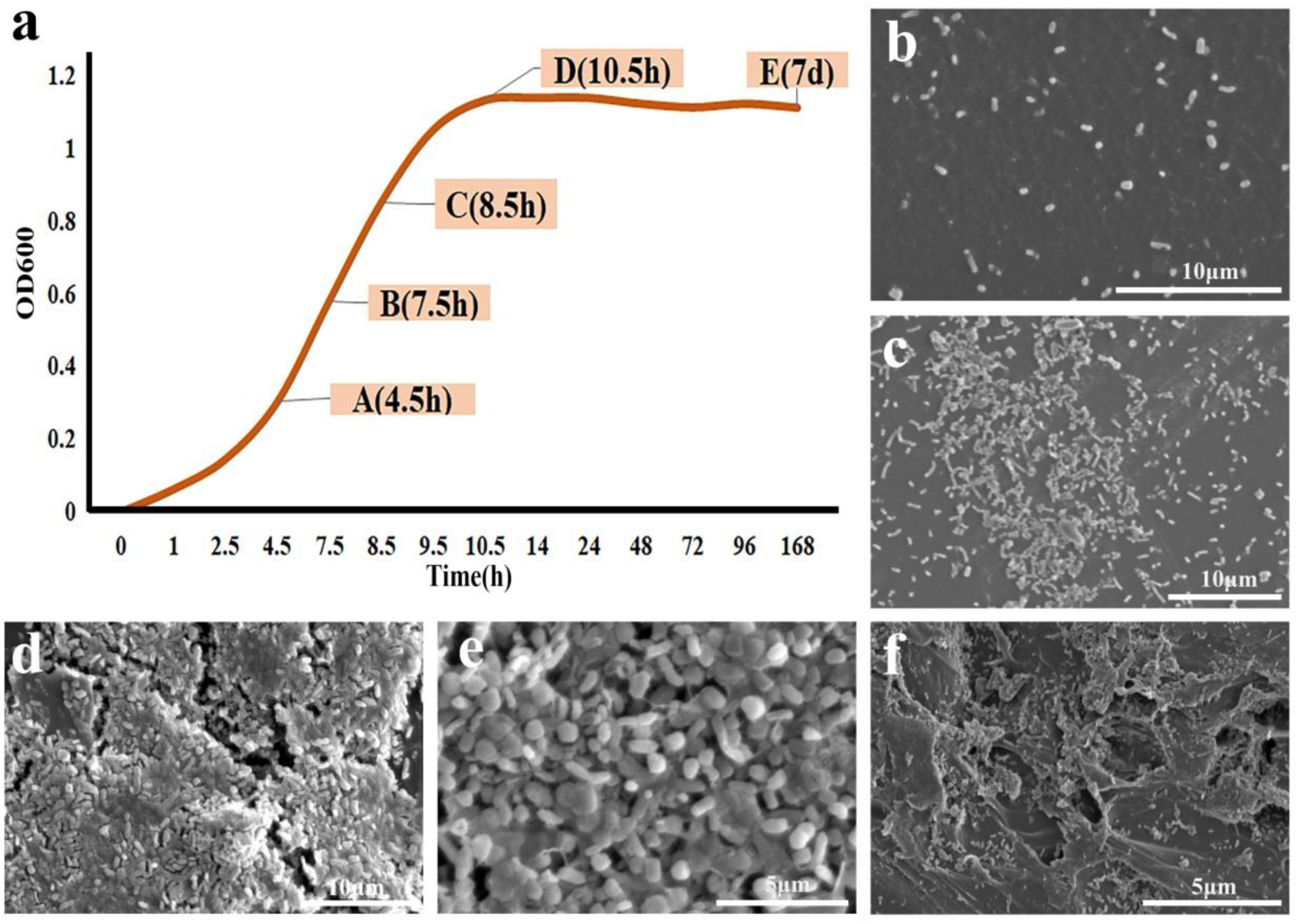
**Growth assay, colonization and degradation effects of the marine bacterial community towards PET and PE films. a**, Growth assay of the marine bacterial community in the presence of PET and PE films in five stages including OD_600_=0.3 (cultivated for 4.5 h), OD_600_=0.58 (cultivated for 7.5 h), OD_600_=0.85 (cultivated for 8.5 h), OD_600_=1.05 (cultivated for 9.5 h), OD_600_=1.13 (cultivated for 7 d). **b**, SEM observation of the colonization of the marine bacterial community on the plastic surface after cultivation for 4.5 h. **c**, SEM observation of the colonization of the marine bacterial community on the plastic surface after cultivation for 7.5 h. **d**, SEM observation of the colonization of the marine bacterial community on the plastic surface after cultivation for 8.5 h. **e**, SEM observation of the colonization of the marine bacterial community on the plastic surface after cultivation for 10.5 h. **f**, SEM observation of the degradation effects of the marine bacterial community on the plastic after cultivation 7 d. The type of PET film used for this assay is ES301450 (0.25 mm in thickness). The type of PE film used for this assay is ET311350 (0.25mm in thickness).

**Supplementary Fig. 5.**
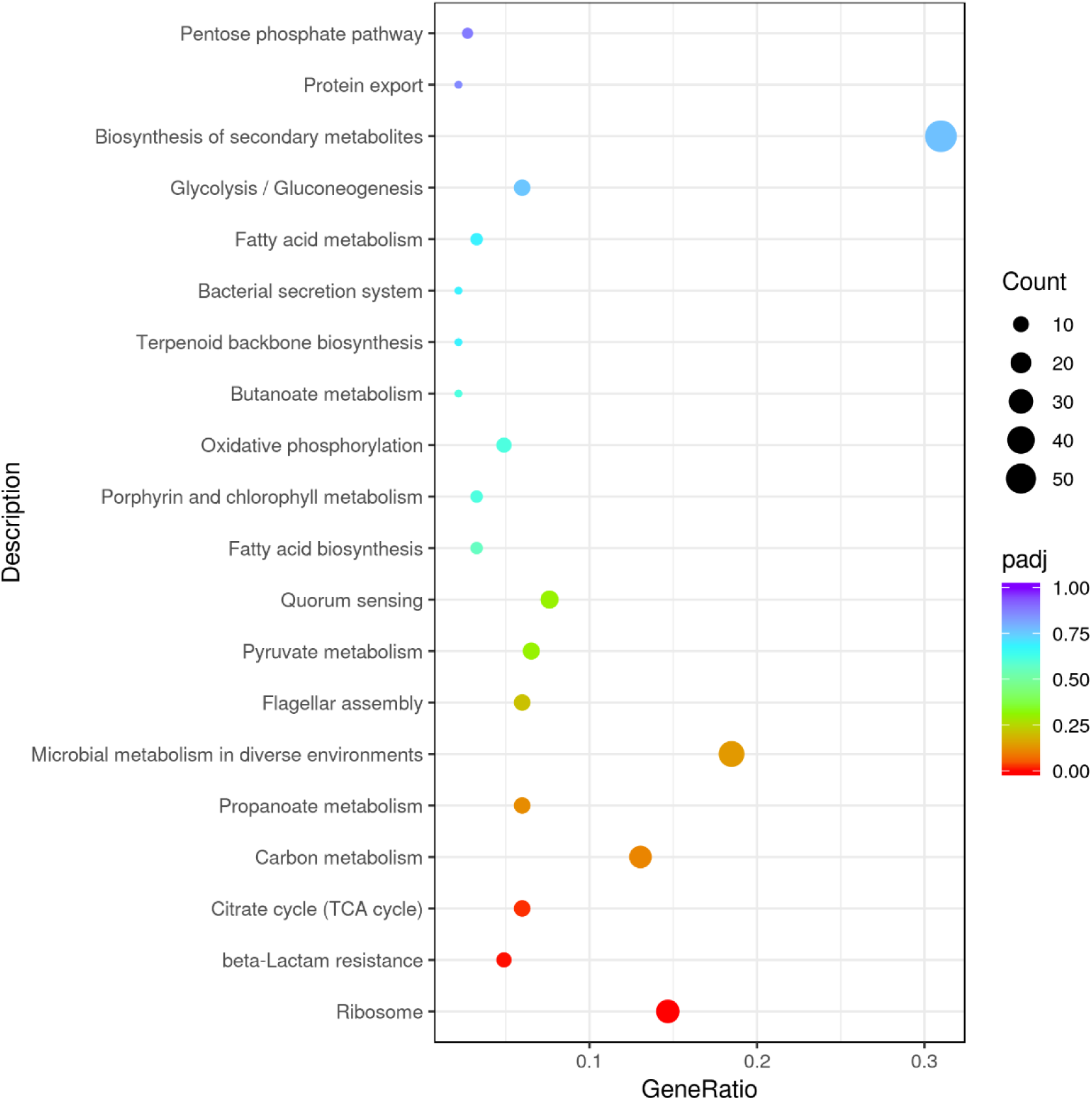
**Up-regulation KEGG pathways enrichment (scatter plot) based on the transcriptomic analysis of PET degradation by *Exiguobacterium* sp. for 8 h.**

**Supplementary Fig. 6.**
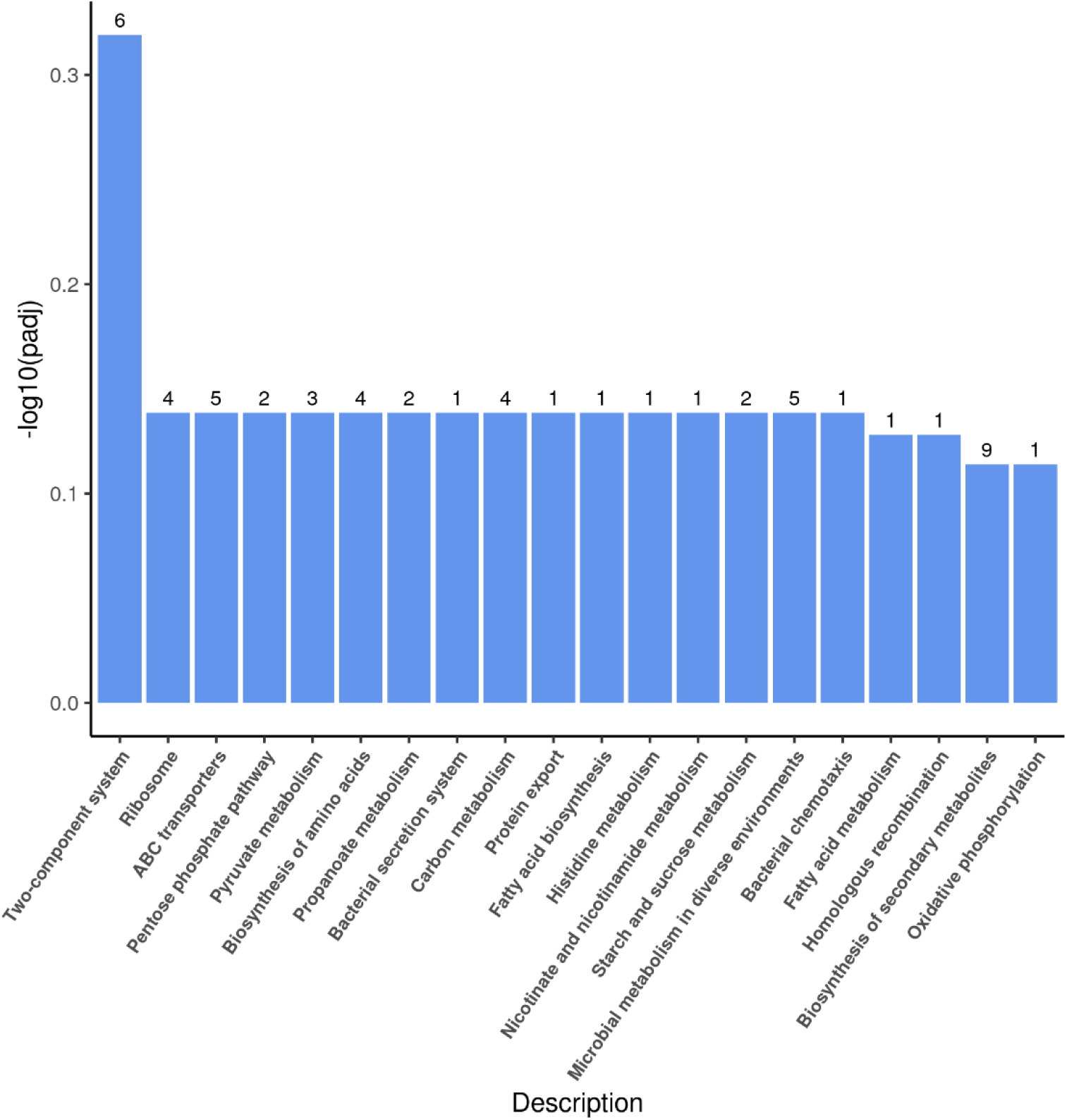
**Up-regulation KEGG pathways enrichment (histogram) based on the transcriptomic analysis of PET degradation by *Exiguobacterium* sp. for 7 d.** The numbers above the column are corresponding genes number related to different pathways.

**Supplementary Fig. 7.**
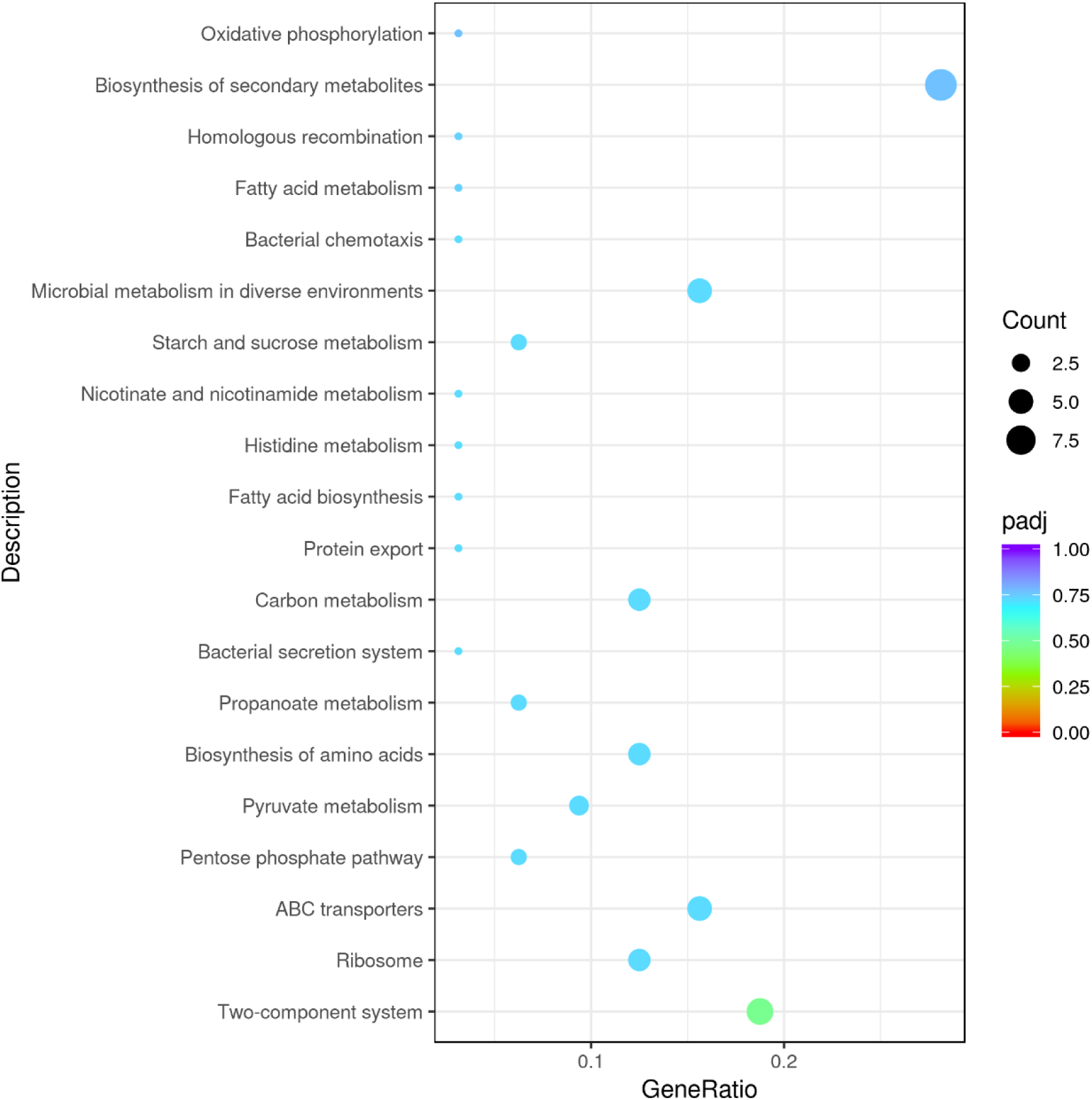
**Up-regulation KEGG pathways enrichment (scatter plot) based on the transcriptomic analysis of PET degradation by *Exiguobacterium* sp. for 7 d.**

**Supplementary Fig. 8.**
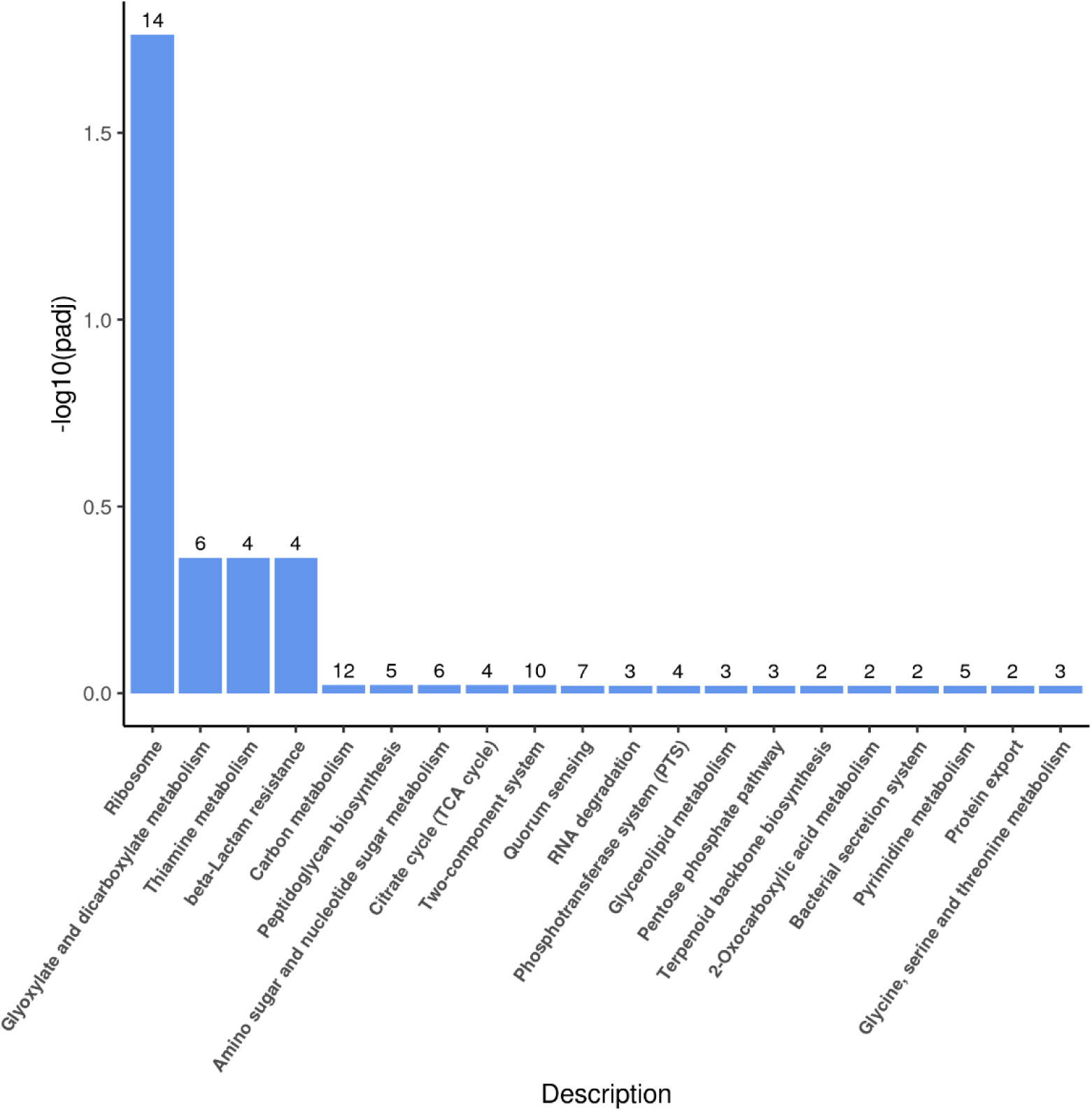
**Up-regulation KEGG pathways enrichment (histogram) based on the transcriptomic analysis of PET degradation by *Exiguobacterium* sp. for 14 d.** The numbers above the column are corresponding genes number related to different pathways.

**Supplementary Fig. 9.**
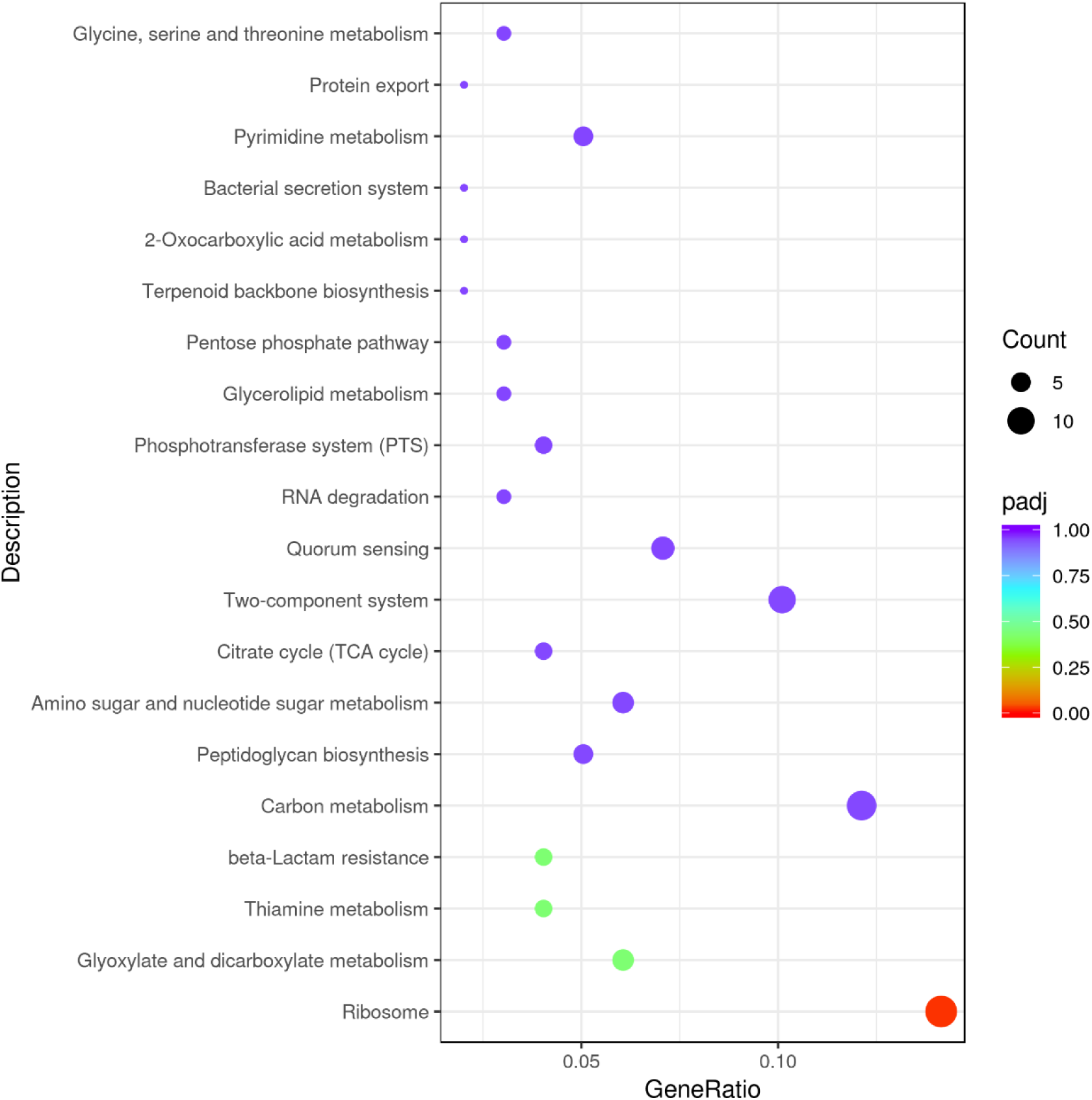
**Up-regulation KEGG pathways enrichment (scatter plot) based on the transcriptomic analysis of PET degradation by *Exiguobacterium* sp. for 14 d.**

**Supplementary Fig. 10.**
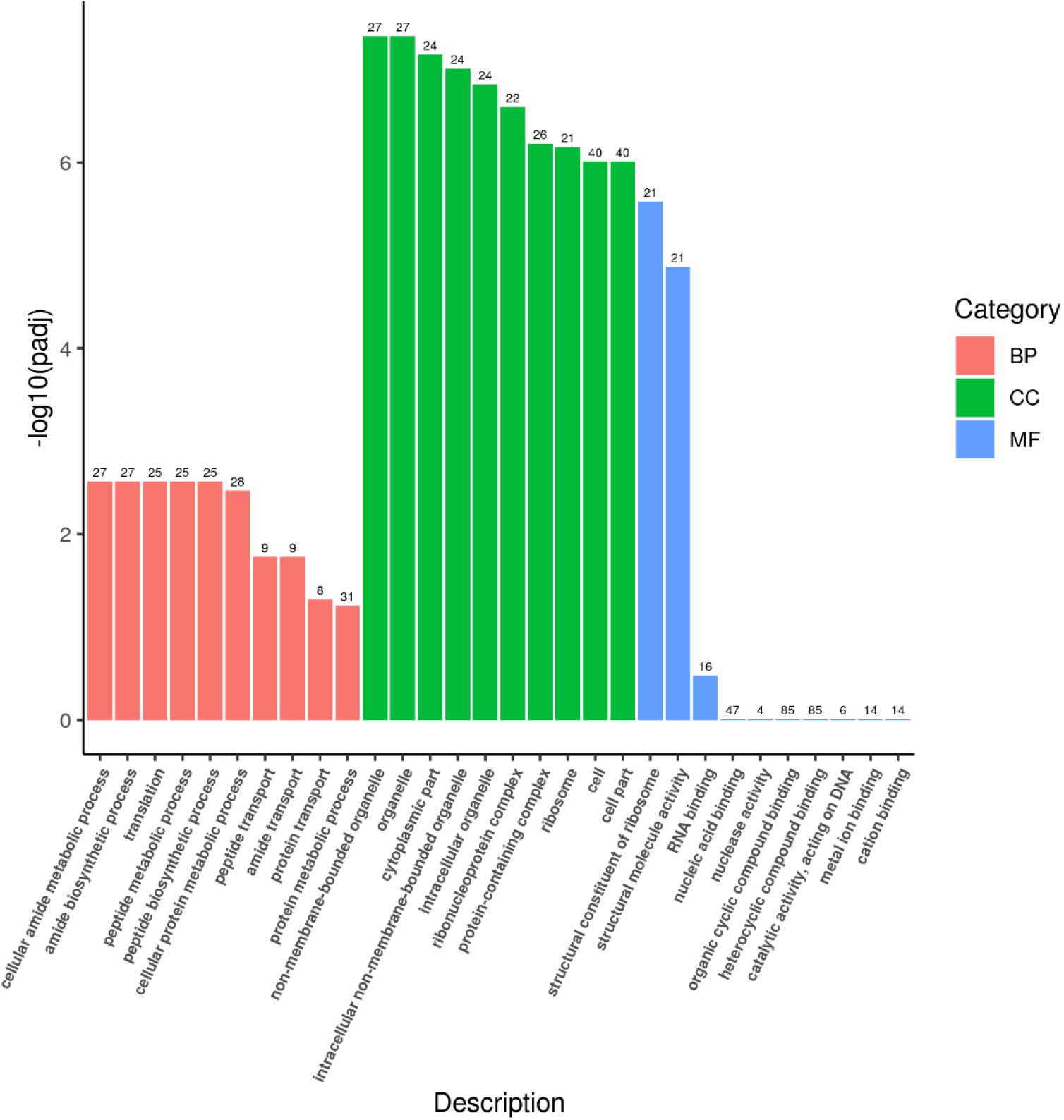
**Up-regulation Go enrichment (histogram) based on the transcriptomic analysis of PET degradation by *Exiguobacterium* sp. for 8 h.** The numbers above the column are corresponding genes number related to different pathways.

**Supplementary Fig. 11.**
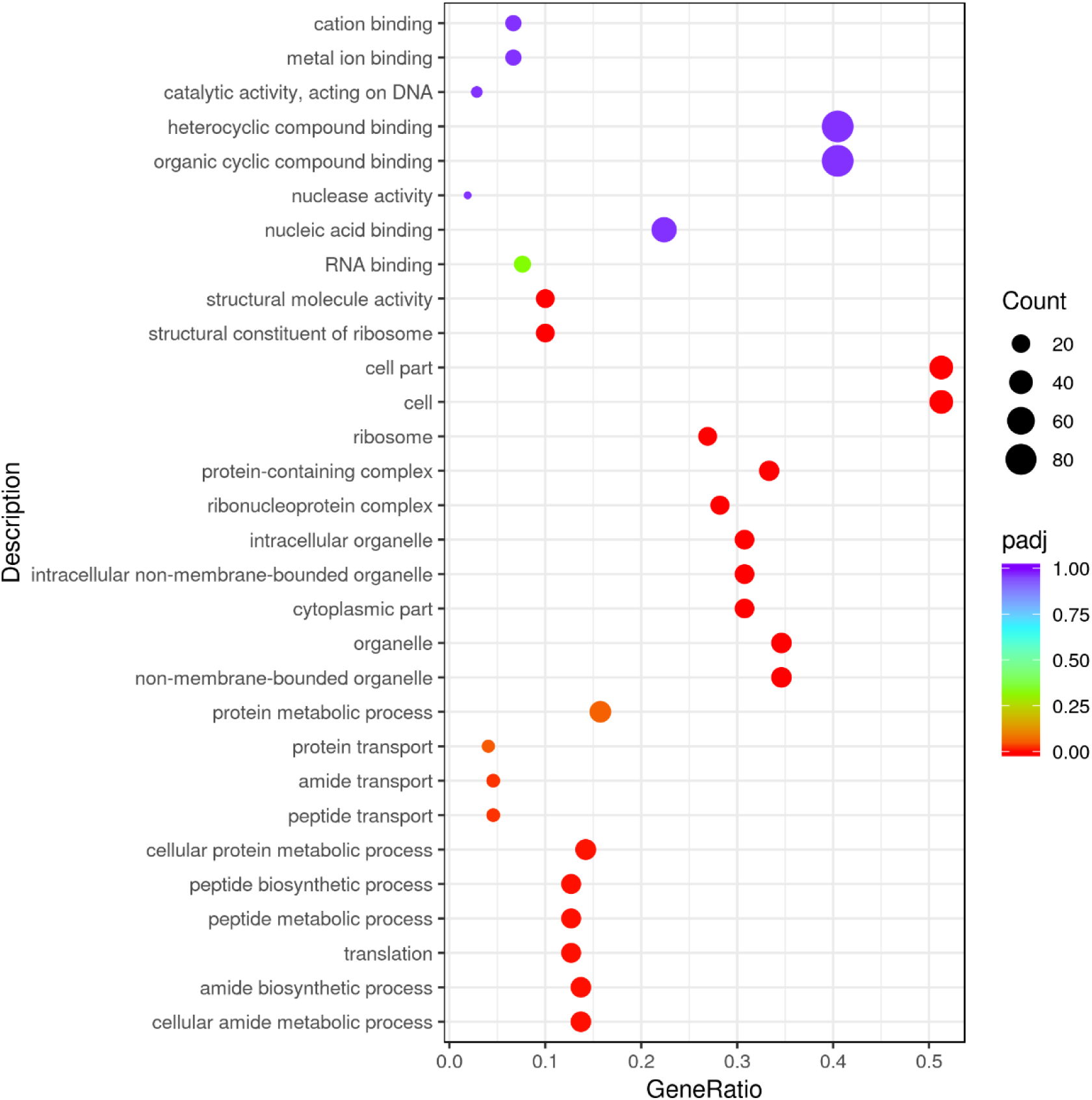
**Up-regulation GO enrichment (scatter plot) based on the transcriptomic analysis of PET degradation by *Exiguobacterium* sp. for 8 h.**

**Supplementary Fig. 12.**
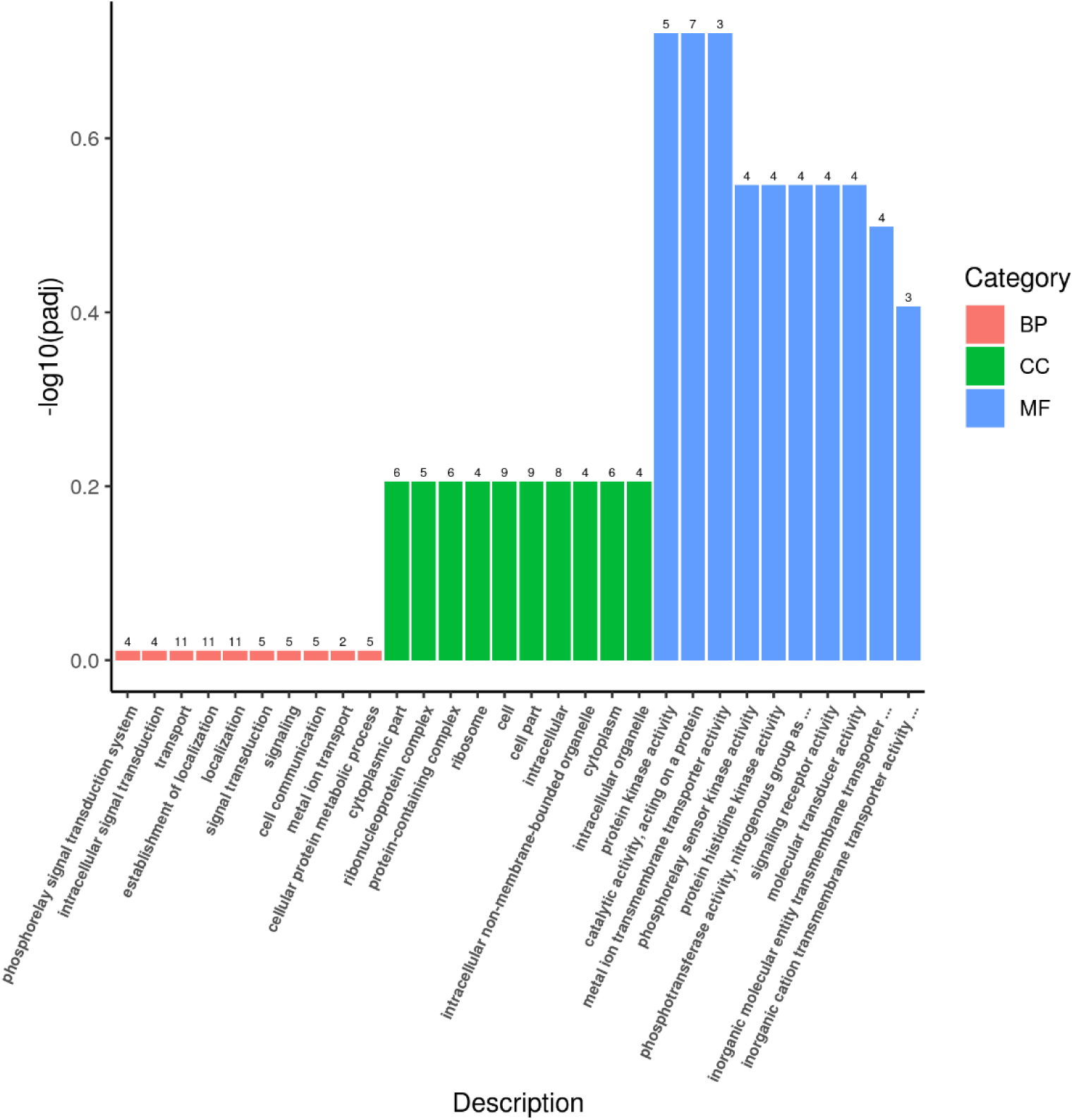
**Up-regulation Go enrichment (histogram) based on the transcriptomic analysis of PET degradation by *Exiguobacterium* sp. for 7 d.** The numbers above the column are corresponding genes number related to different pathways.

**Supplementary Fig. 13.**
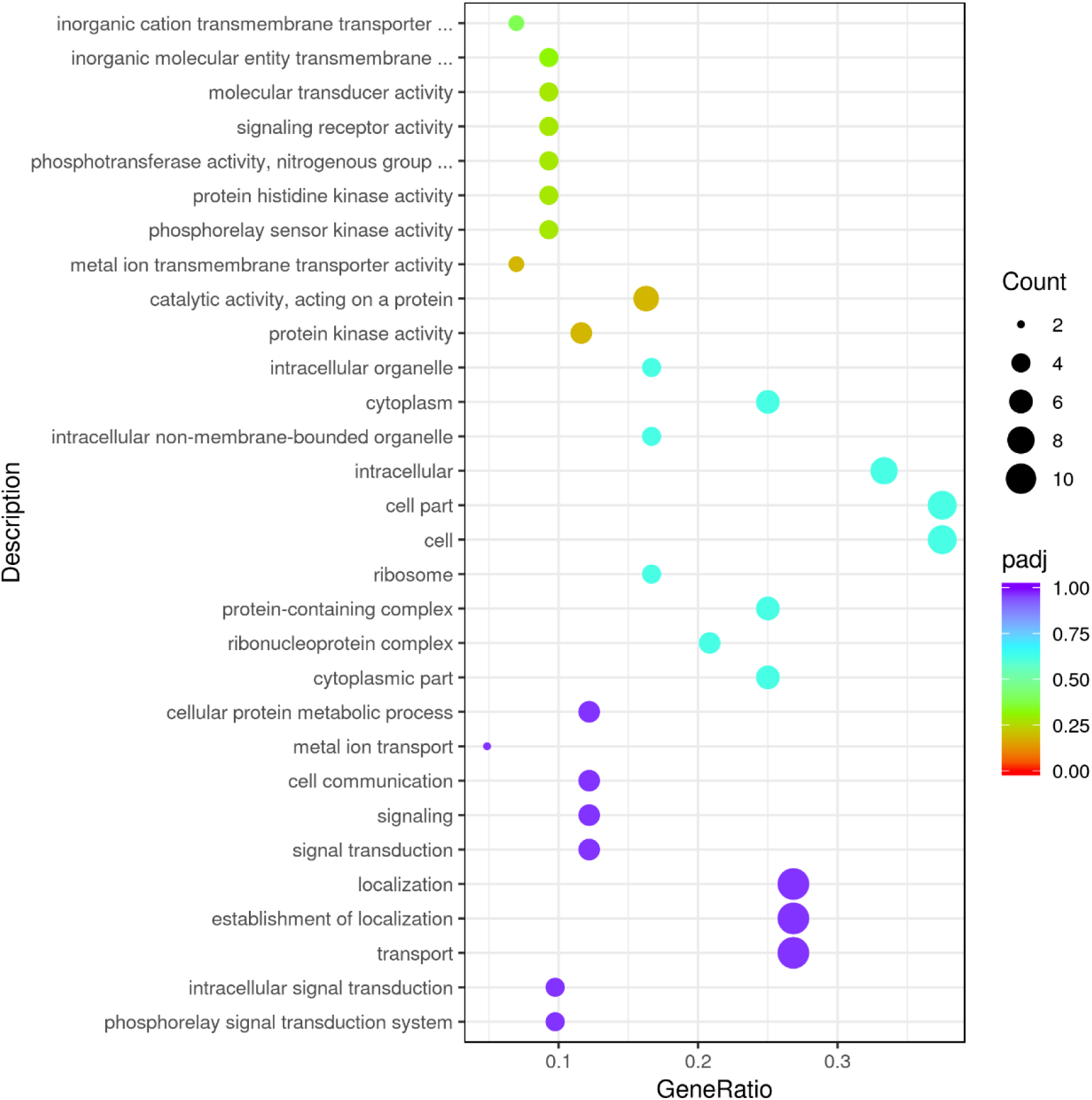
**Up-regulation GO enrichment (scatter plot) based on the transcriptomic analysis of PET degradation by *Exiguobacterium* sp. for 7 d.**

**Supplementary Fig. 14.**
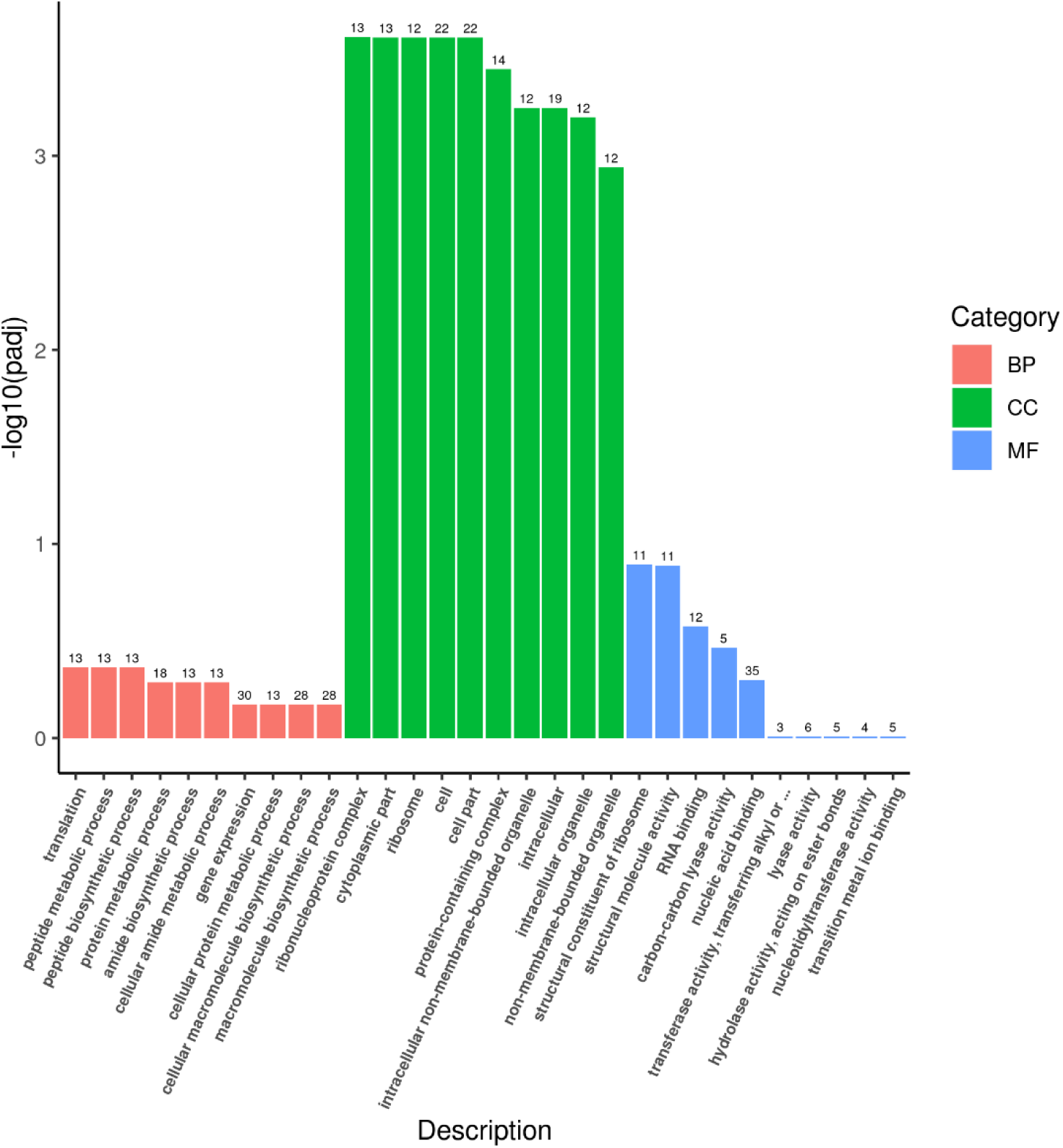
**Up-regulation Go enrichment (histogram) based on the transcriptomic analysis of PET degradation by *Exiguobacterium* sp. for 14 d.** The numbers above the column are corresponding genes number related to different pathways.

**Supplementary Fig. 15.**
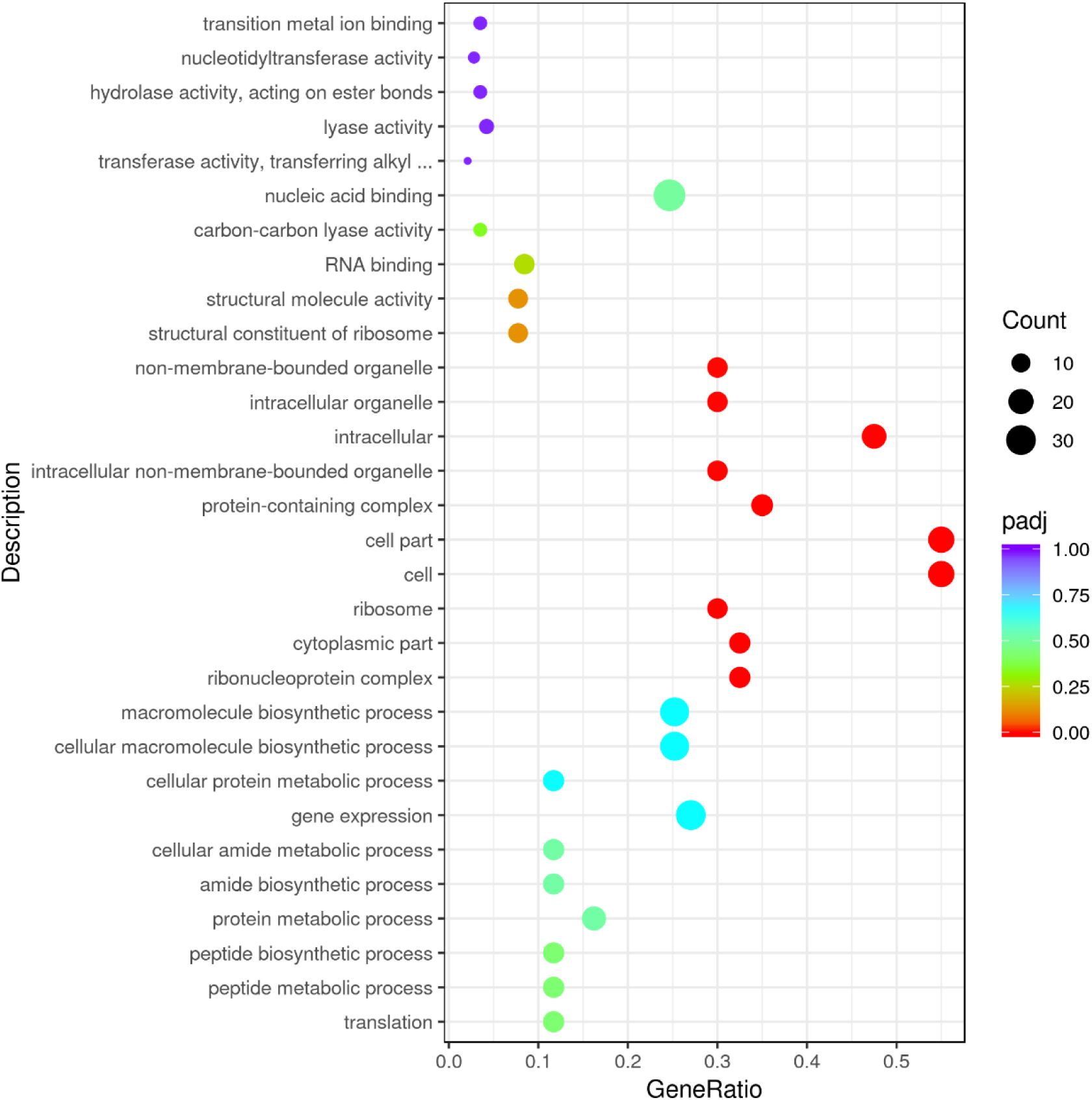
**Up-regulation GO enrichment (scatter plot) based on the transcriptomic analysis of PET degradation by *Exiguobacterium* sp. for 14 d.**

**Supplementary Fig. 16.**
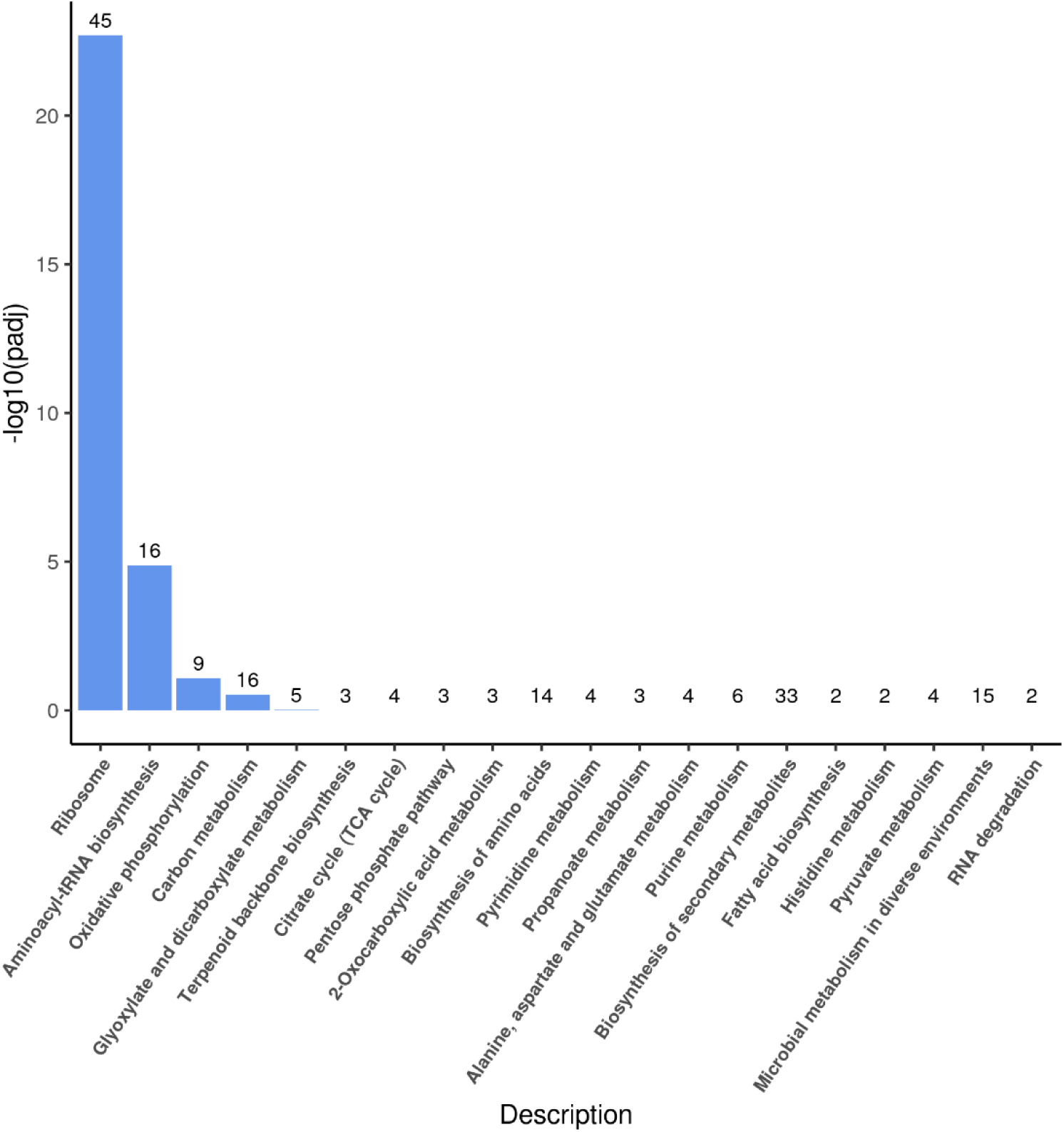
**Up-regulation KEGG pathways enrichment (histogram) based on the transcriptomic analysis of PET degradation by *Halomonas* sp. for 8 h.** The numbers above the column are corresponding genes number related to different pathways.

**Supplementary Fig. 17.**
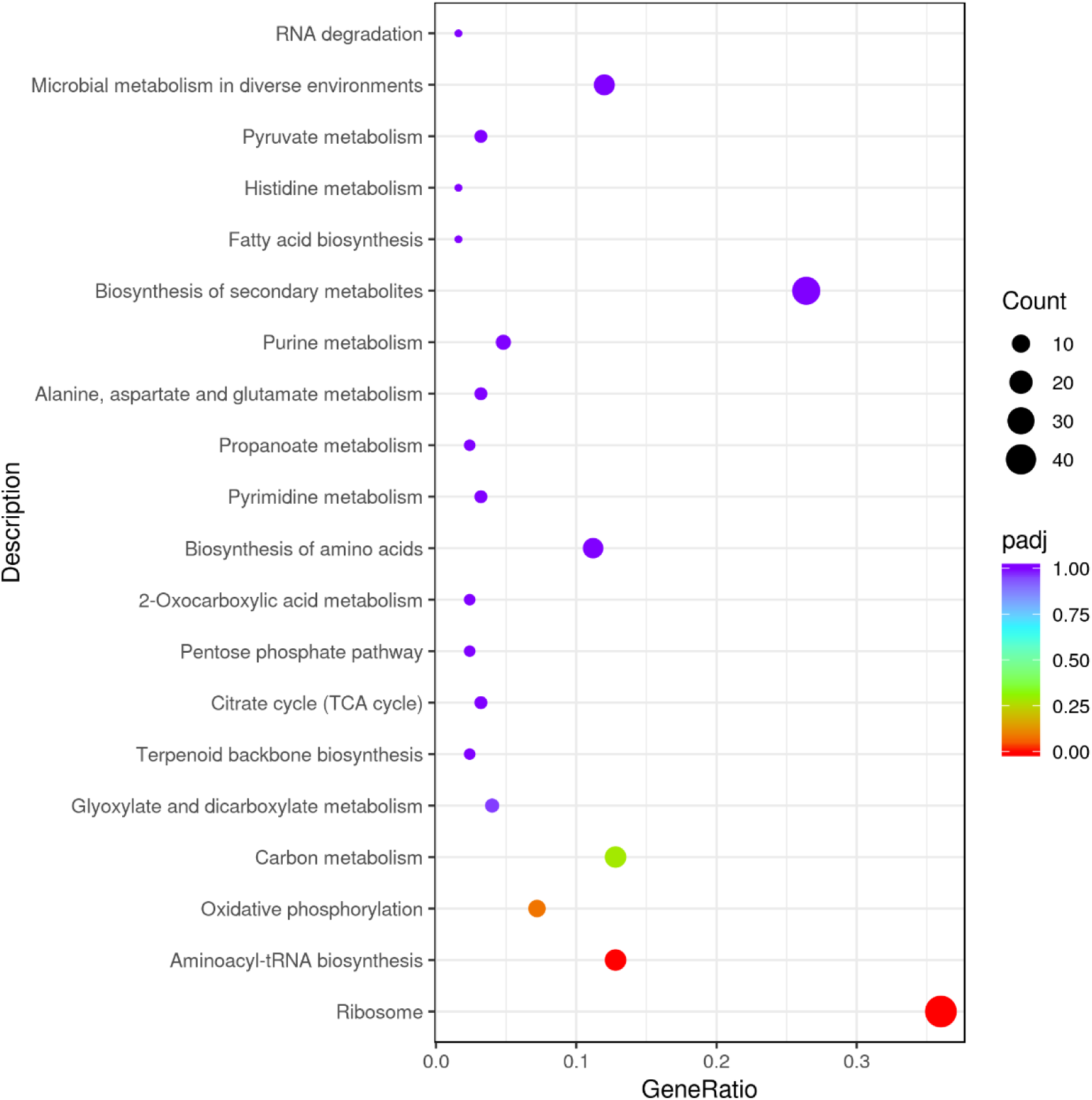
**Up-regulation KEGG pathways enrichment (scatter plot) based on the transcriptomic analysis of PET degradation by *Halomonas* sp. for 8 h.**

**Supplementary Fig. 18.**
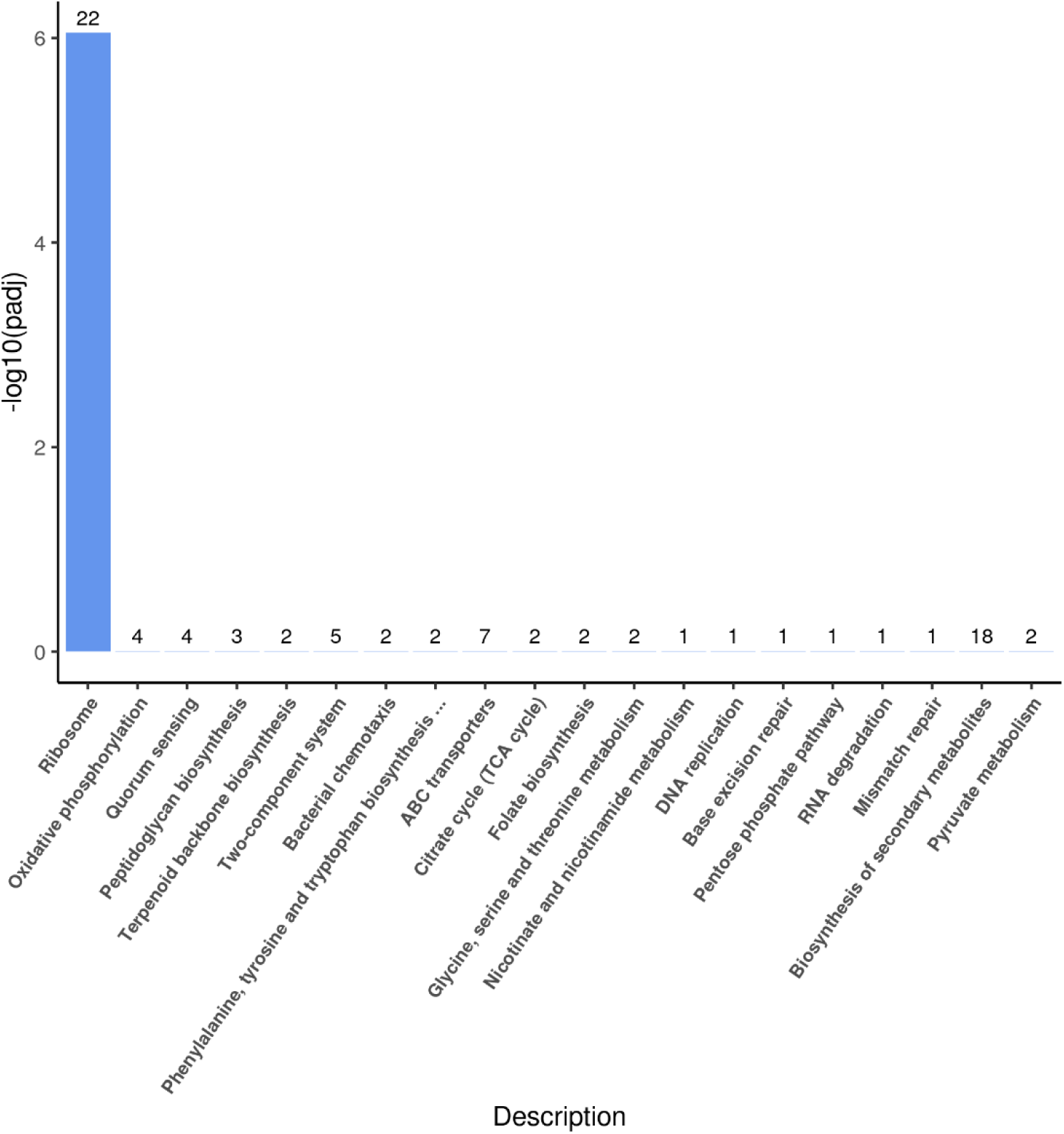
**Up-regulation KEGG pathways enrichment (histogram) based on the transcriptomic analysis of PET degradation by *Halomonas* sp. for 7 d.** The numbers above the column are corresponding genes number related to different pathways.

**Supplementary Fig. 19.**
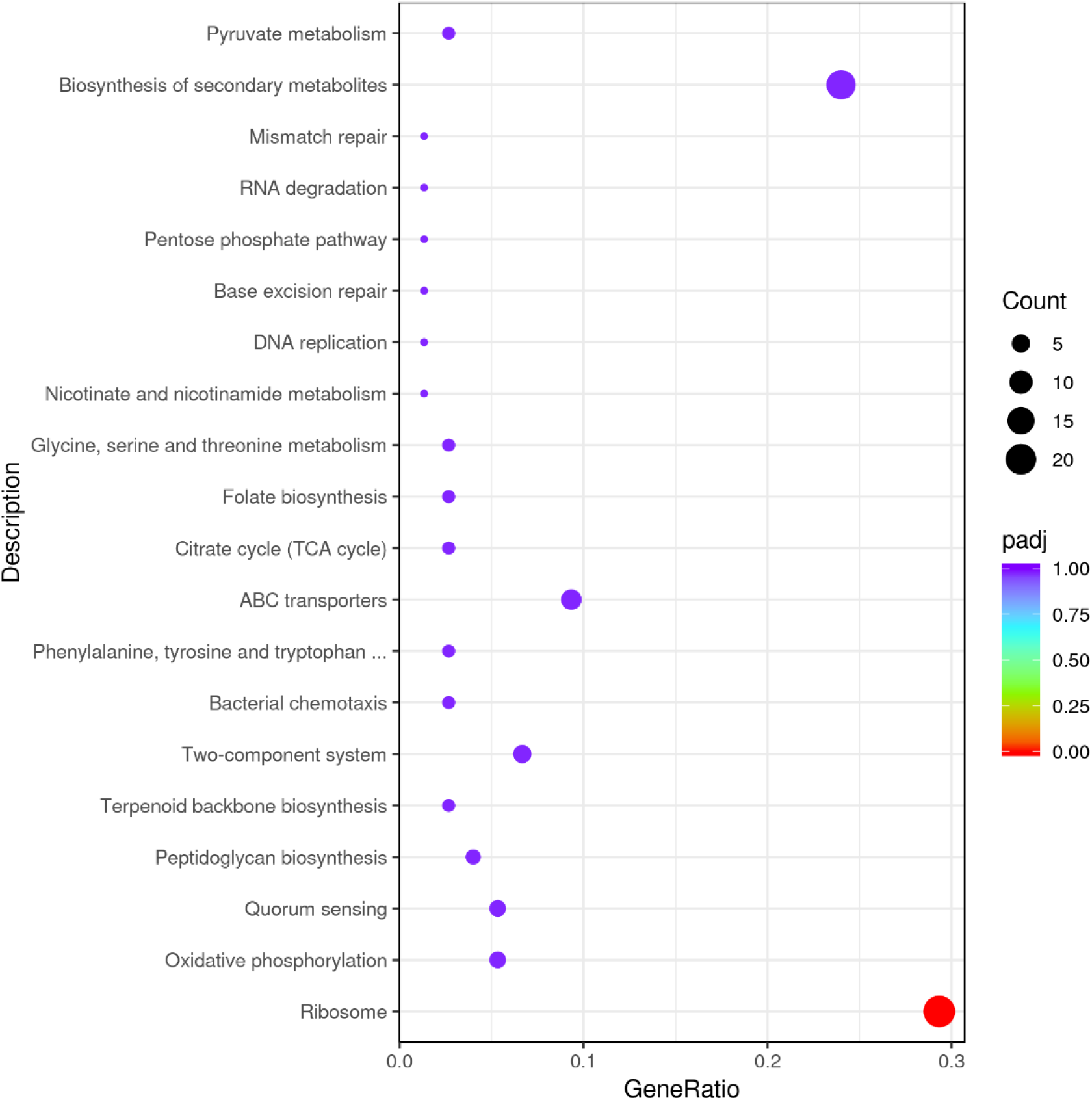
**Up-regulation KEGG pathways enrichment (scatter plot) based on the transcriptomic analysis of PET degradation by *Halomonas* sp. for 7 d.**

**Supplementary Fig. 20.**
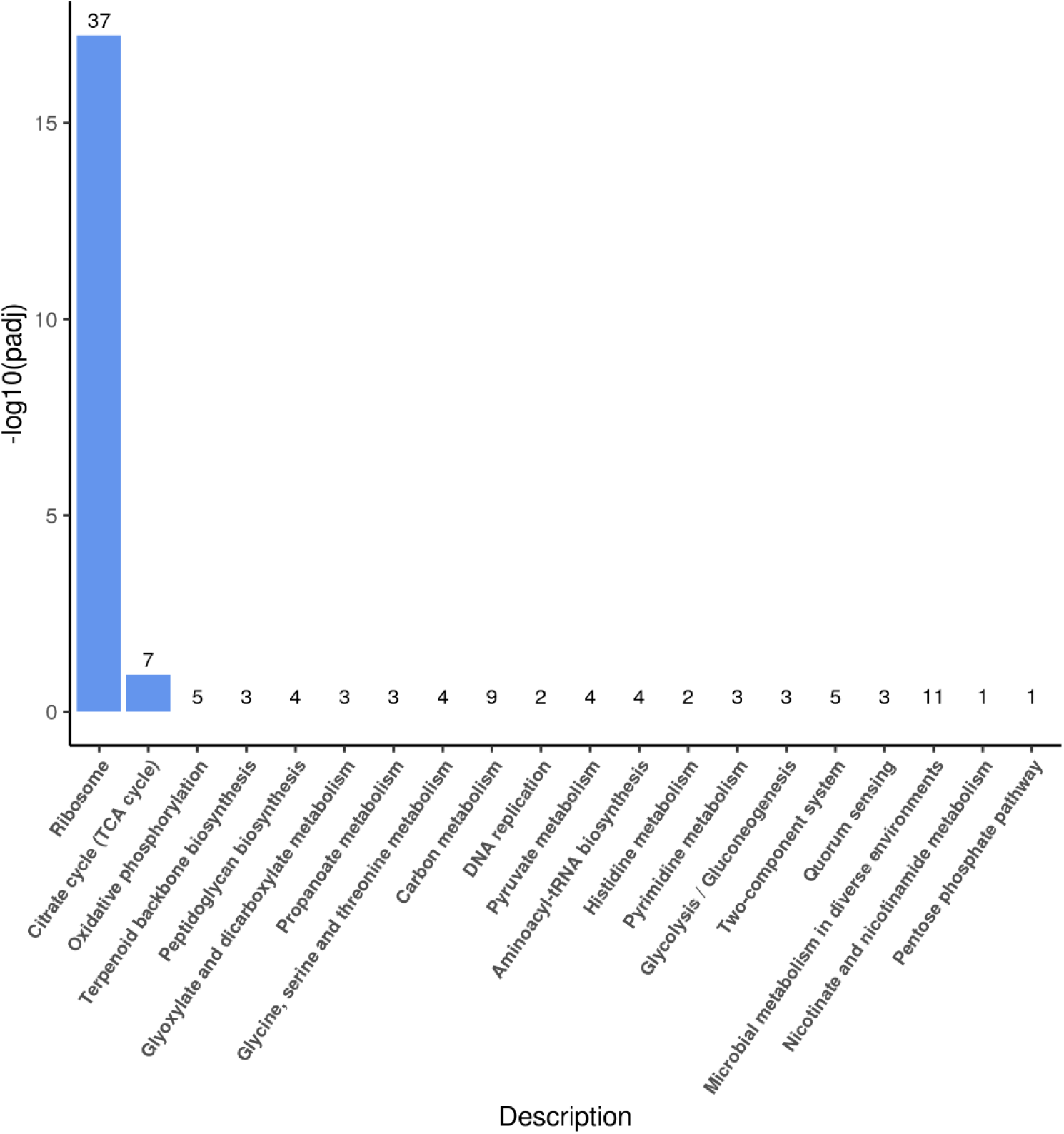
**Up-regulation KEGG pathways enrichment (histogram) based on the transcriptomic analysis of PET degradation by *Halomonas* sp. for 14 d.** The numbers above the column are corresponding genes number related to different pathways.

**Supplementary Fig. 21.**
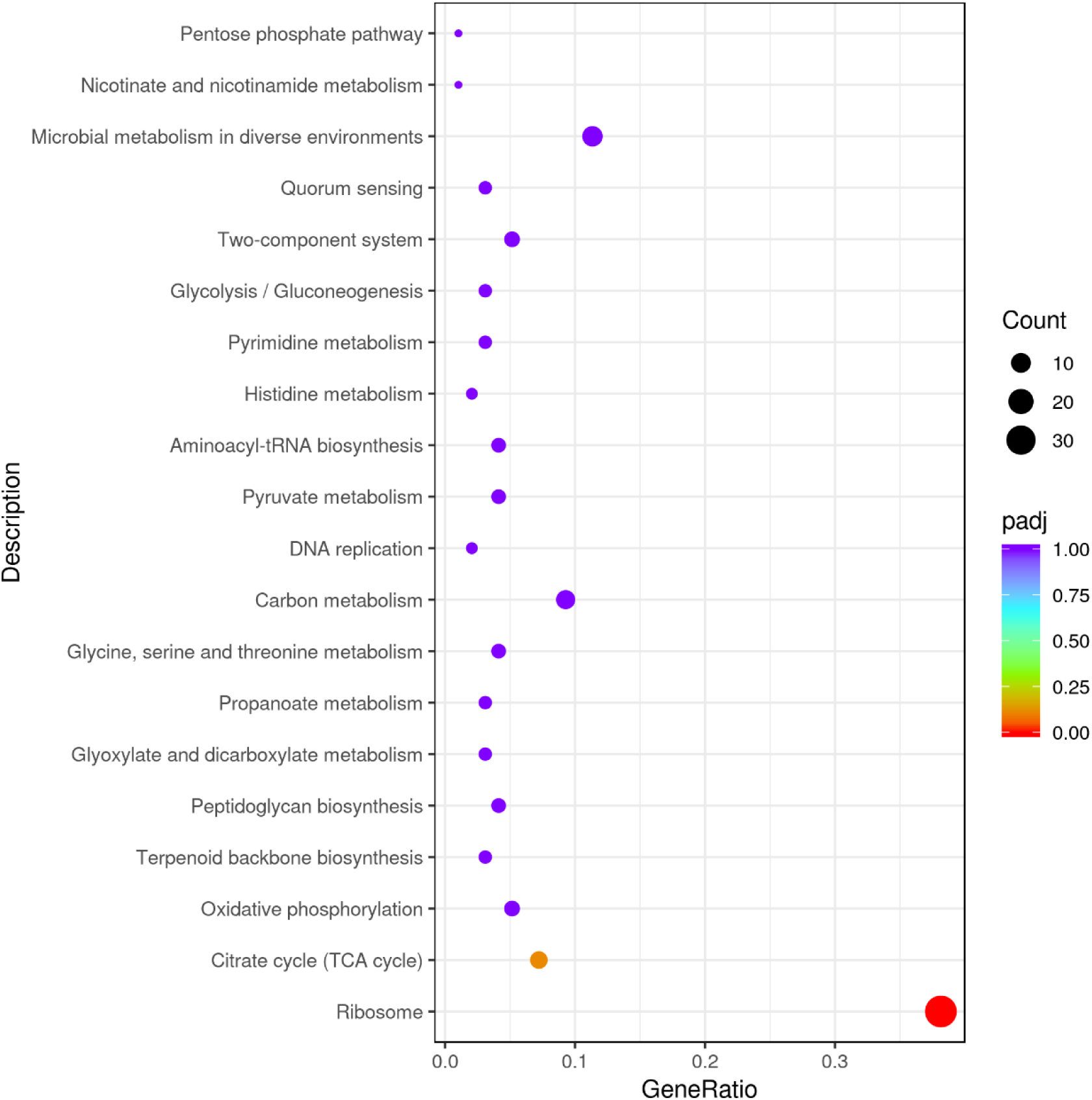
**Up-regulation KEGG pathways enrichment (scatter plot) based on the transcriptomic analysis of PET degradation by *Halomonas* sp. for 14 d.**

**Supplementary Fig. 22.**
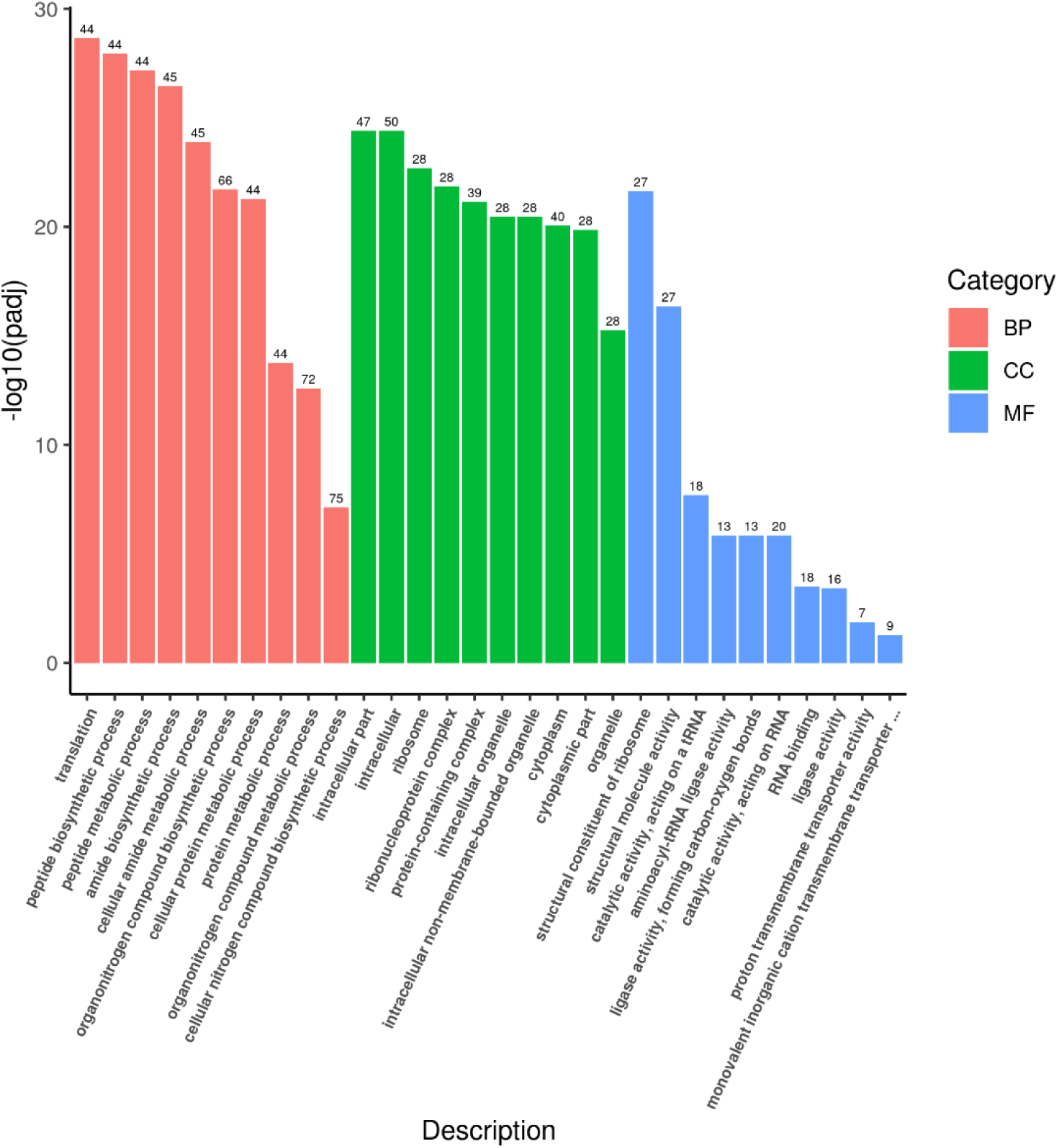
**Up-regulation Go enrichment (histogram) based on the transcriptomic analysis of PET degradation by *Halomonas* sp. for 8 h.** The numbers above the column are corresponding genes number related to different pathways.

**Supplementary Fig. 23.**
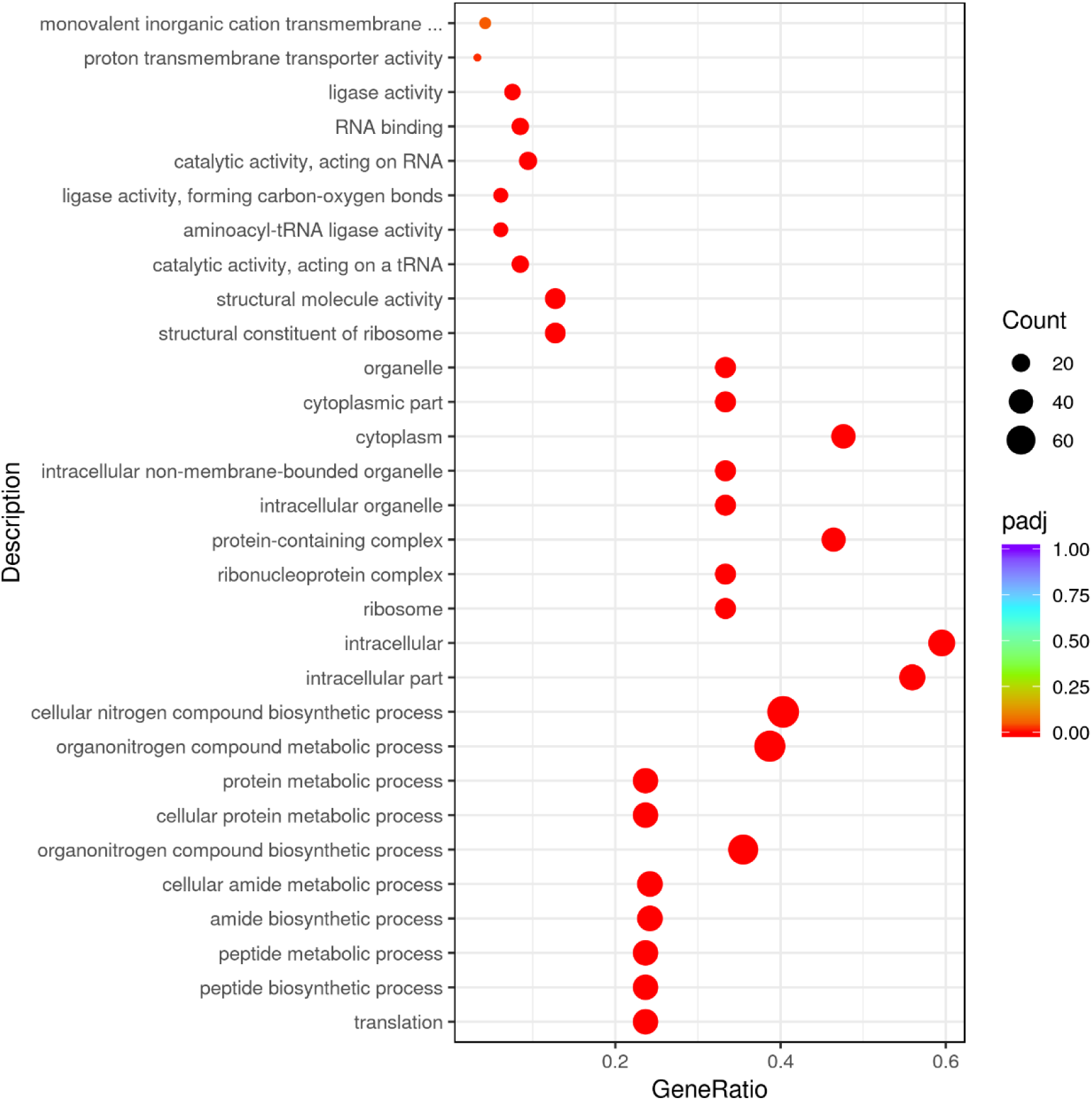
**Up-regulation GO enrichment (scatter plot) based on the transcriptomic analysis of PET degradation by *Halomonas* sp. for 8 h.**

**Supplementary Fig. 24.**
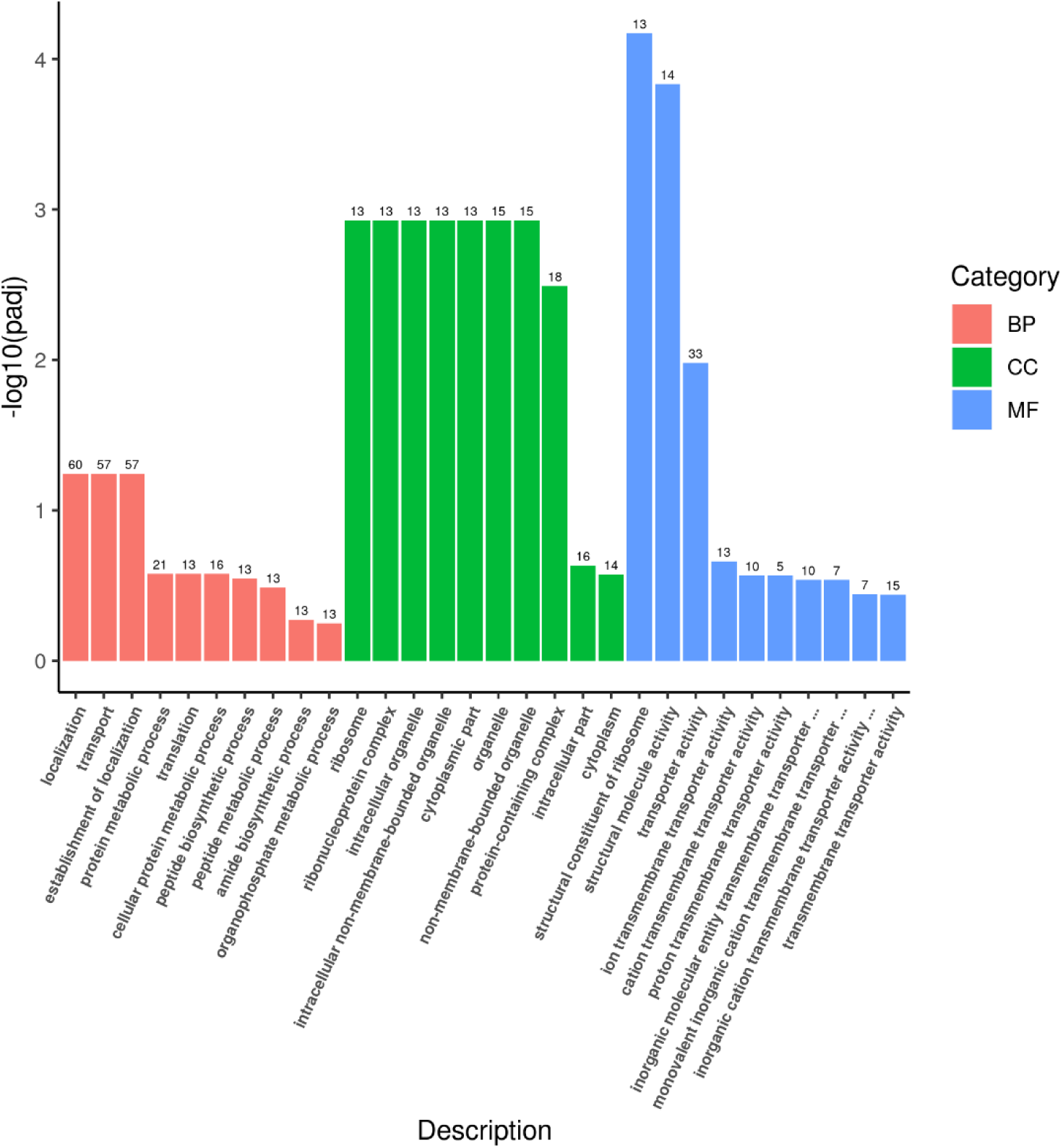
**Up-regulation Go enrichment (histogram) based on the transcriptomic analysis of PET degradation by *Halomonas* sp. for 7 d.** The numbers above the column are corresponding genes number related to different pathways.

**Supplementary Fig. 25.**
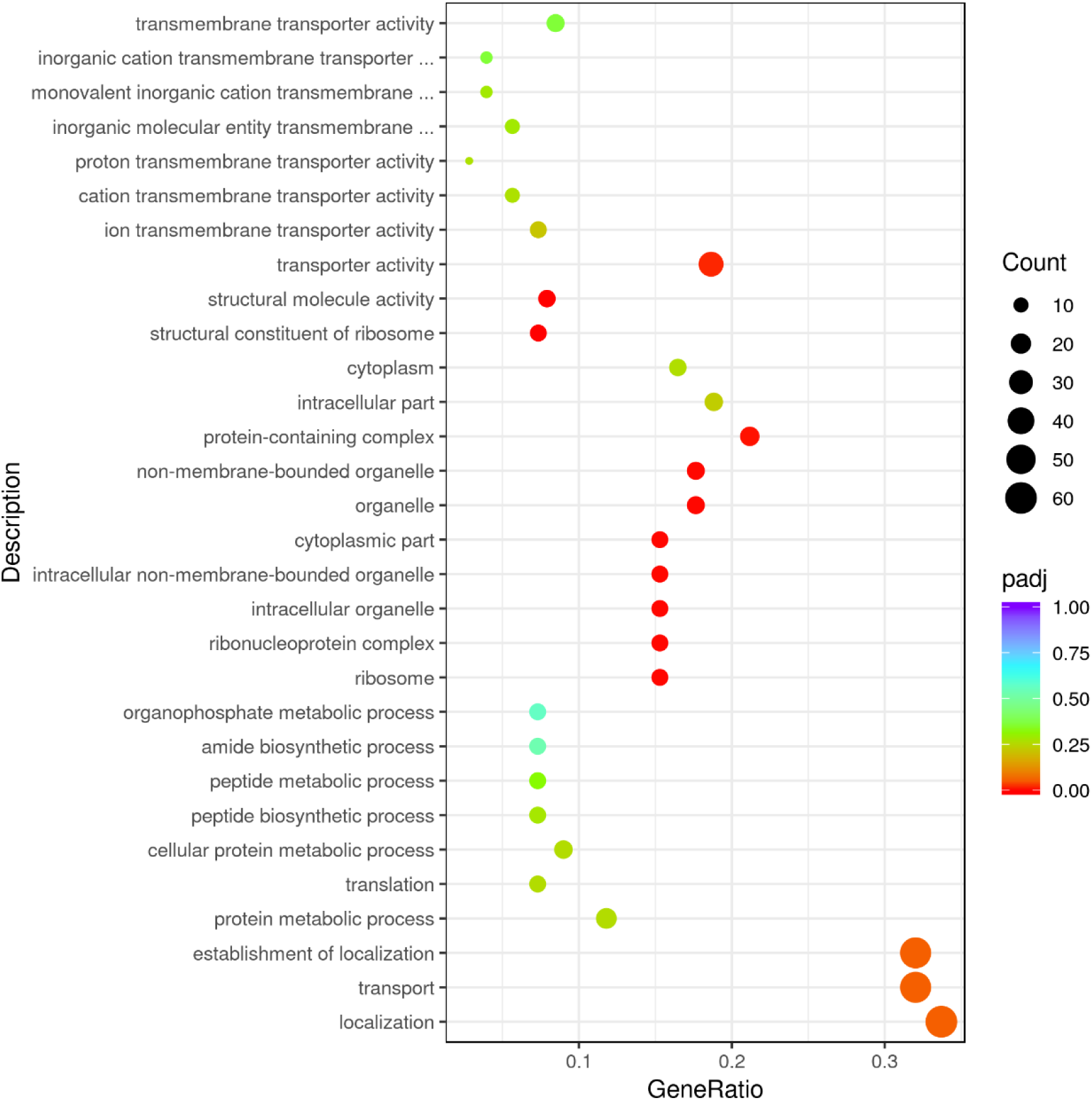
**Up-regulation GO enrichment (scatter plot) based on the transcriptomic analysis of PET degradation by *Halomonas* sp. for 7 d.**

**Supplementary Fig. 26.**
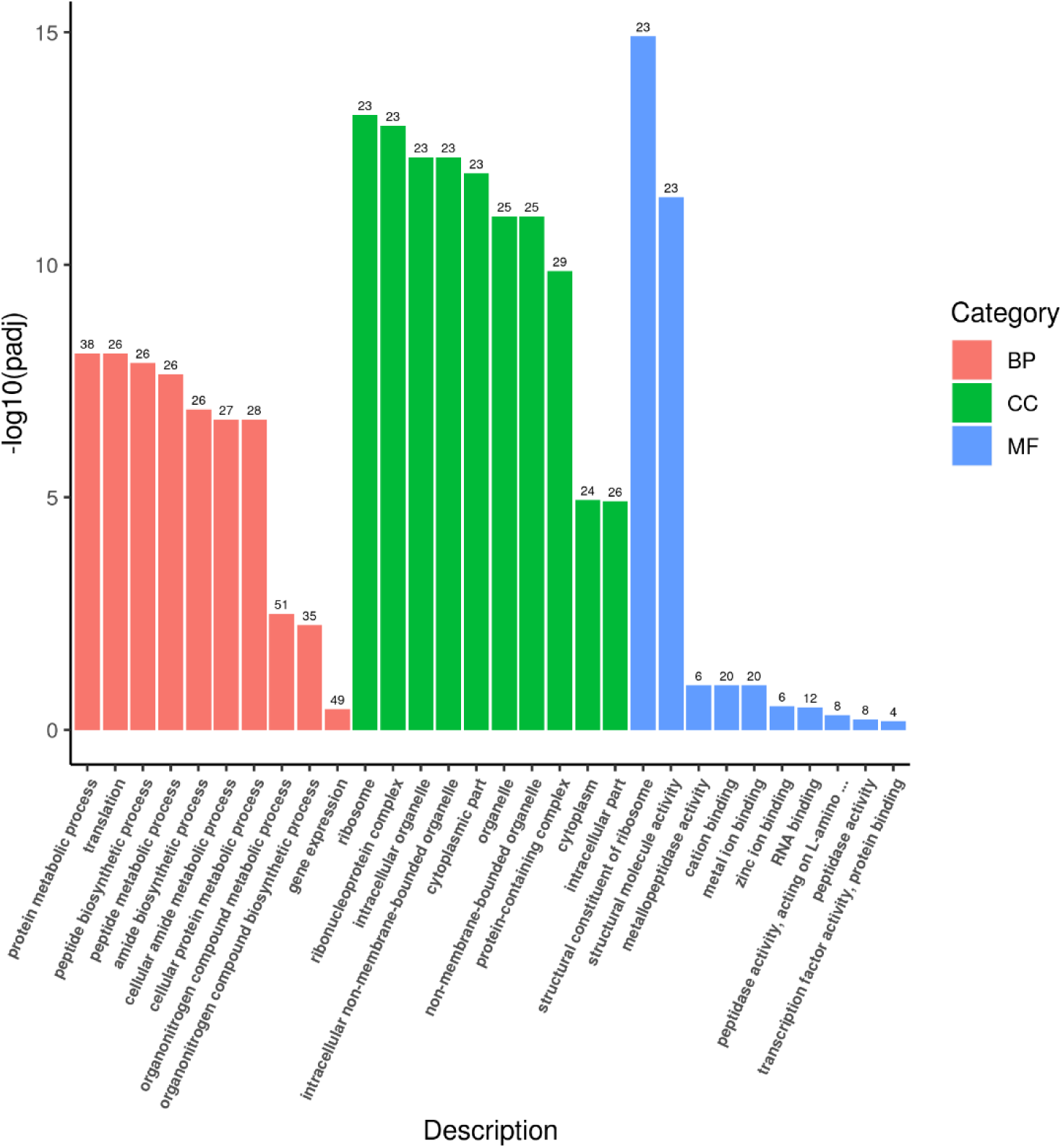
**Up-regulation Go enrichment (histogram) based on the transcriptomic analysis of PET degradation by *Halomonas* sp. for 14 d.** The numbers above the column are corresponding genes number related to different pathways.

**Supplementary Fig. 27.**
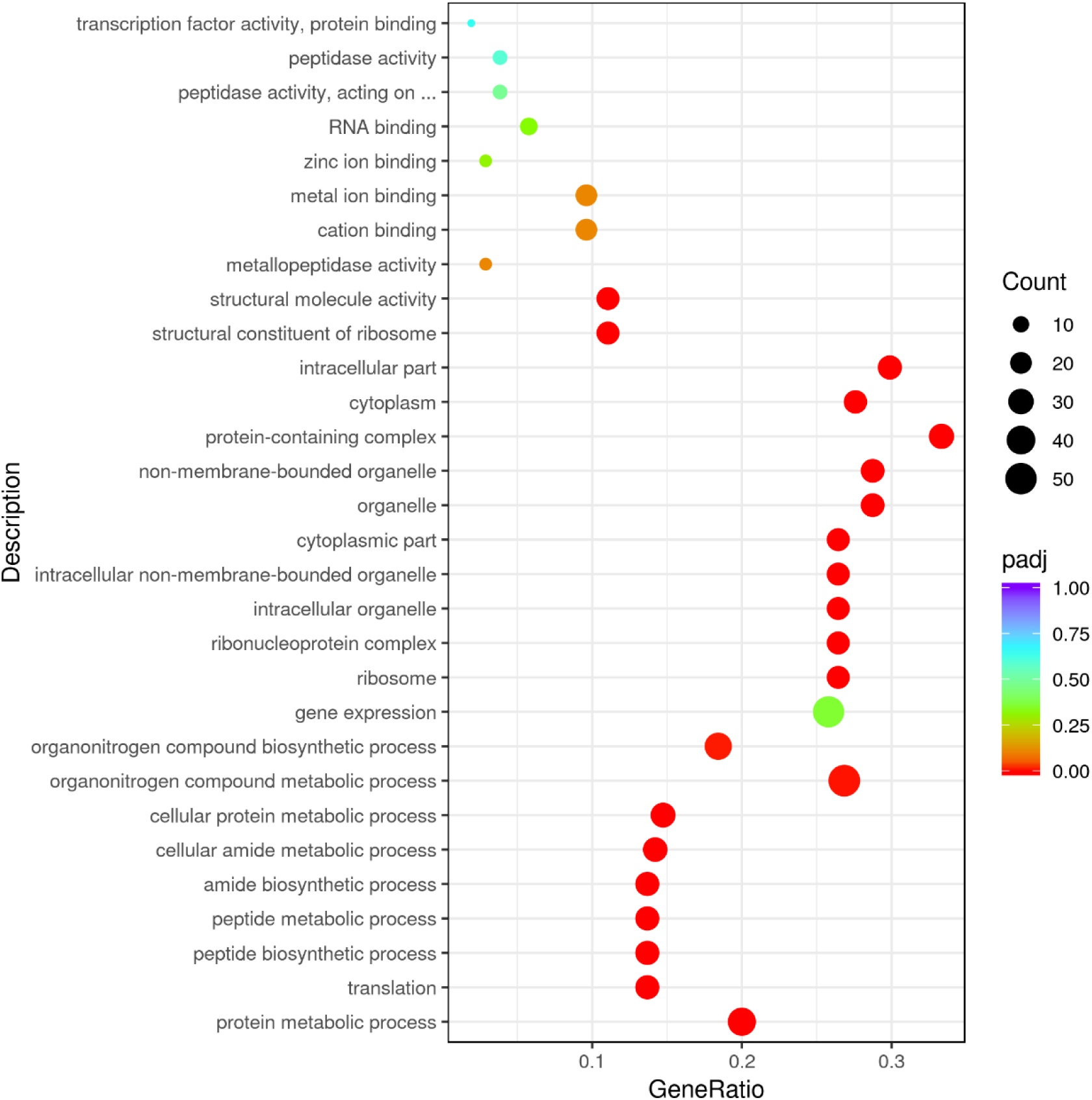
**Up-regulation GO enrichment (scatter plot) based on the transcriptomic analysis of PET degradation by *Halomonas* sp. for 14 d.**

**Supplementary Fig. 28.**
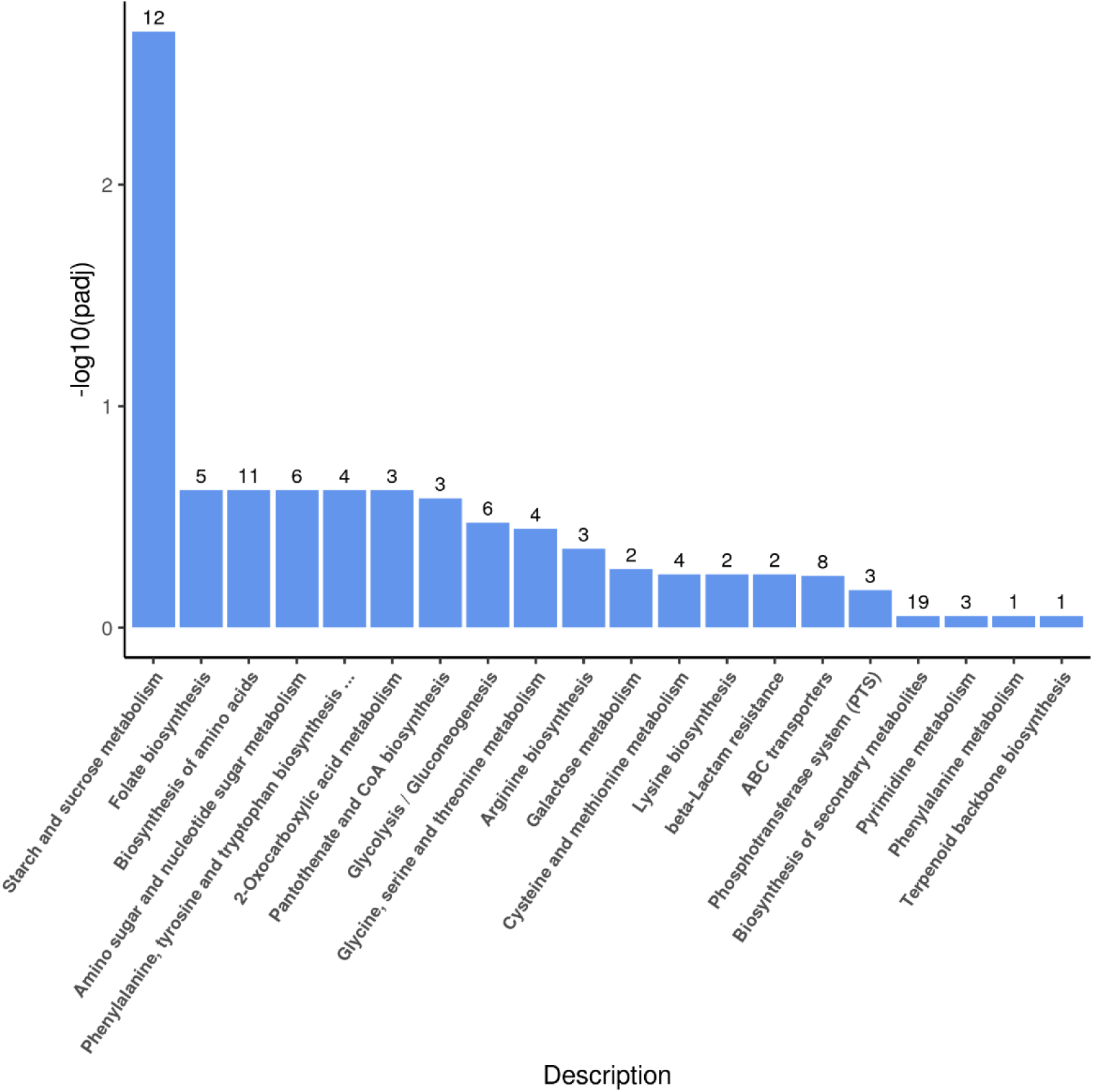
**Up-regulation KEGG pathways enrichment (histogram) based on the transcriptomic analysis of PE degradation by *Exiguobacterium* sp. for 8 h.** The numbers above the column are corresponding genes number related to different pathways.

**Supplementary Fig. 29.**
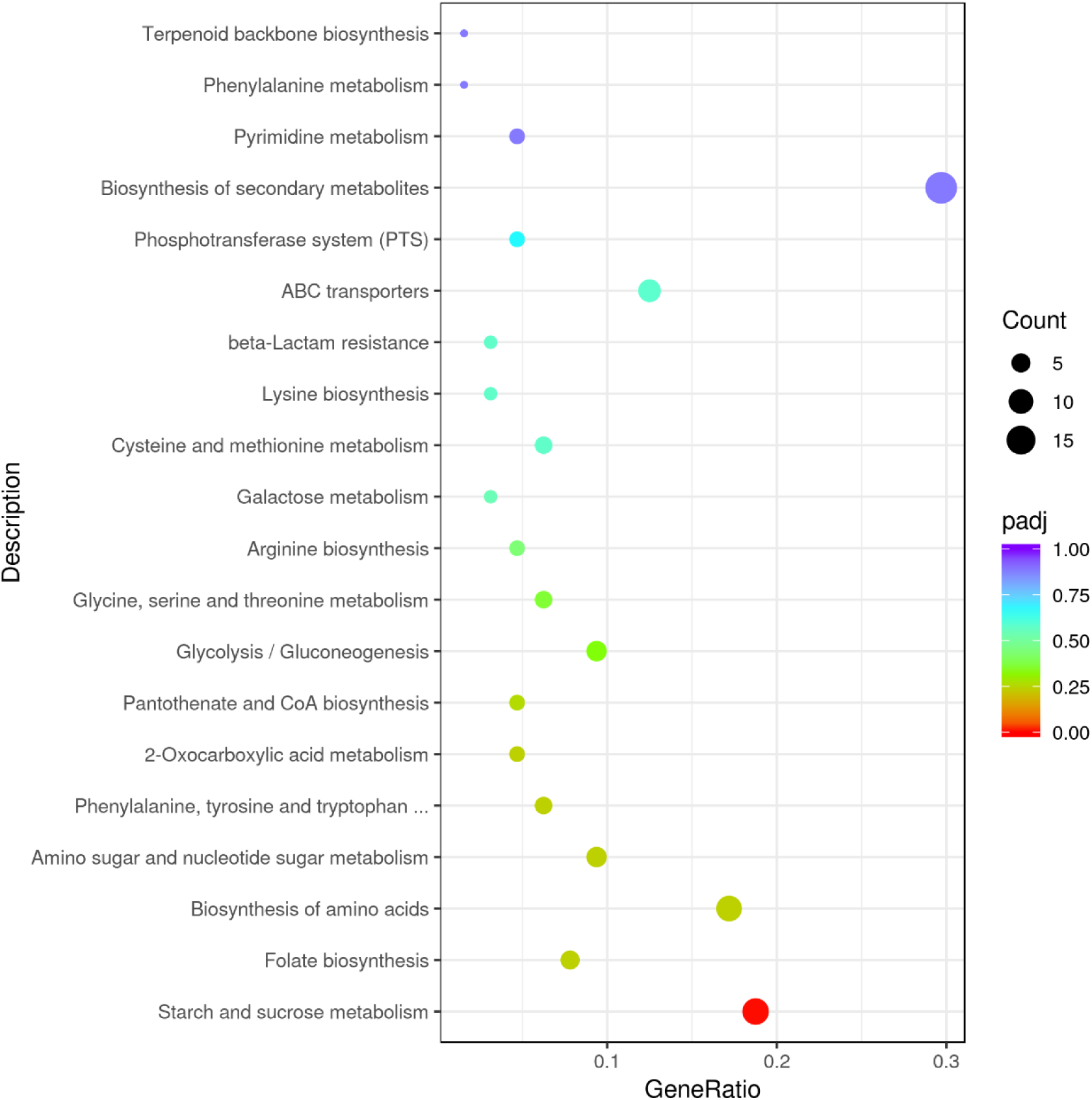
**Up-regulation KEGG pathways enrichment (scatter plot) based on the transcriptomic analysis of PE degradation by *Exiguobacterium* sp. for 8 h.**

**Supplementary Fig. 30.**
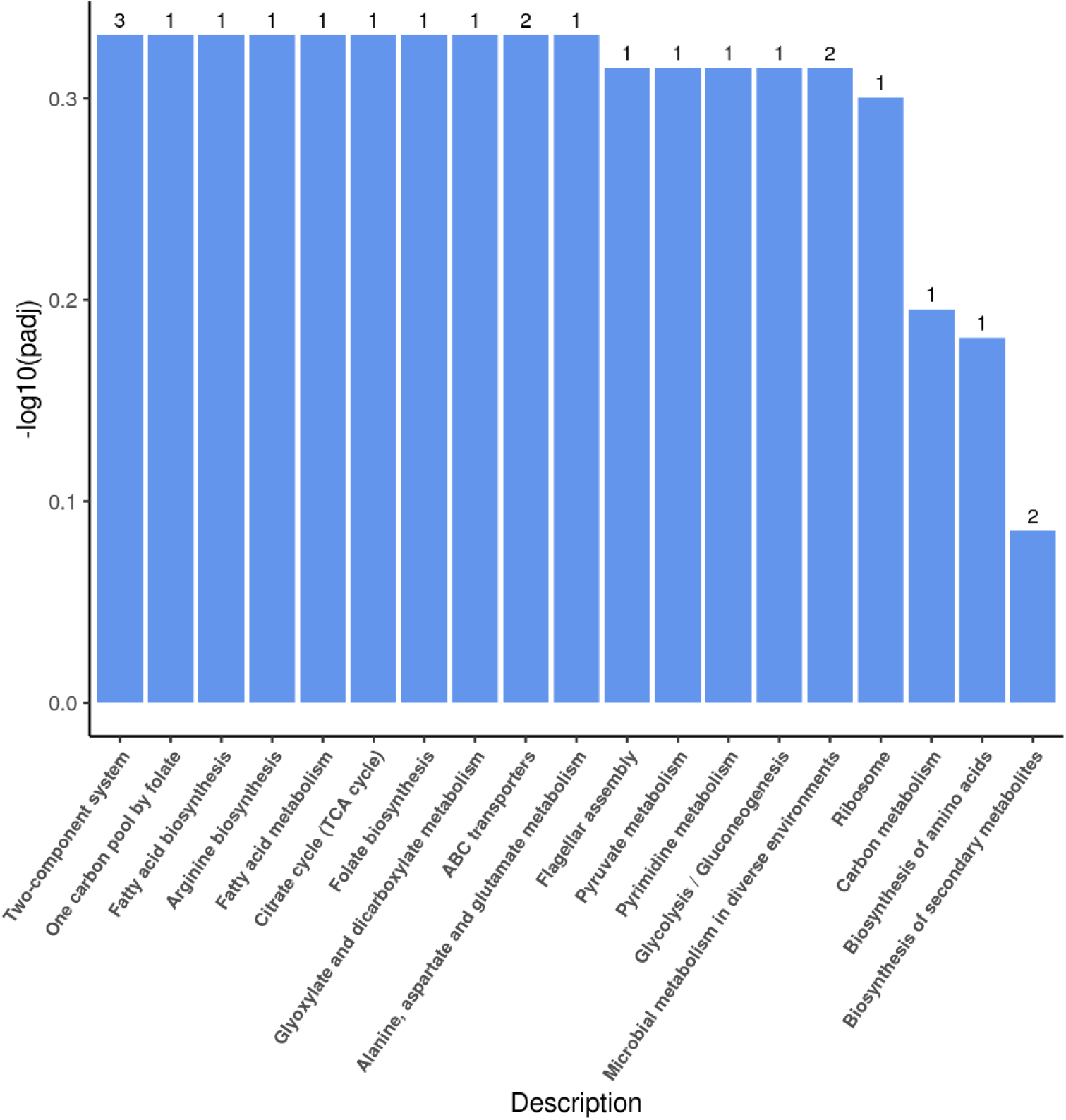
**Up-regulation KEGG pathways enrichment (histogram) based on the transcriptomic analysis of PE degradation by *Exiguobacterium* sp. for 7 d.** The numbers above the column are corresponding genes number related to different pathways.

**Supplementary Fig. 31.**
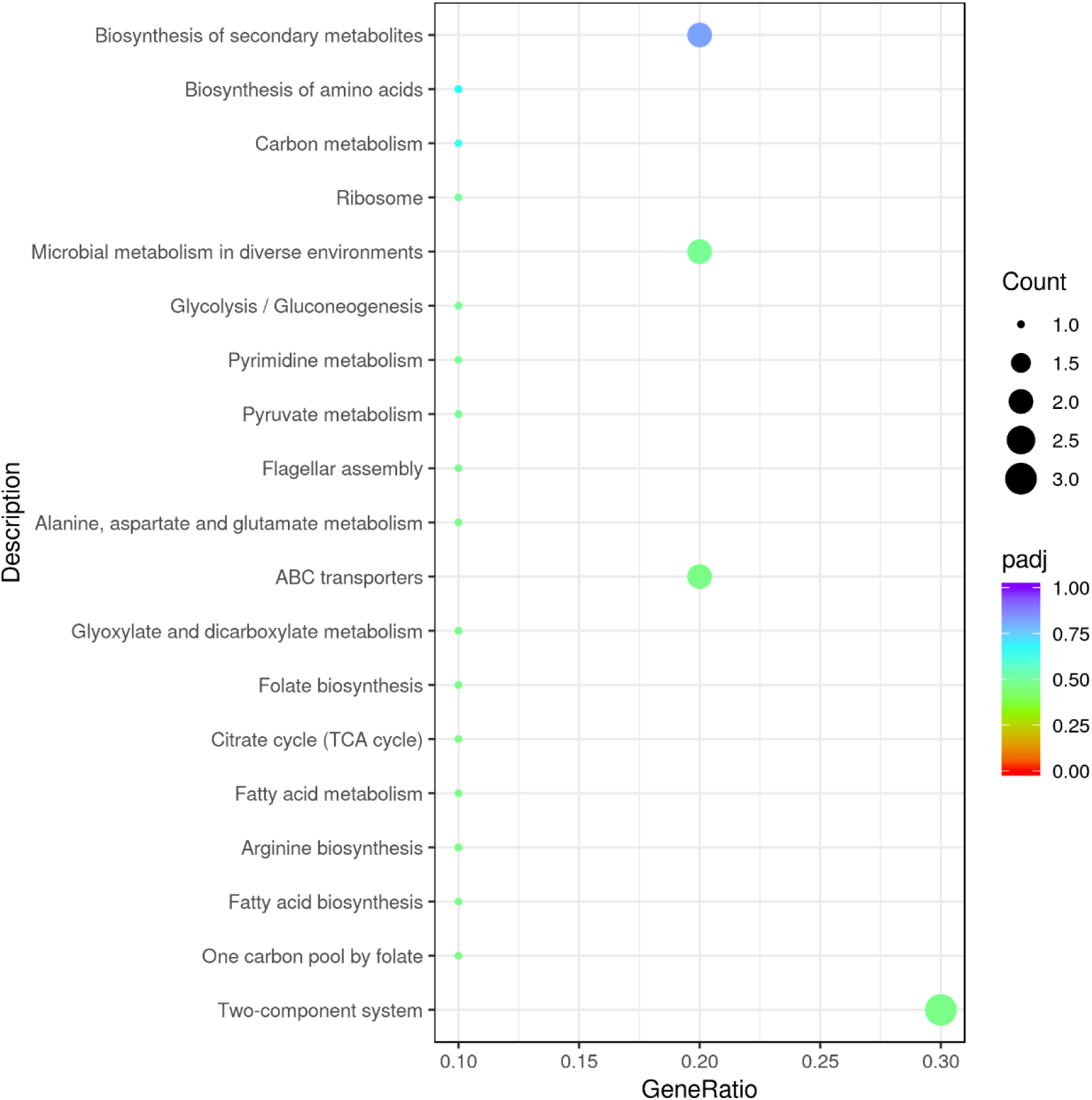
**Up-regulation KEGG pathways enrichment (scatter plot) based on the transcriptomic analysis of PE degradation by *Exiguobacterium* sp. for 7 d.**

**Supplementary Fig. 32.**
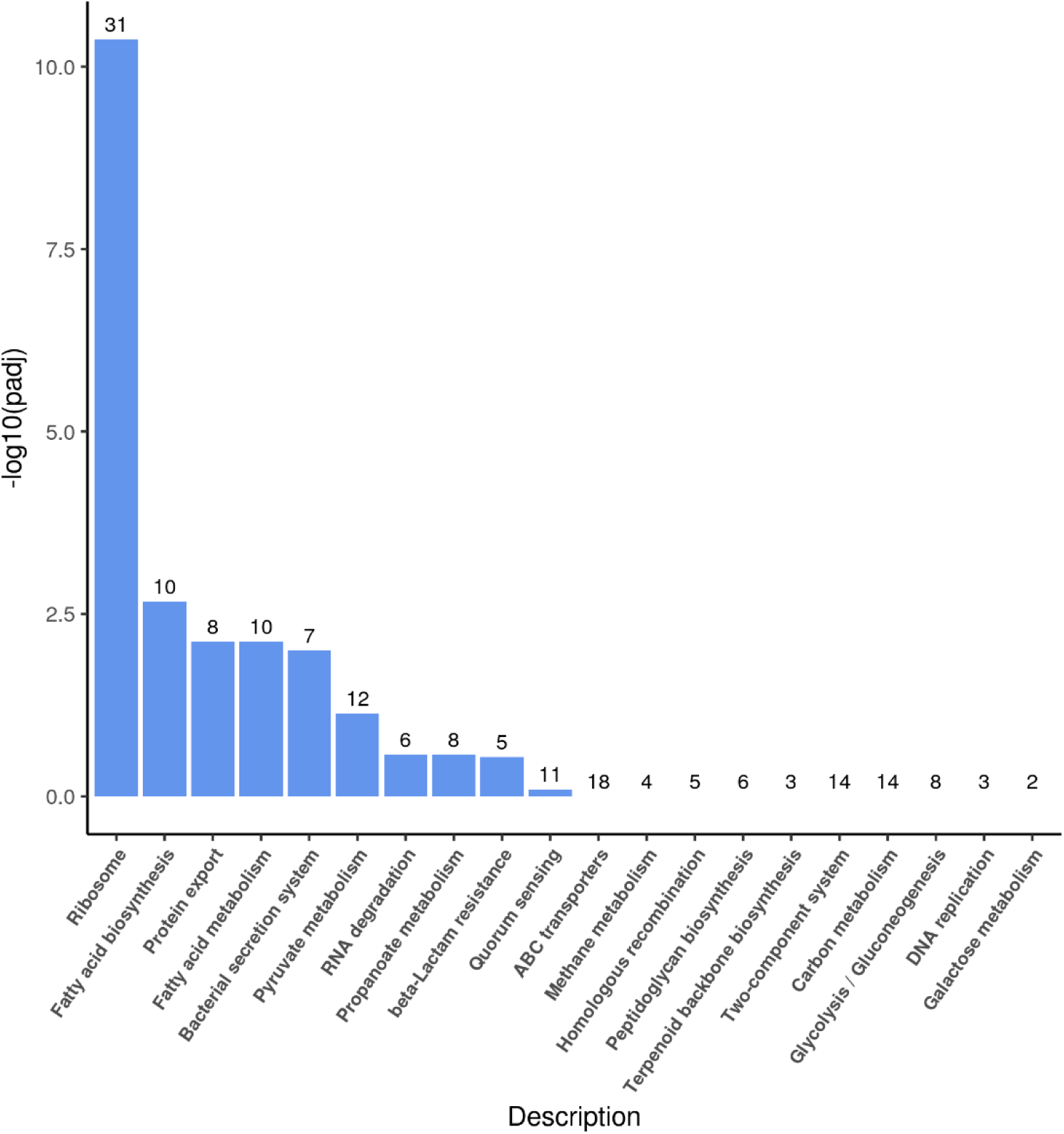
**Up-regulation KEGG pathways enrichment (histogram) based on the transcriptomic analysis of PE degradation by *Exiguobacterium* sp. for 14 d.** The numbers above the column are corresponding genes number related to different pathways.

**Supplementary Fig. 33.**
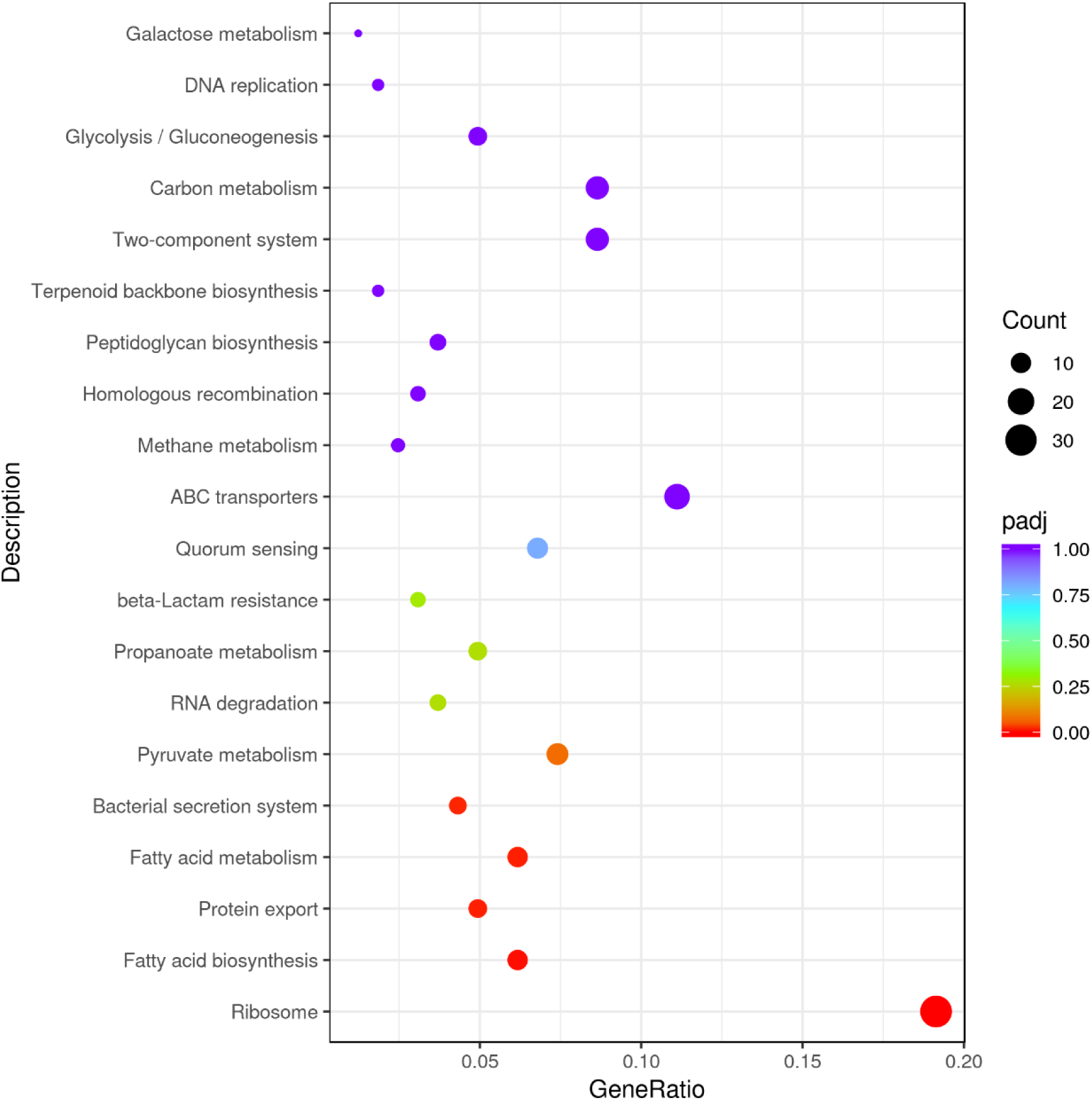
**Up-regulation KEGG pathways enrichment (scatter plot) based on the transcriptomic analysis of PE degradation by *Exiguobacterium* sp. for 14 d.**

**Supplementary Fig. 34.**
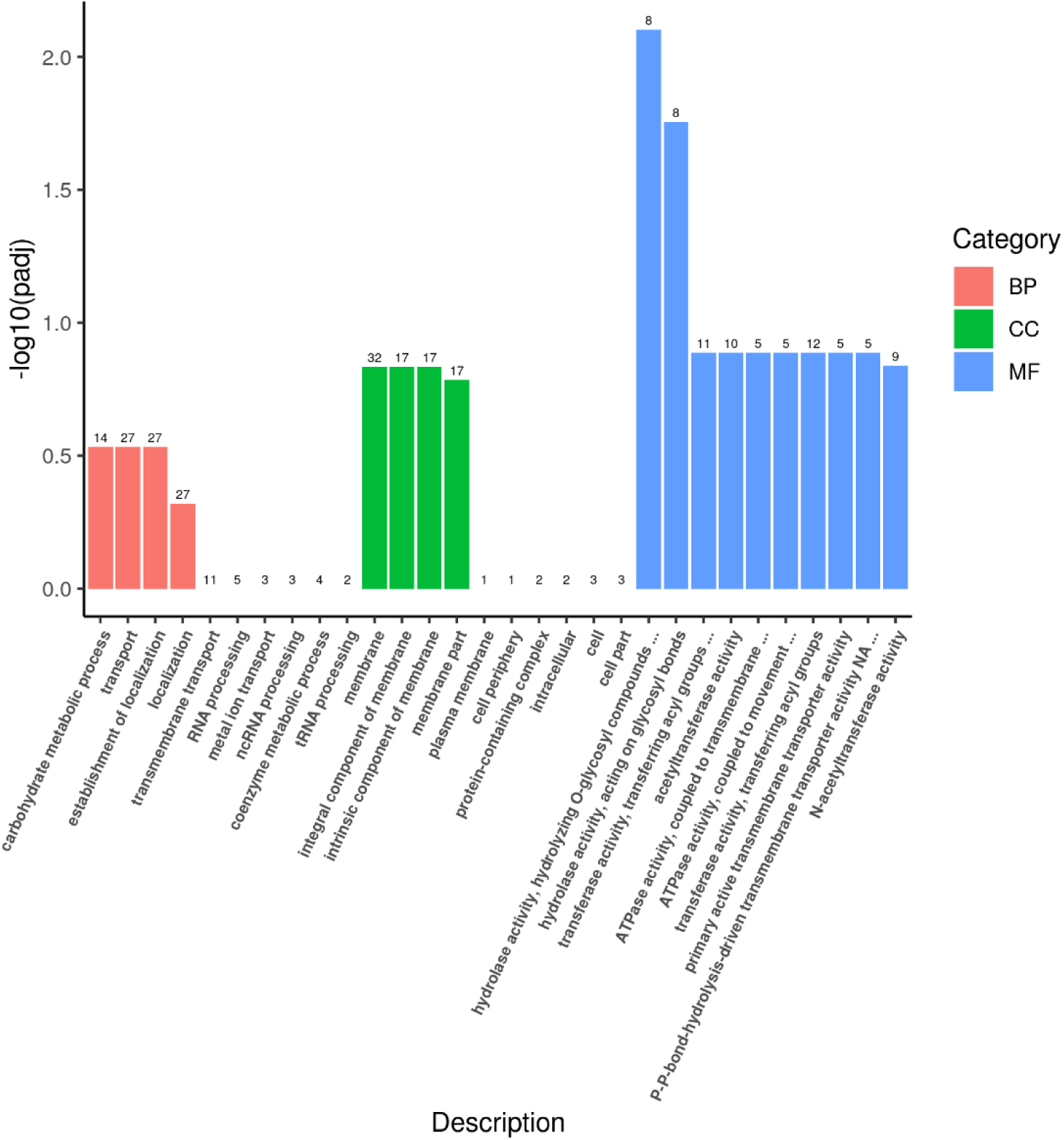
**Up-regulation Go enrichment (histogram) based on the transcriptomic analysis of PE degradation by *Exiguobacterium* sp. for 8 h.** The numbers above the column are corresponding genes number related to different pathways.

**Supplementary Fig. 35.**
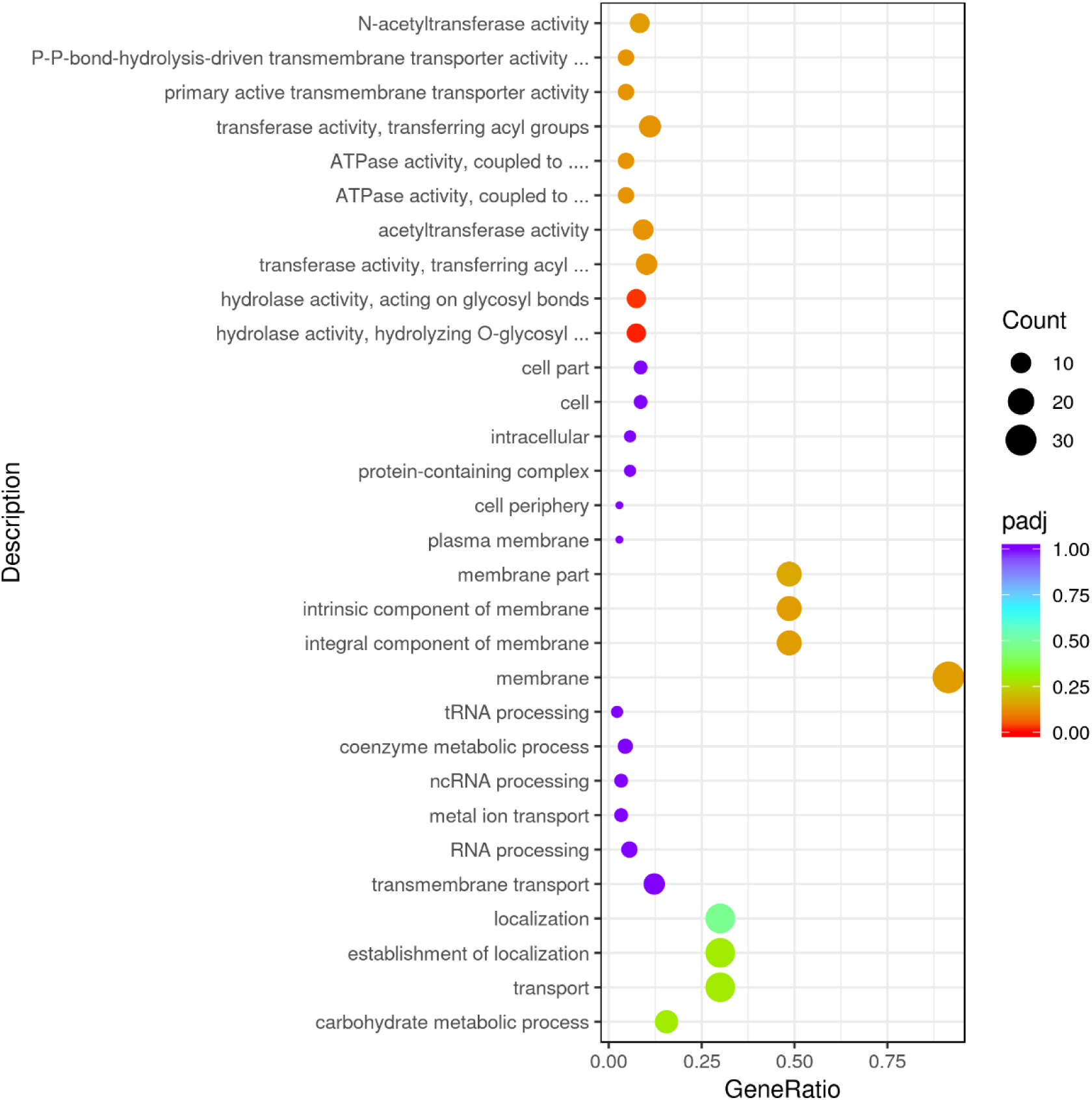
**Up-regulation GO enrichment (scatter plot) based on the transcriptomic analysis of PE degradation by *Exiguobacterium* sp. for 8 h.**

**Supplementary Fig. 36.**
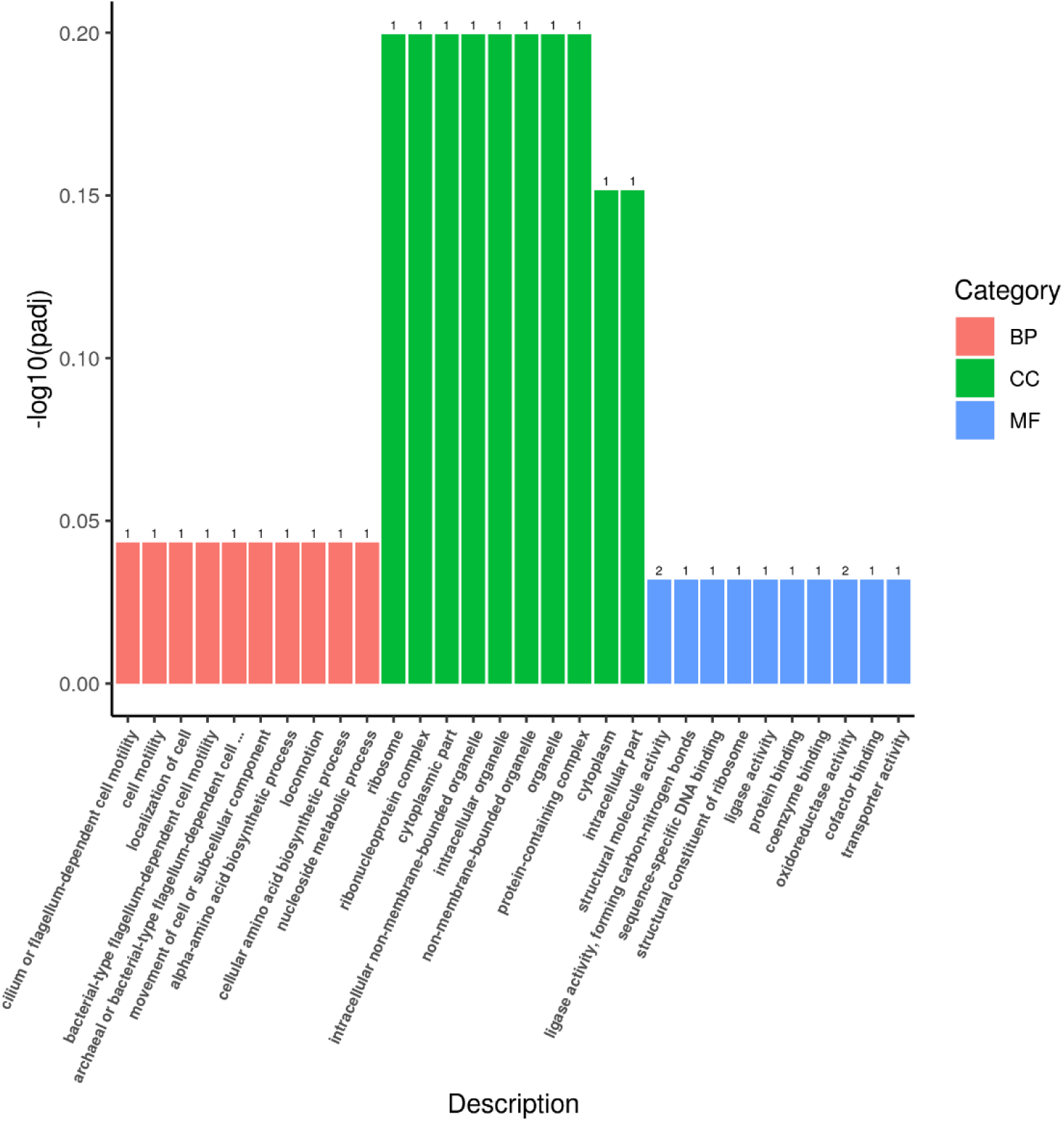
**Up-regulation Go enrichment (histogram) based on the transcriptomic analysis of PE degradation by *Exiguobacterium* sp. for 7 d.** The numbers above the column are corresponding genes number related to different pathways.

**Supplementary Fig. 37.**
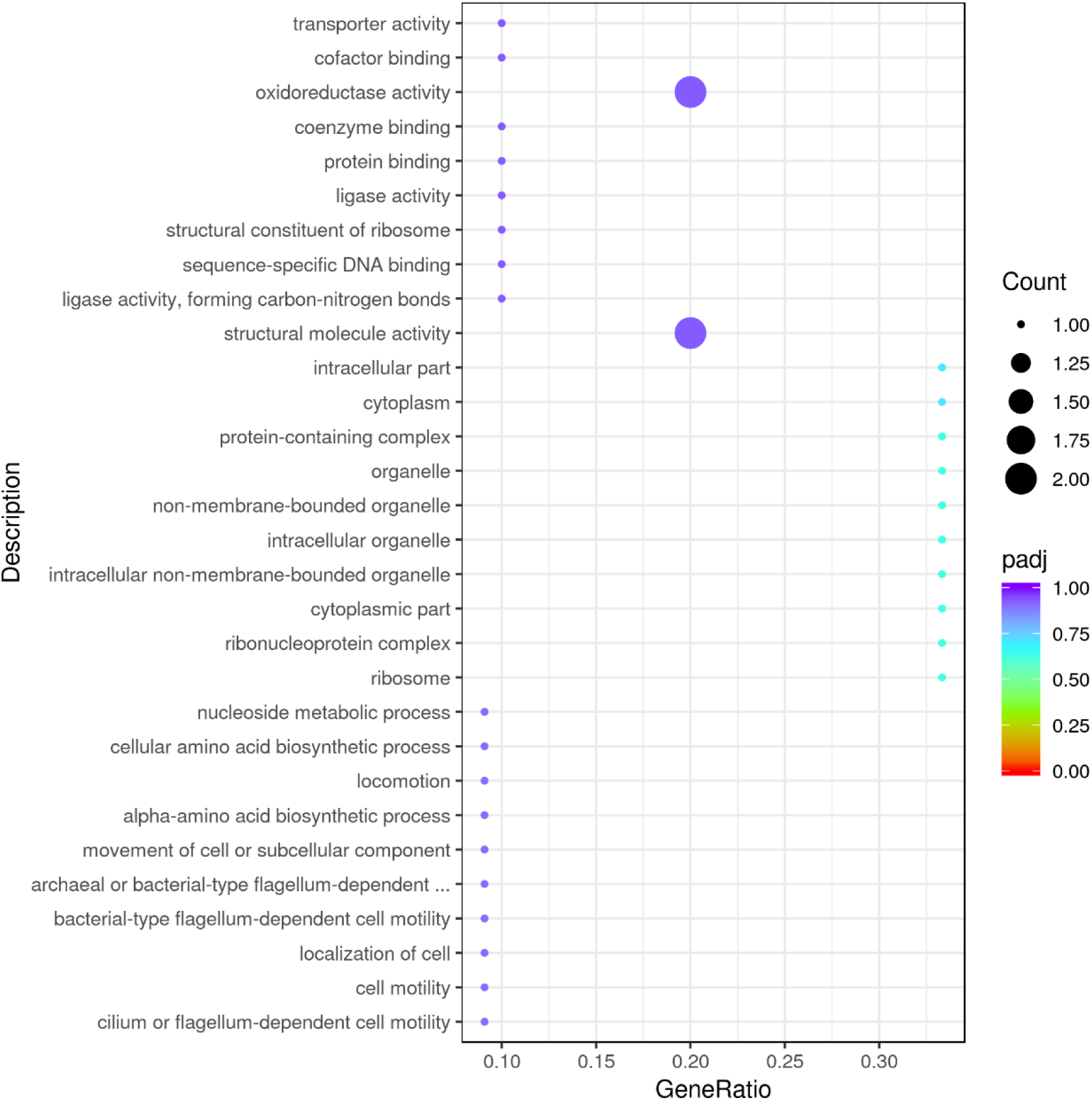
**Up-regulation GO enrichment (scatter plot) based on the transcriptomic analysis of PE degradation by *Exiguobacterium* sp. for 7 d.**

**Supplementary Fig. 38.**
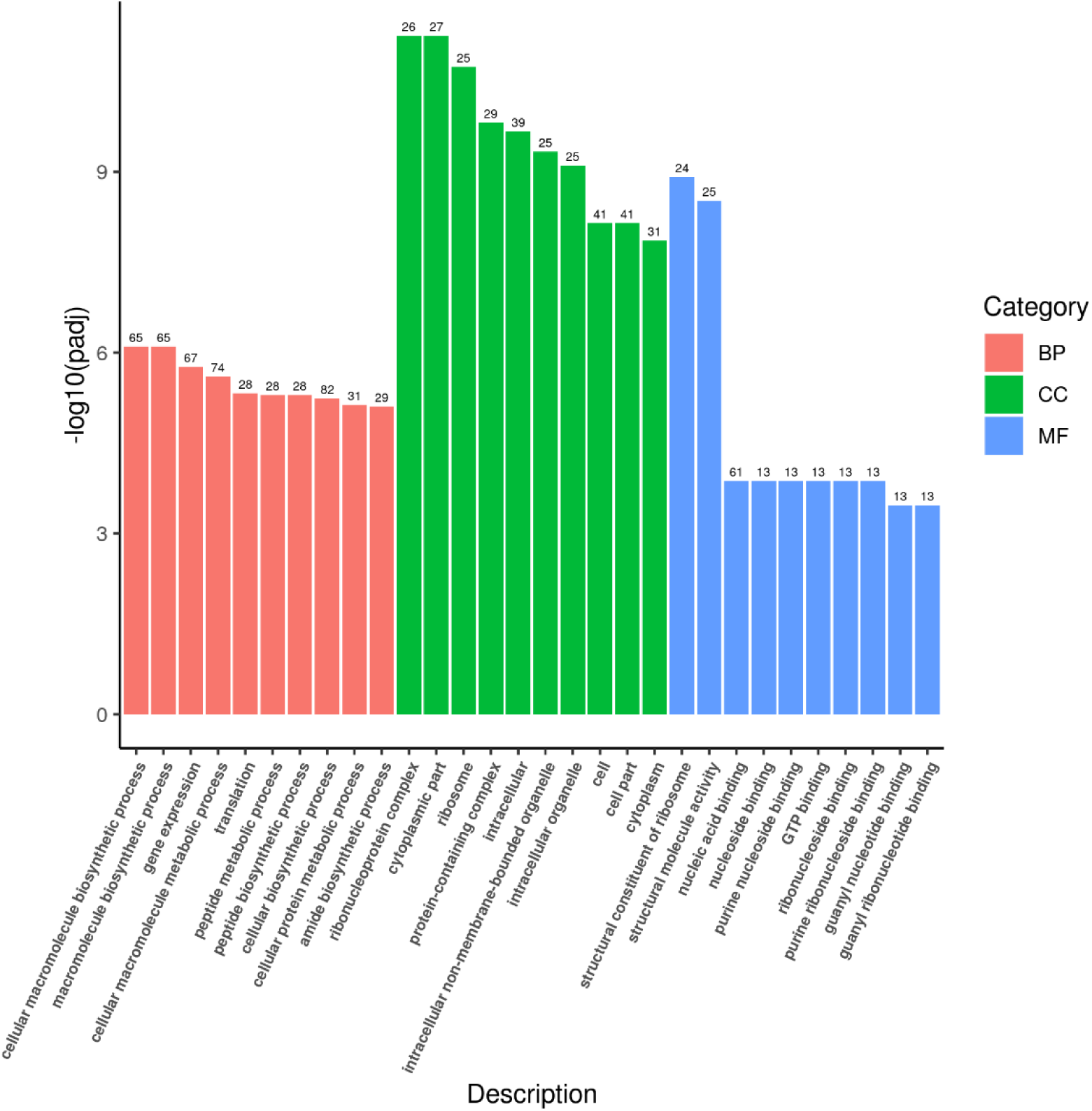
**Up-regulation Go enrichment (histogram) based on the transcriptomic analysis of PE degradation by *Exiguobacterium* sp. for 14 d.** The numbers above the column are corresponding genes number related to different pathways.

**Supplementary Fig. 39.**
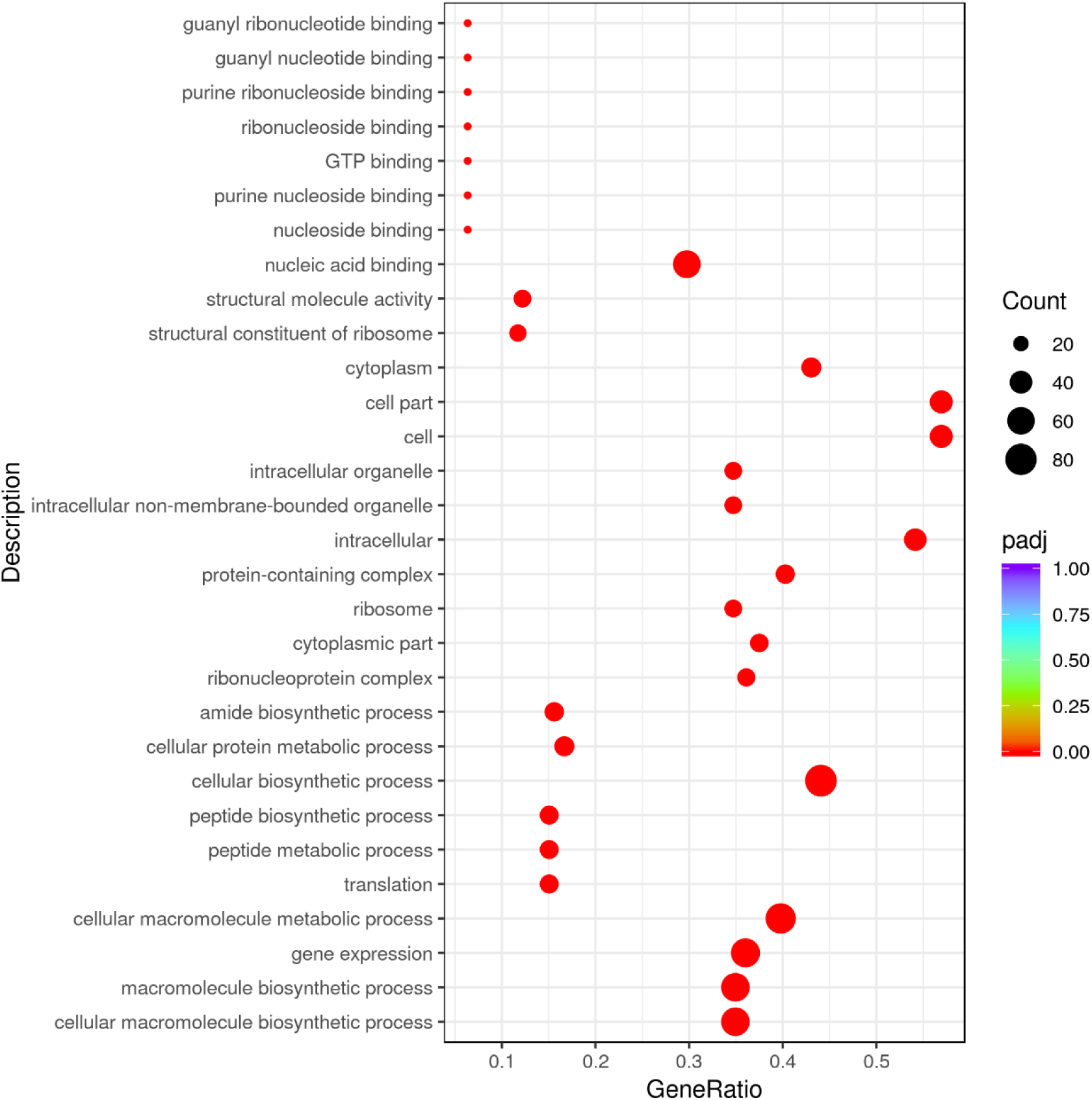
**Up-regulation GO enrichment (scatter plot) based on the transcriptomic analysis of PE degradation by *Exiguobacterium* sp. for 14 d.**

**Supplementary Fig. 40.**
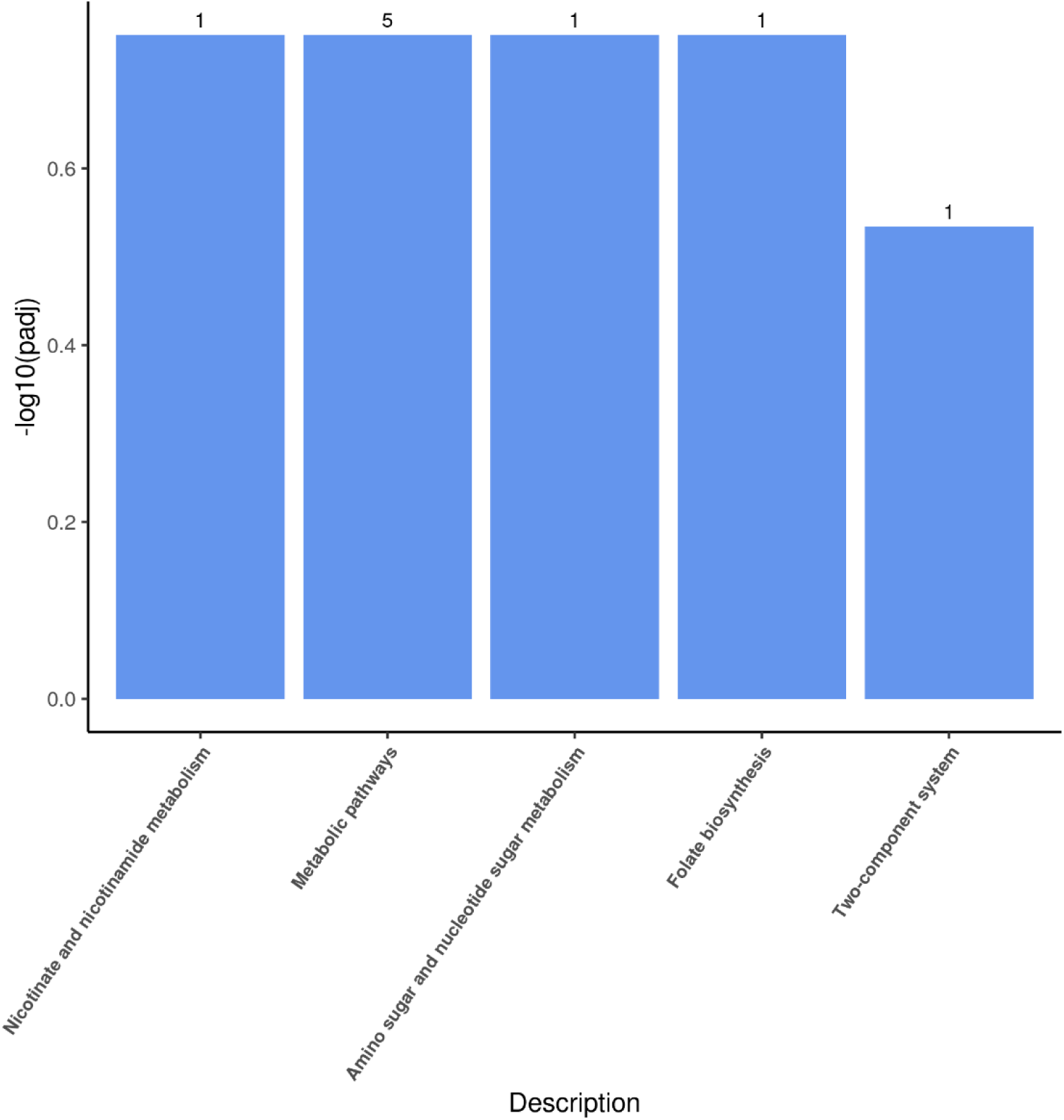
**Up-regulation KEGG pathways enrichment (histogram) based on the transcriptomic analysis of PE degradation by *Halomonas* sp. for 8 h.** The numbers above the column are corresponding genes number related to different pathways.

**Supplementary Fig. 41.**
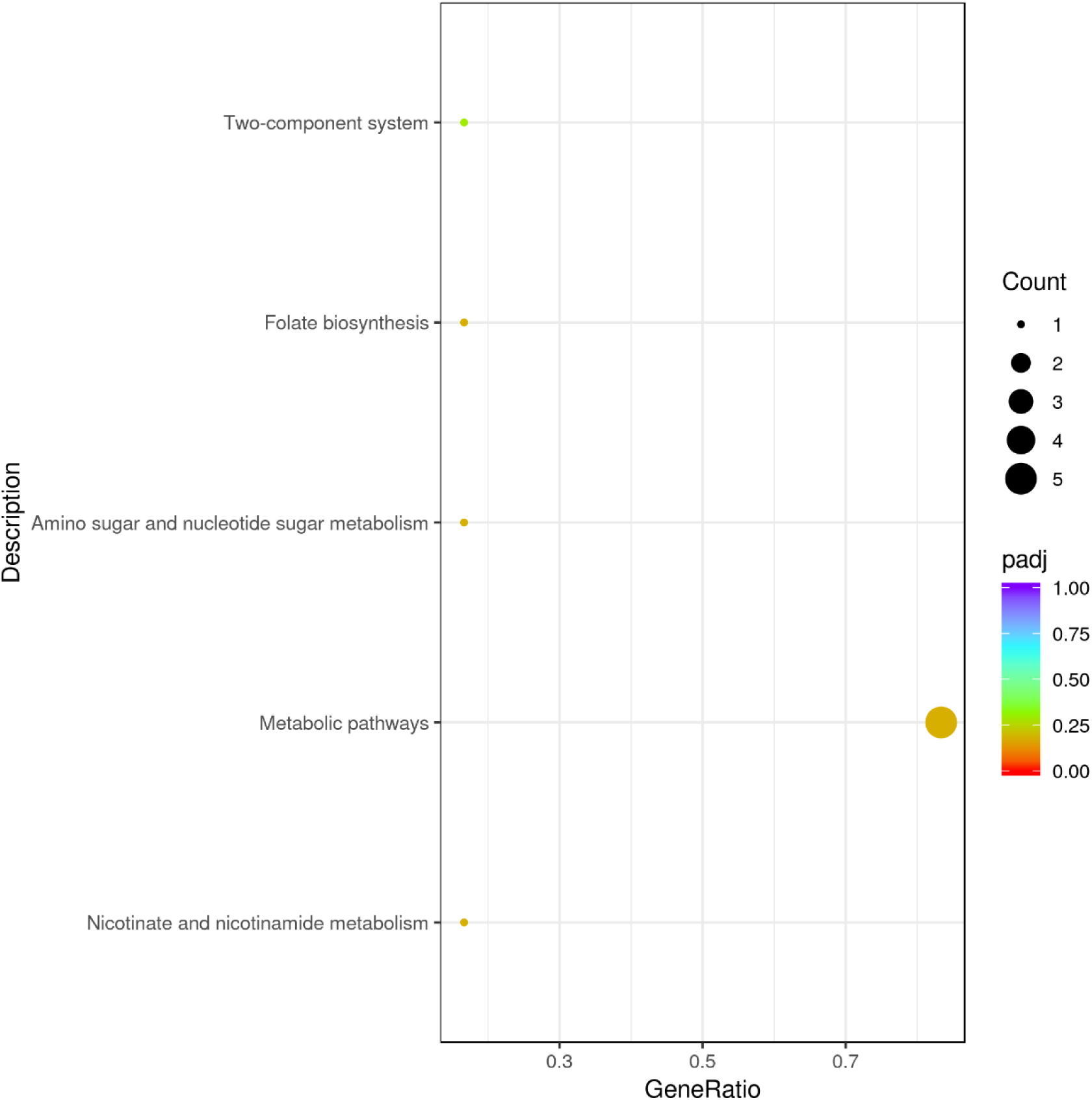
**Up-regulation KEGG pathways enrichment (scatter plot) based on the transcriptomic analysis of PE degradation by *Halomonas* sp. for 8 h.**

**Supplementary Fig. 42.**
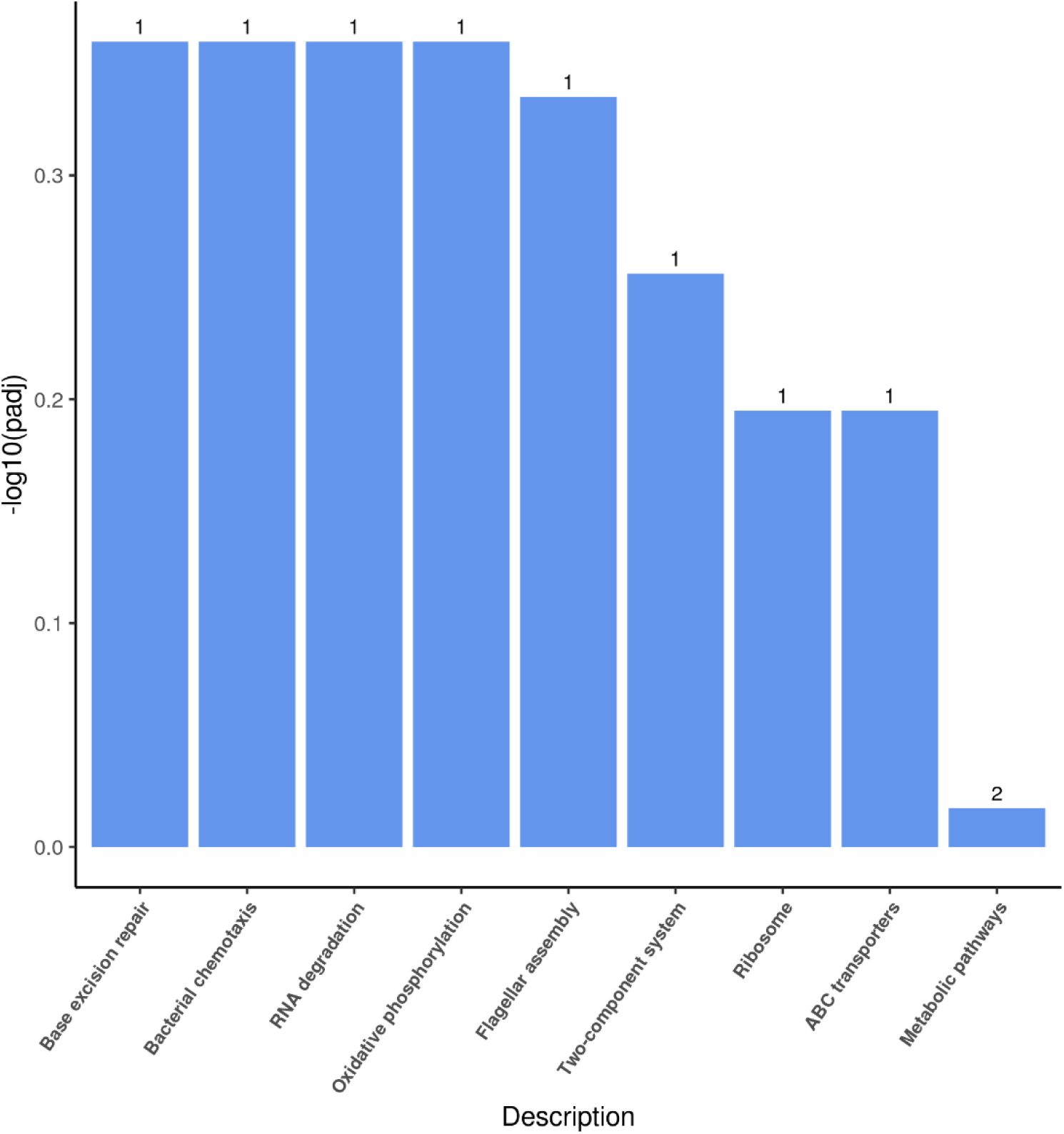
**Up-regulation KEGG pathways enrichment (histogram) based on the transcriptomic analysis of PE degradation by *Halomonas* sp. for 7 d.** The numbers above the column are corresponding genes number related to different pathways.

**Supplementary Fig. 43.**
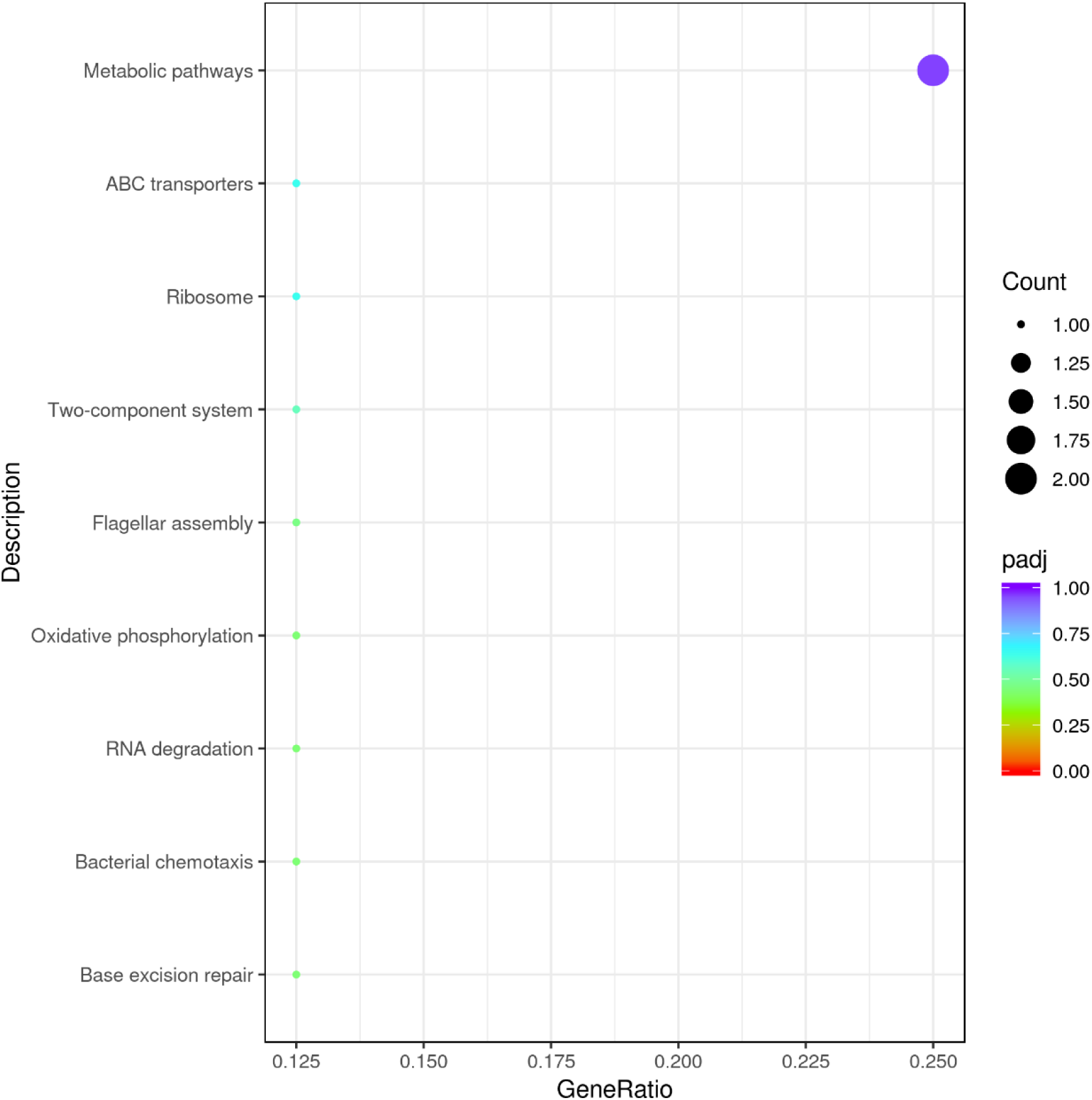
**Up-regulation KEGG pathways enrichment (scatter plot) based on the transcriptomic analysis of PE degradation by *Halomonas* sp. for 7 d.**

**Supplementary Fig. 44.**
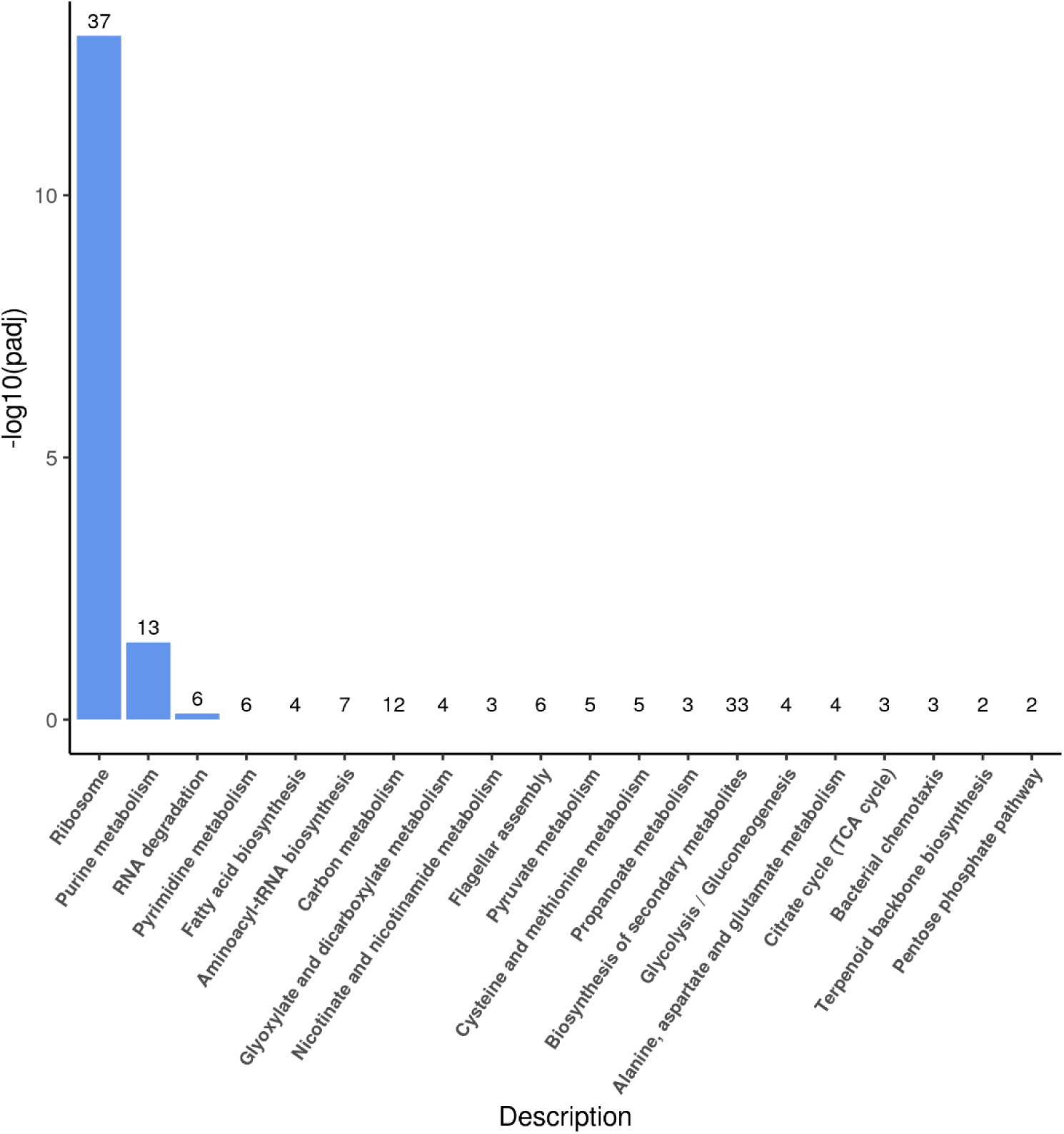
**Up-regulation KEGG pathways enrichment (histogram) based on the transcriptomic analysis of PE degradation by *Halomonas* sp. for 14 d.** The numbers above the column are corresponding genes number related to different pathways.

**Supplementary Fig. 45.**
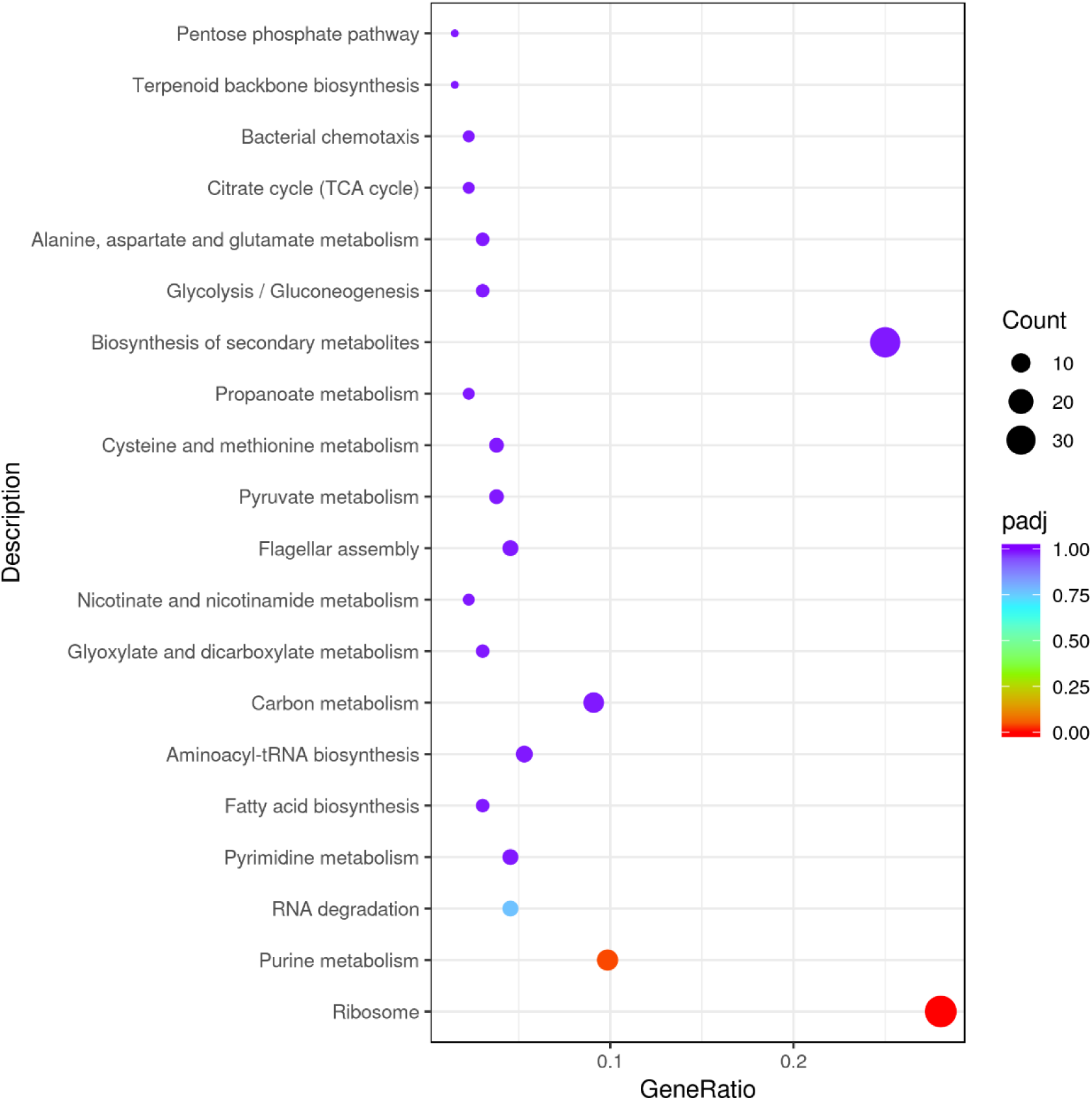
**Up-regulation KEGG pathways enrichment (scatter plot) based on the transcriptomic analysis of PE degradation by *Halomonas* sp. for 14 d.**

**Supplementary Fig. 46.**
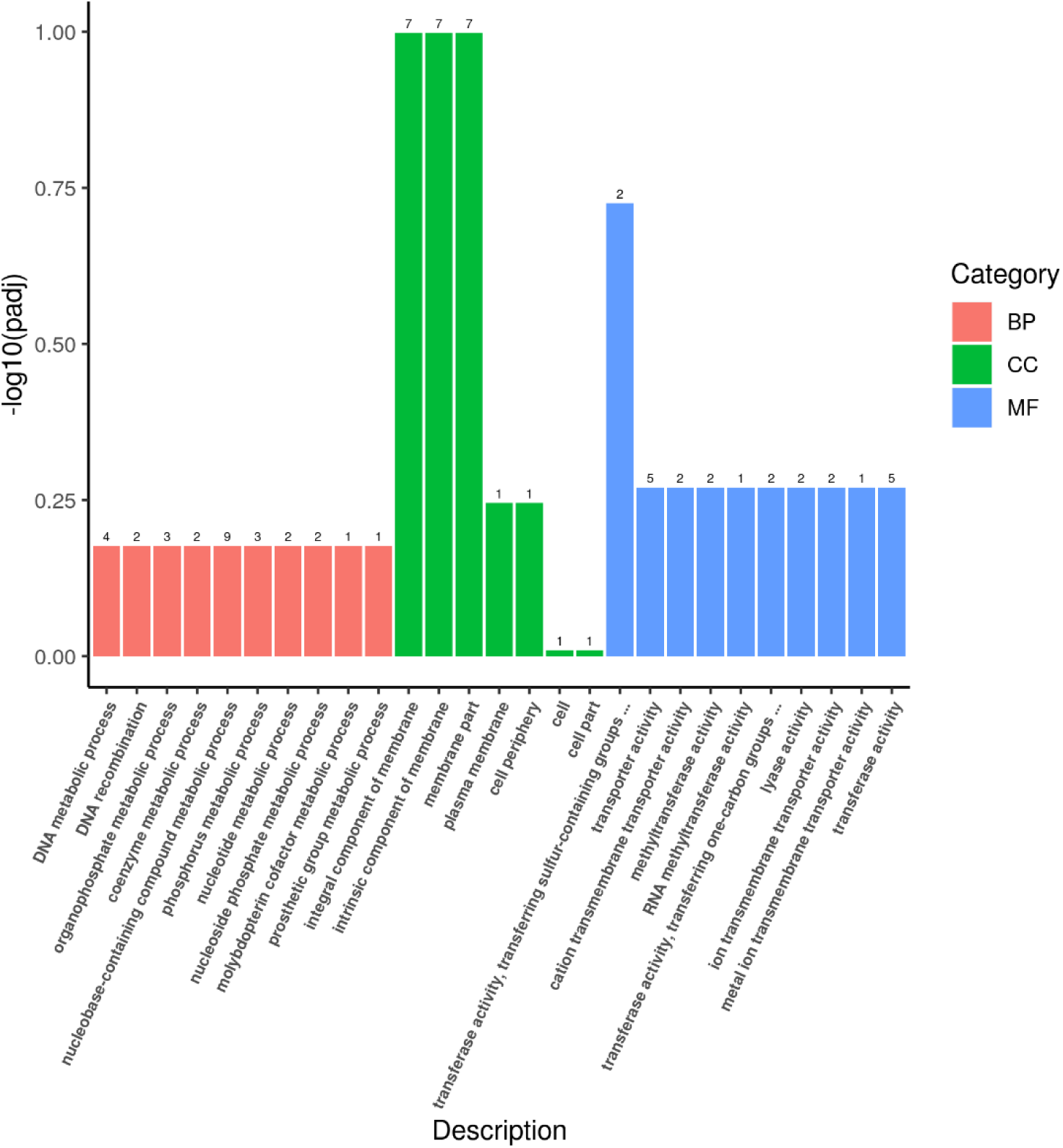
**Up-regulation Go enrichment (histogram) based on the transcriptomic analysis of PE degradation by *Halomonas* sp. for 8 h.** The numbers above the column are corresponding genes number related to different pathways.

**Supplementary Fig. 47.**
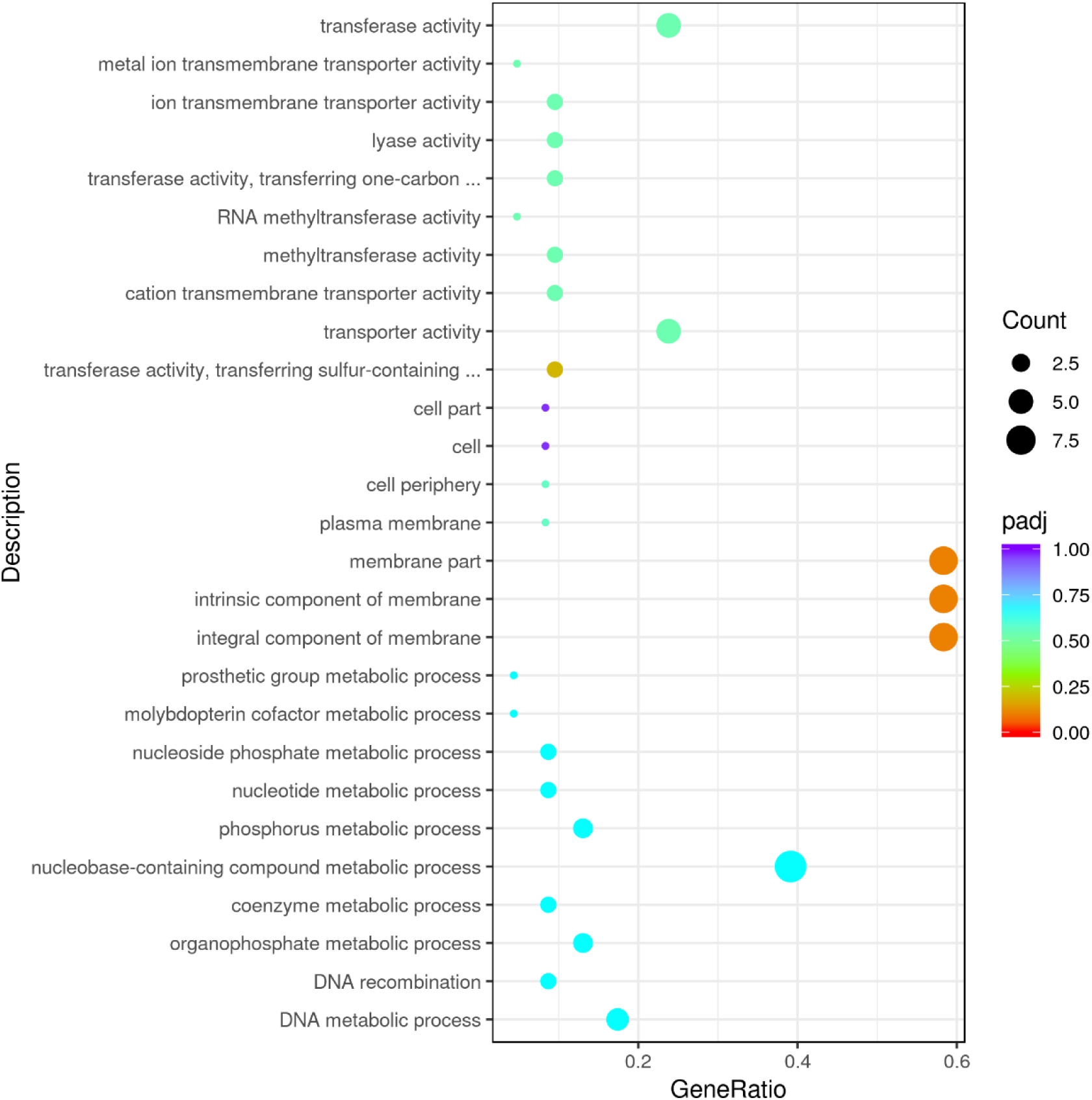
**Up-regulation GO enrichment (scatter plot) based on the transcriptomic analysis of PE degradation by *Halomonas* sp. for 8 h.**

**Supplementary Fig. 48.**
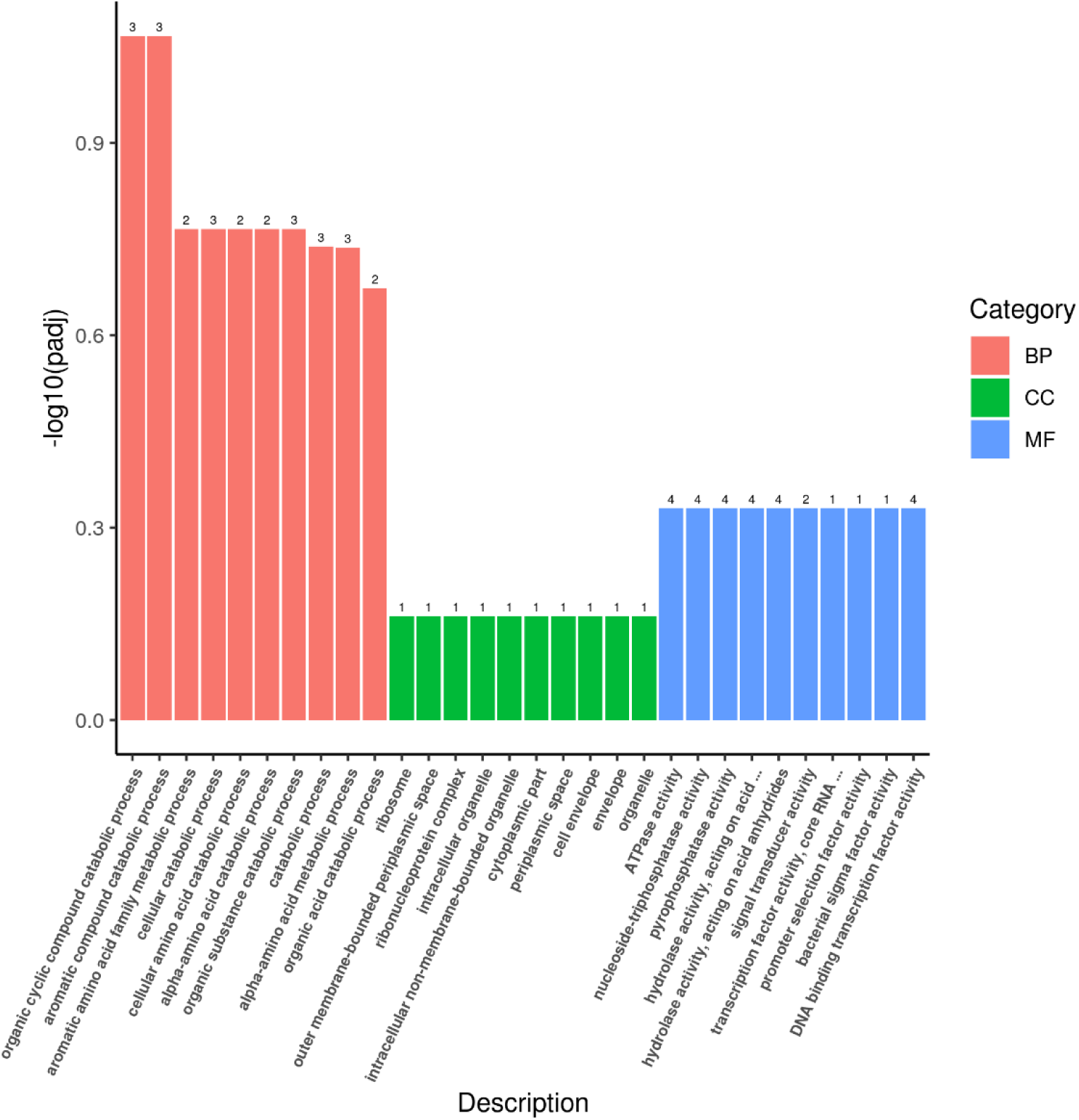
**Up-regulation Go enrichment (histogram) based on the transcriptomic analysis of PE degradation by *Halomonas* sp. for 7 d.** The numbers above the column are corresponding genes number related to different pathways.

**Supplementary Fig. 49.**
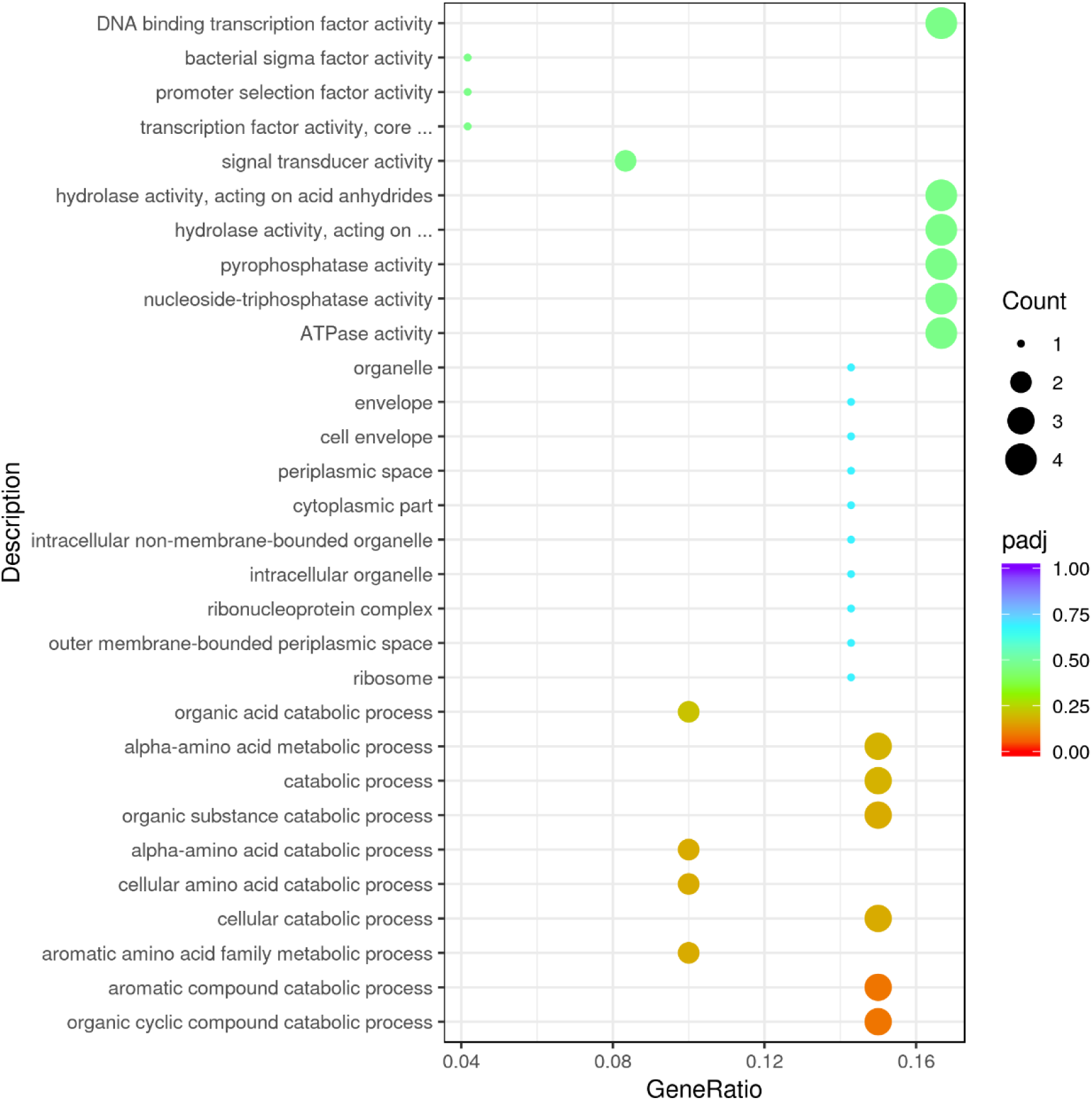
**Up-regulation GO enrichment (scatter plot) based on the transcriptomic analysis of PE degradation by *Halomonas* sp. for 7 d.**

**Supplementary Fig. 50.**
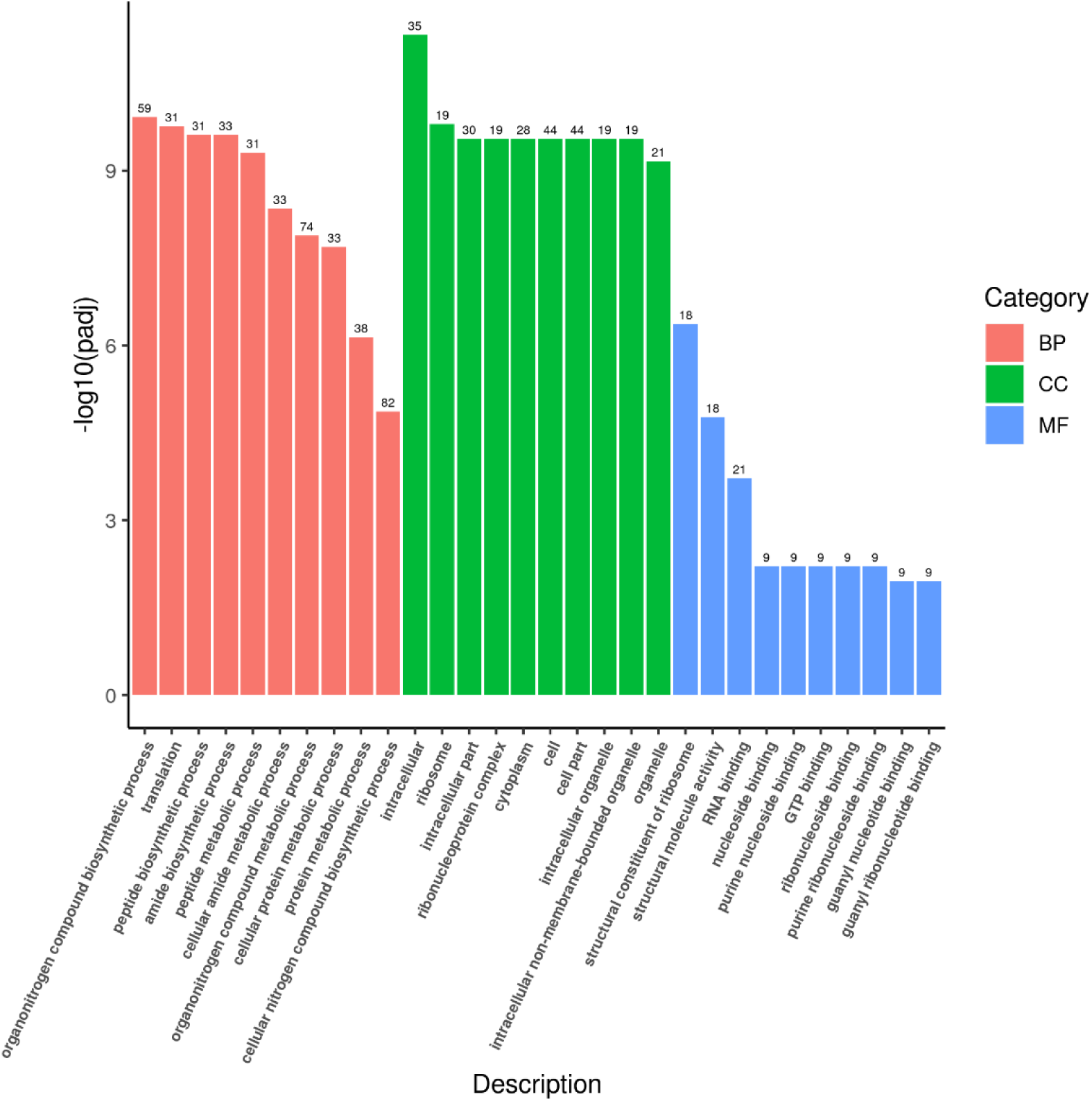
**Up-regulation Go enrichment (histogram) based on the transcriptomic analysis of PE degradation by *Halomonas* sp. for 14 d.** The numbers above the column are corresponding genes number related to different pathways.

**Supplementary Fig. 51.**
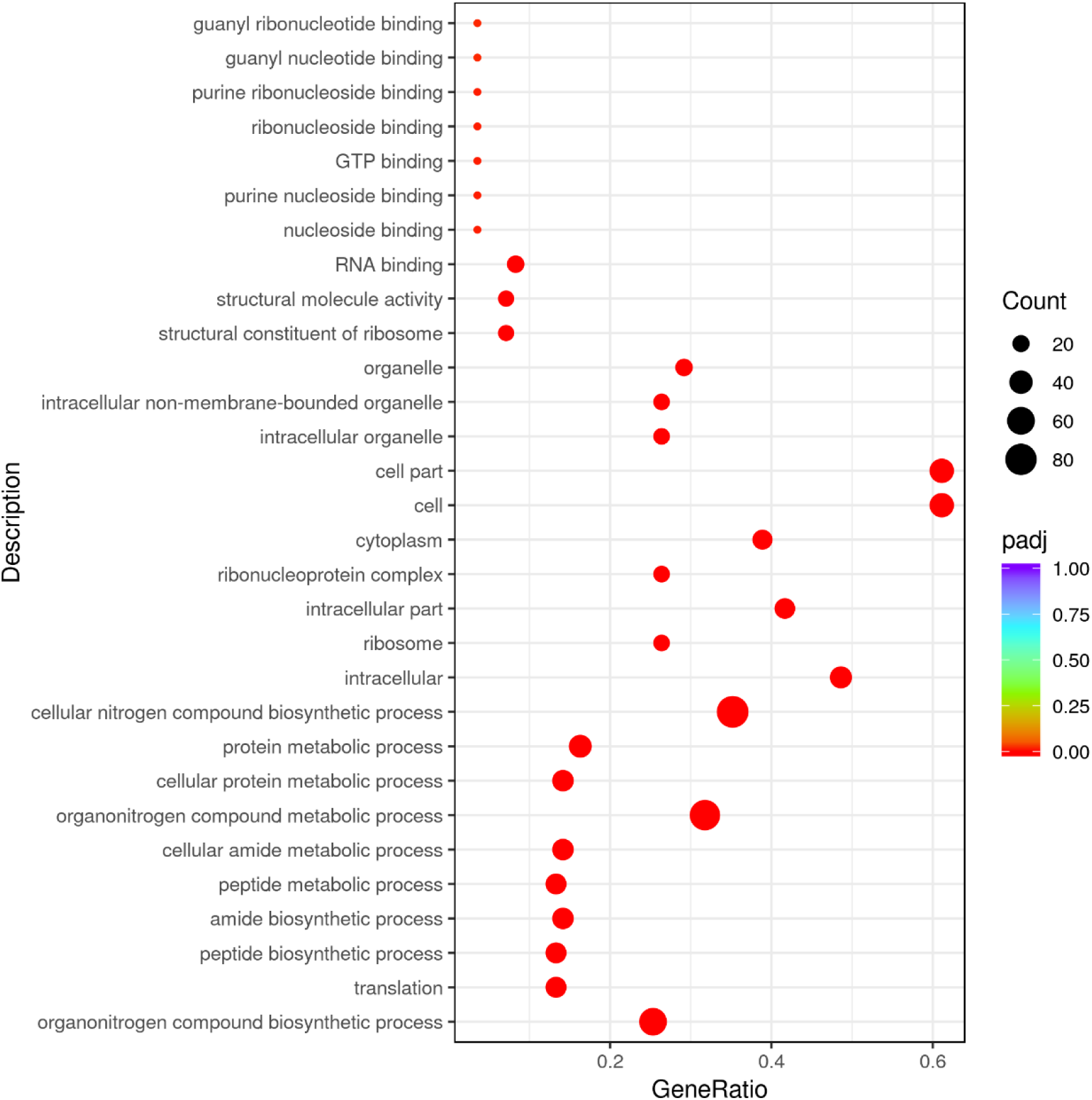
**Up-regulation GO enrichment (scatter plot) based on the transcriptomic analysis of PE degradation by *Halomonas* sp. for 14 d.**

**Supplementary Fig. 52.**
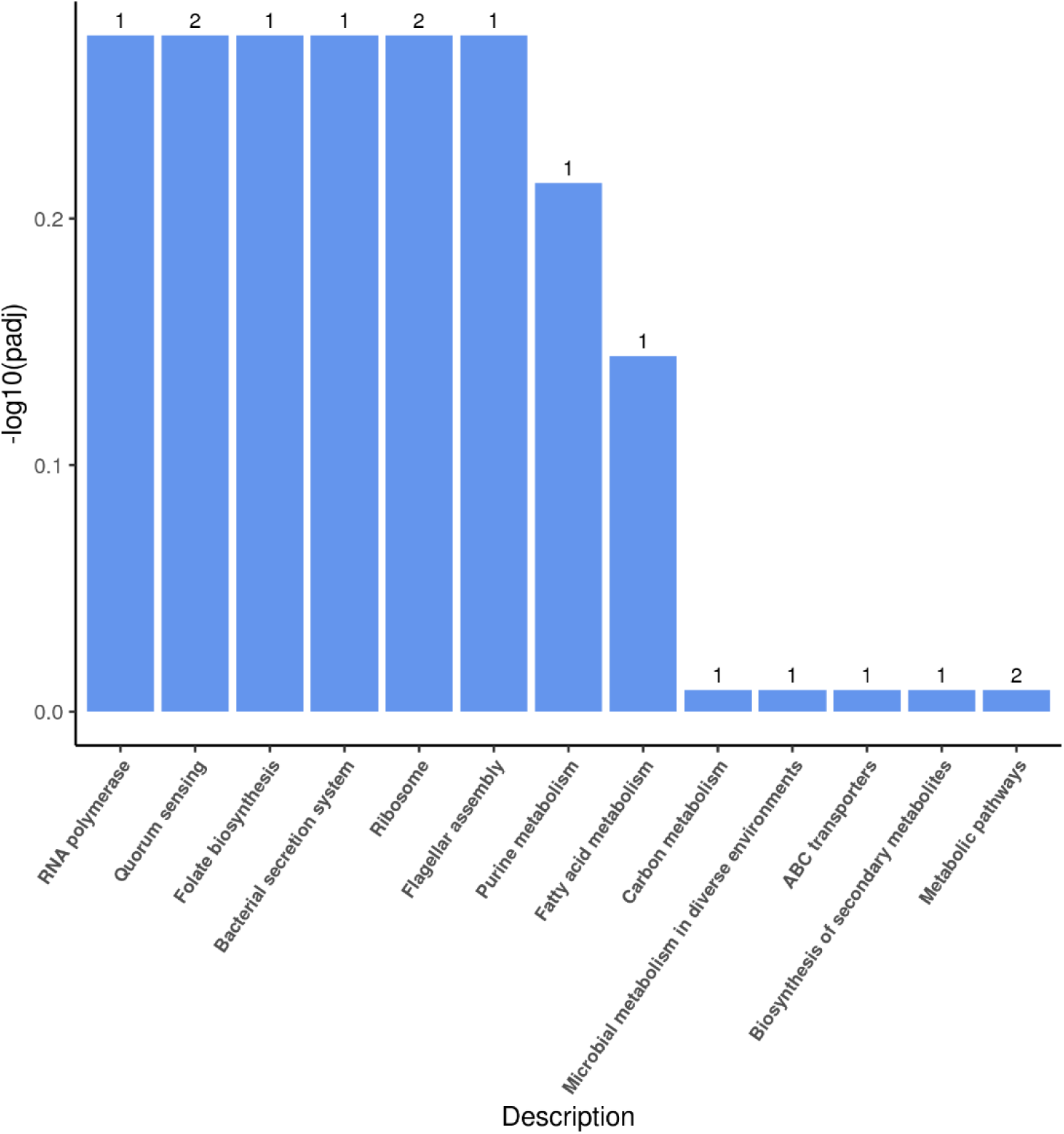
**Up-regulation KEGG pathways enrichment (histogram) based on the transcriptomic analysis of PE degradation by *Ochrobactrum* sp. for 8 h.** The numbers above the column are corresponding genes number related to different pathways.

**Supplementary Fig. 53.**
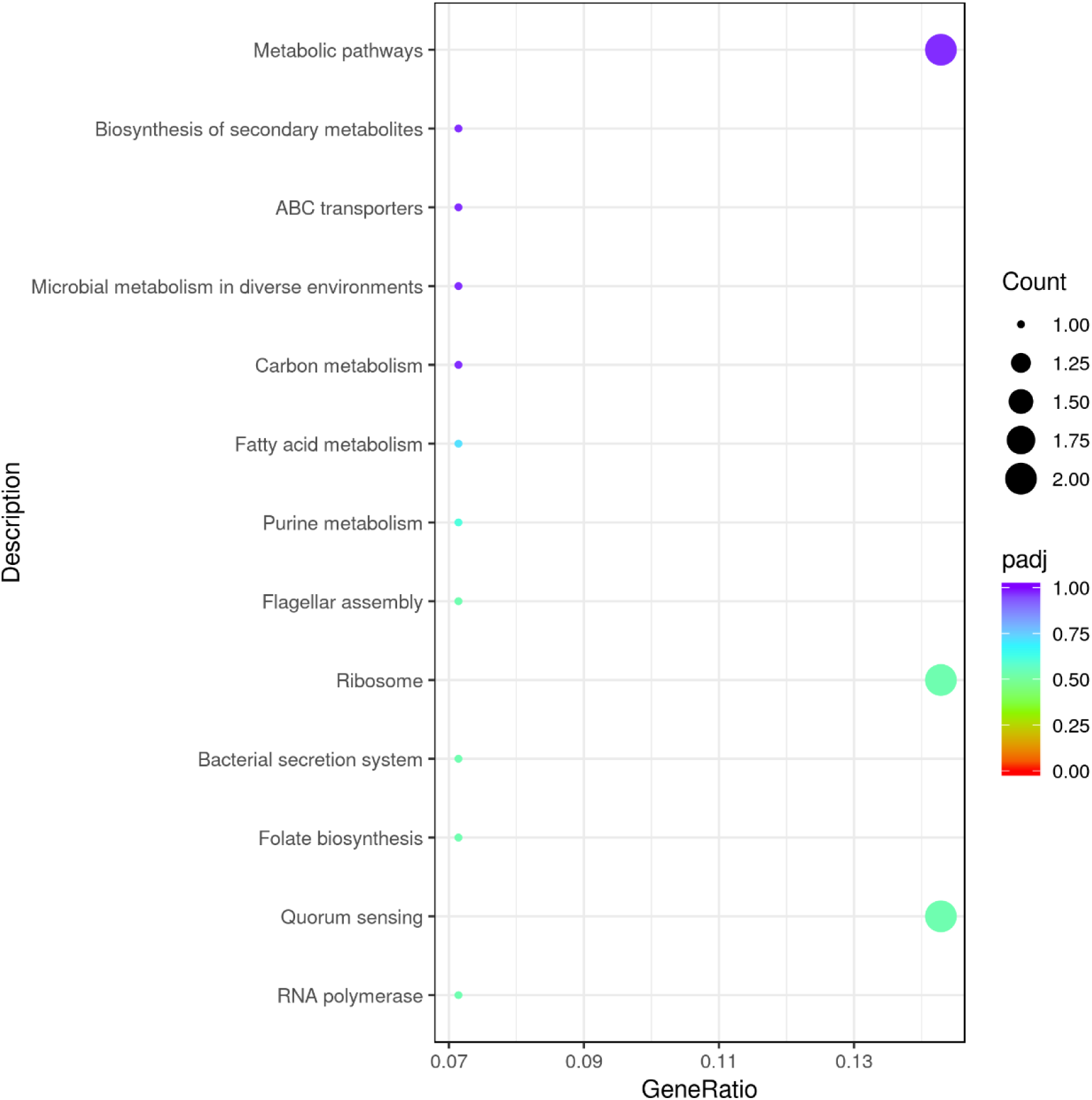
**Up-regulation KEGG pathways enrichment (scatter plot) based on the transcriptomic analysis of PE degradation by *Ochrobactrum* sp. for 8 h.**

**Supplementary Fig. 54.**
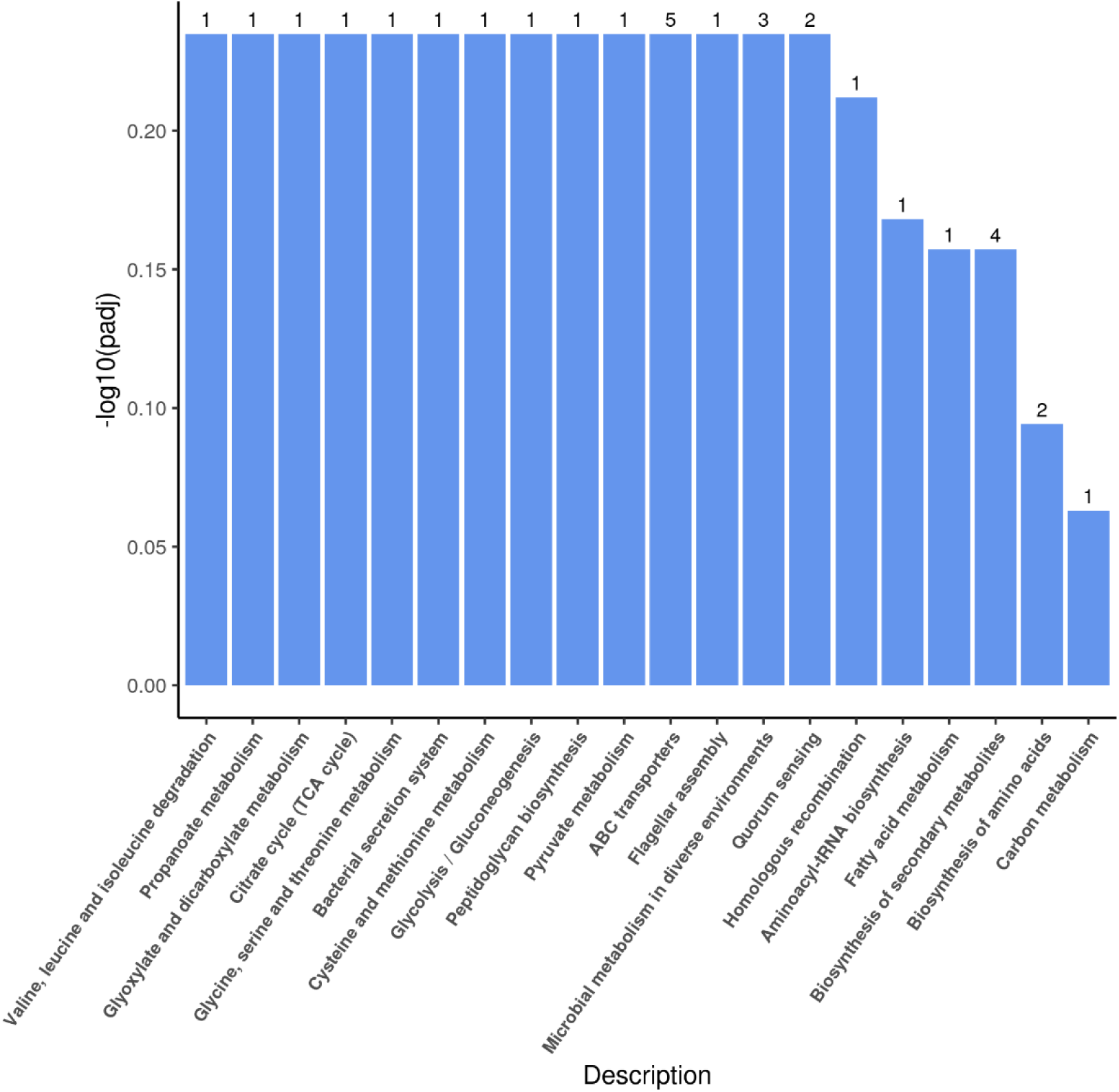
**Up-regulation KEGG pathways enrichment (histogram) based on the transcriptomic analysis of PE degradation by *Ochrobactrum* sp. for 7 d.** The numbers above the column are corresponding genes number related to different pathways.

**Supplementary Fig. 55.**
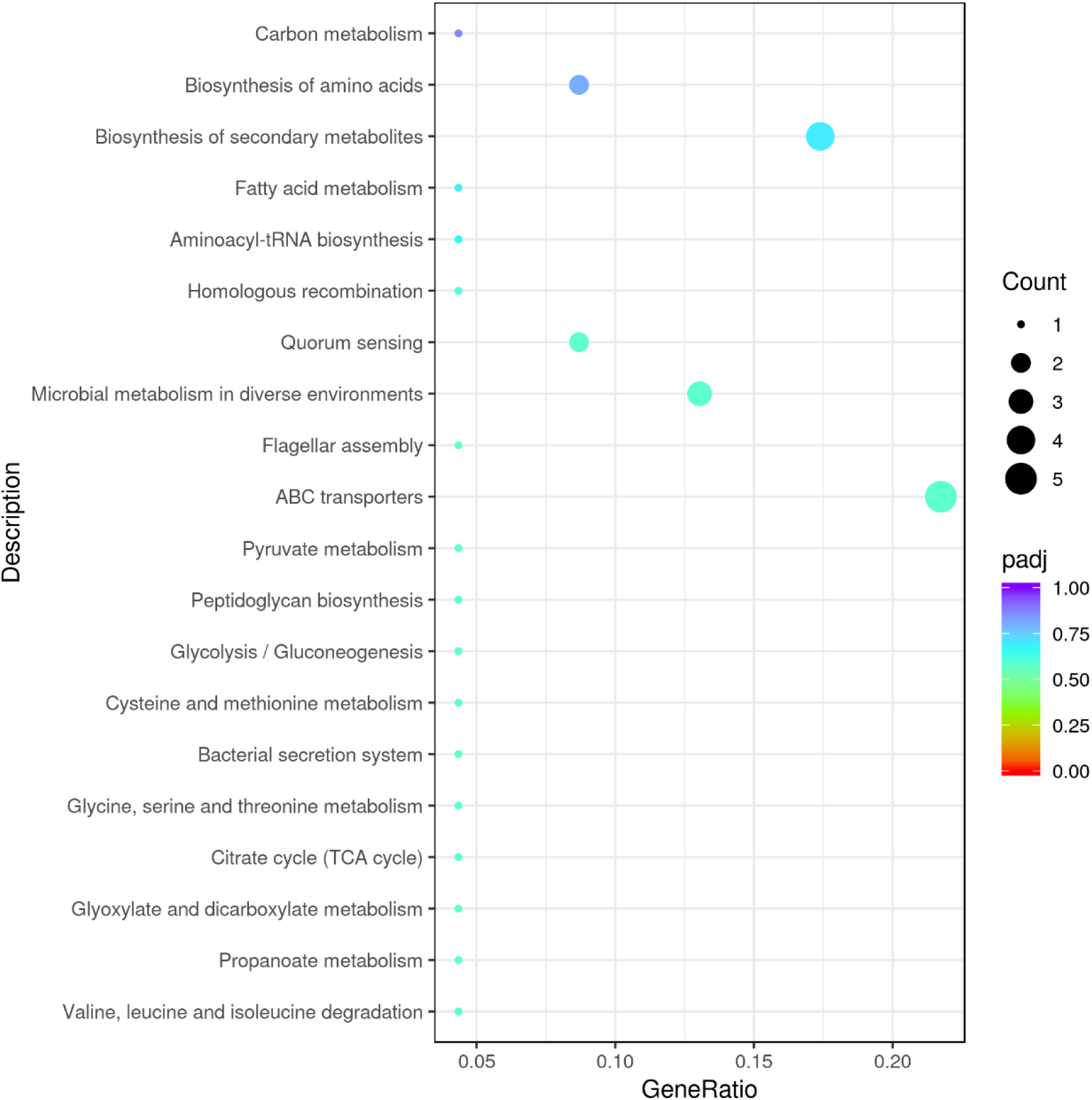
**Up-regulation KEGG pathways enrichment (scatter plot) based on the transcriptomic analysis of PE degradation by *Ochrobactrum* sp. for 7 d.**

**Supplementary Fig. 56.**
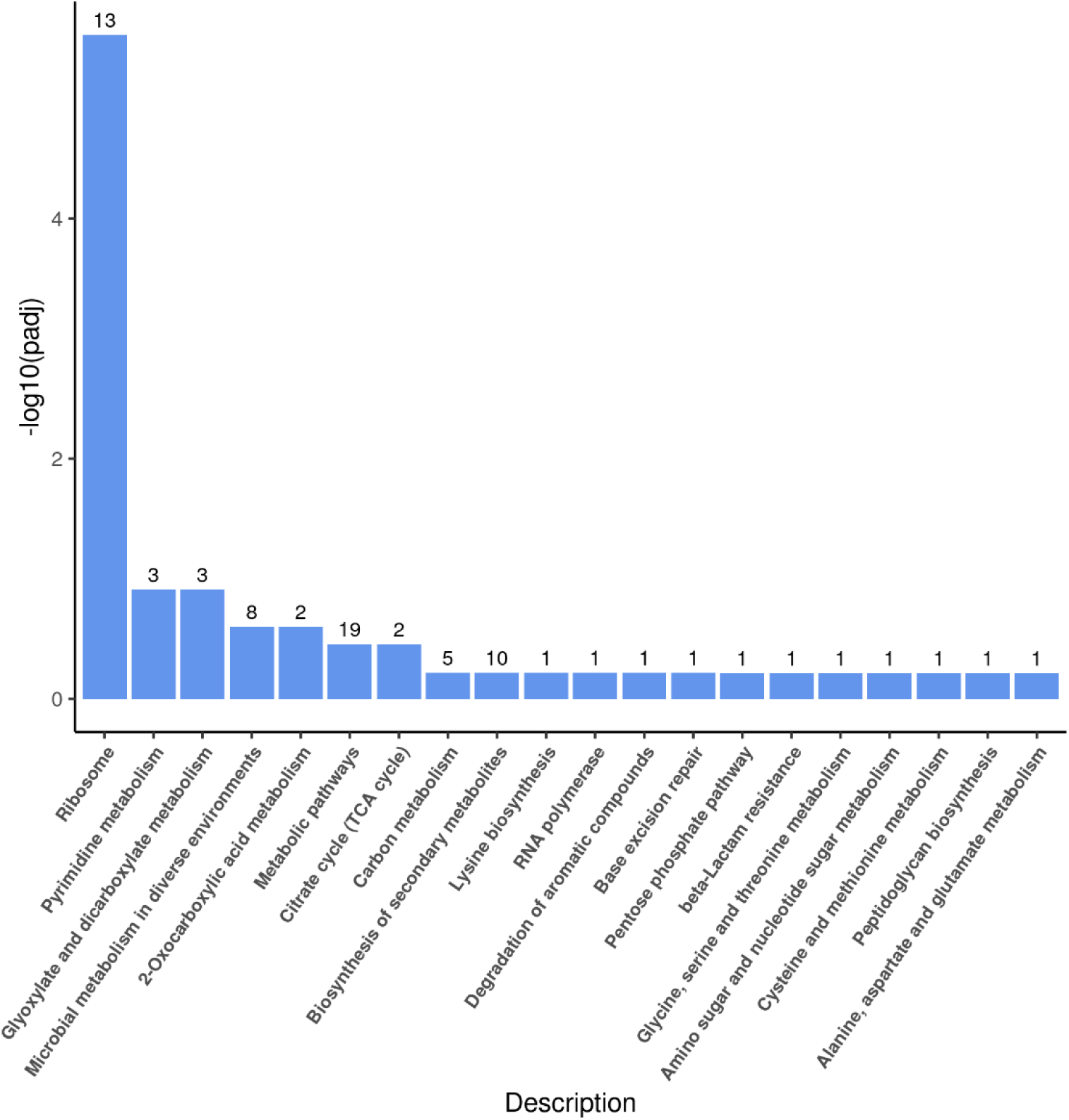
**Up-regulation KEGG pathways enrichment (histogram) based on the transcriptomic analysis of PE degradation by *Ochrobactrum* sp. for 14 d.** The numbers above the column are corresponding genes number related to different pathways.

**Supplementary Fig. 57.**
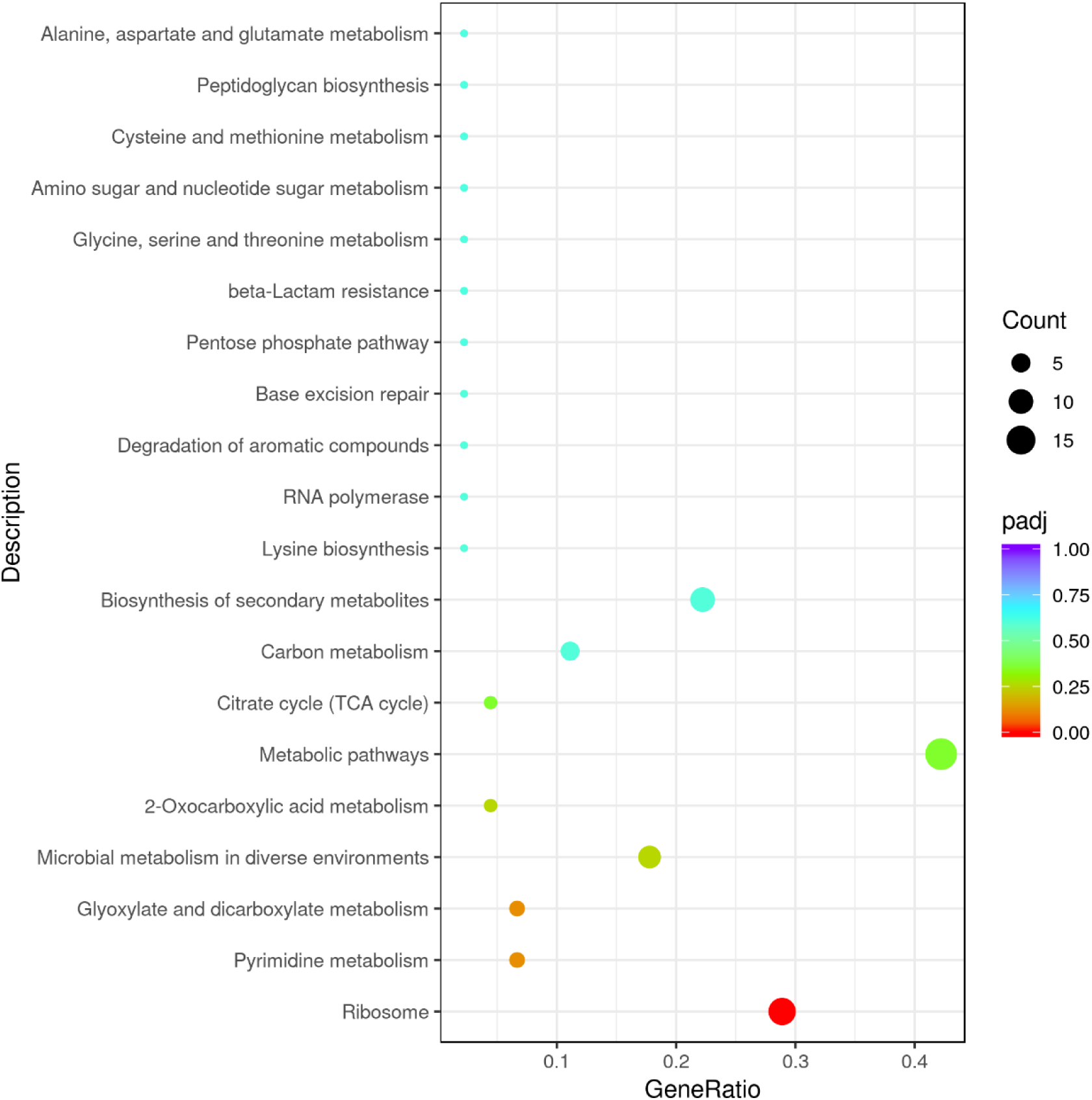
**Up-regulation KEGG pathways enrichment (scatter plot) based on the transcriptomic analysis of PE degradation by *Ochrobactrum* sp. for 14 d.**

**Supplementary Fig. 58.**
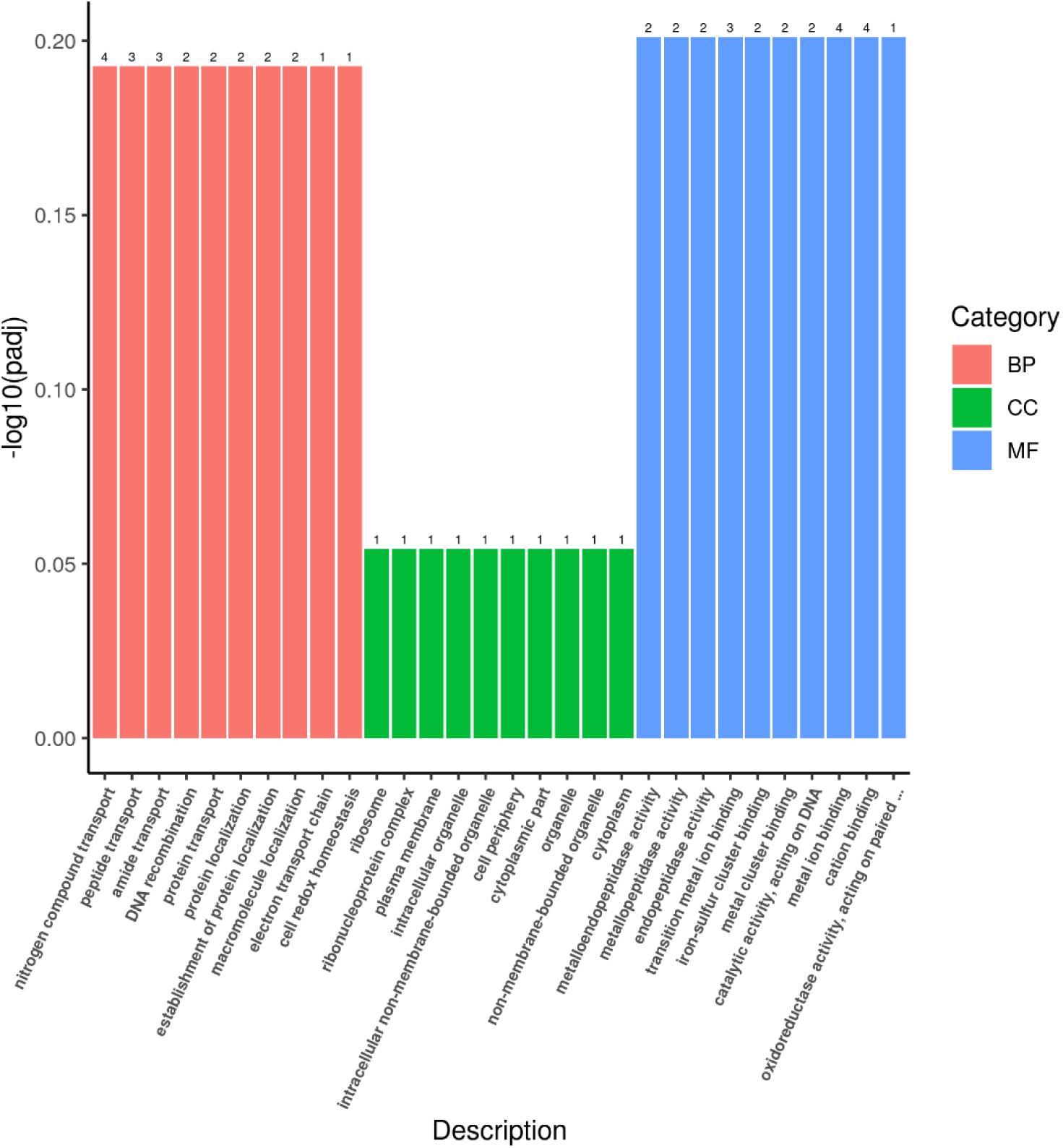
**Up-regulation Go enrichment (histogram) based on the transcriptomic analysis of PE degradation by *Ochrobactrum* sp. for 8 h.** The numbers above the column are corresponding genes number related to different pathways.

**Supplementary Fig. 59.**
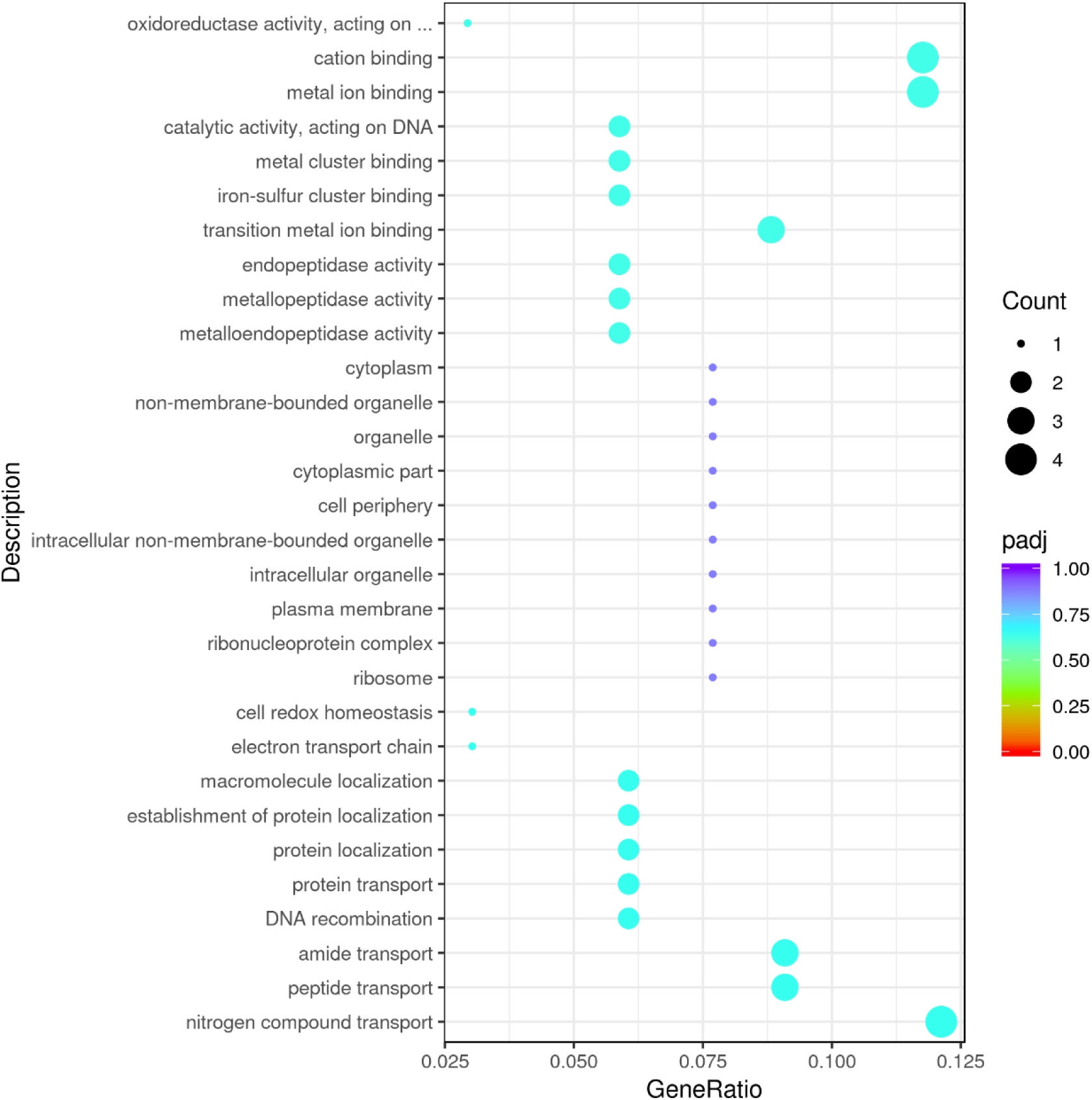
**Up-regulation GO enrichment (scatter plot) based on the transcriptomic analysis of PE degradation by *Ochrobactrum* sp. for 8 h.**

**Supplementary Fig. 60.**
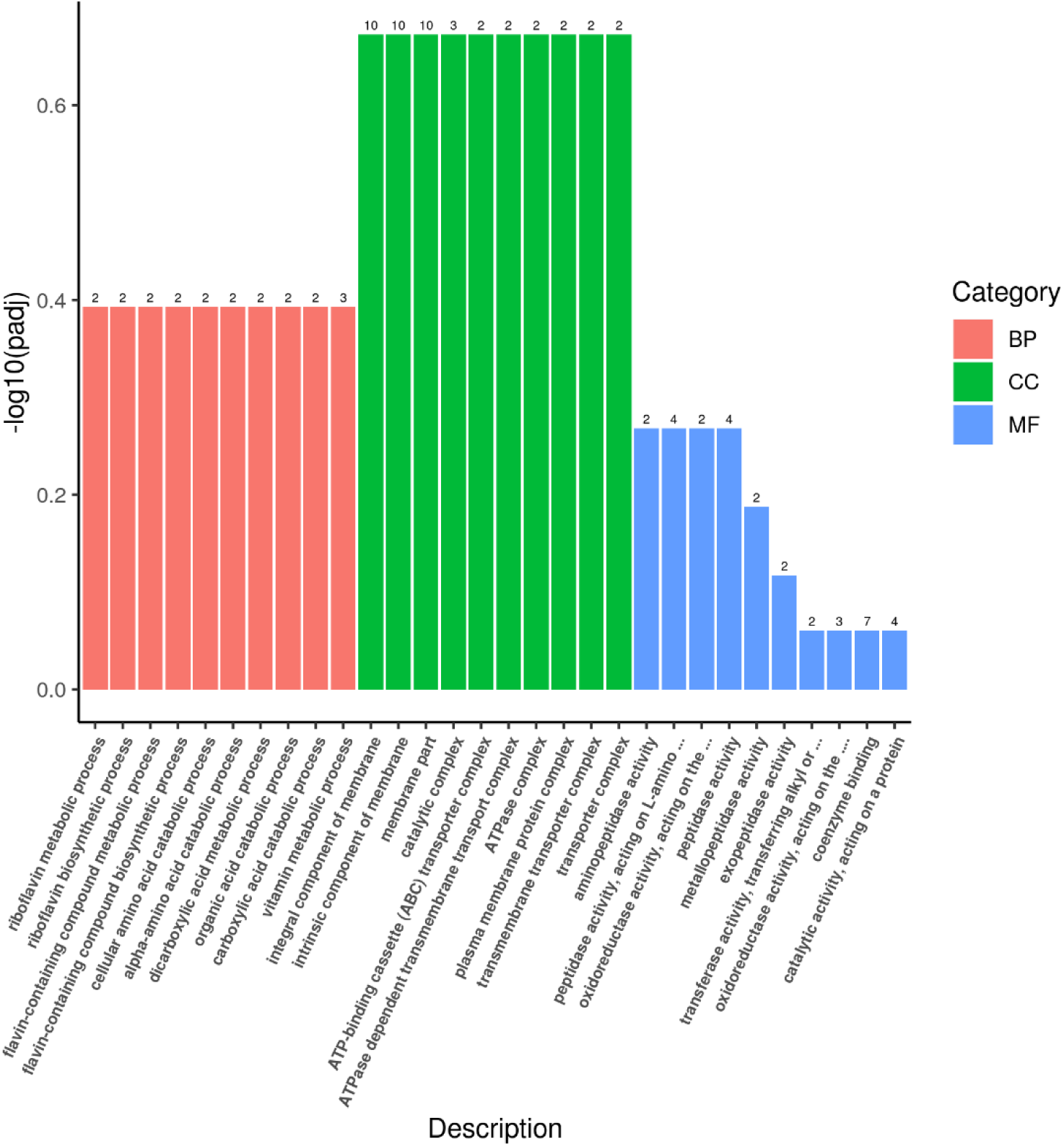
**Up-regulation Go enrichment (histogram) based on the transcriptomic analysis of PE degradation by *Ochrobactrum* sp. for 7 d.** The numbers above the column are corresponding genes number related to different pathways.

**Supplementary Fig. 61.**
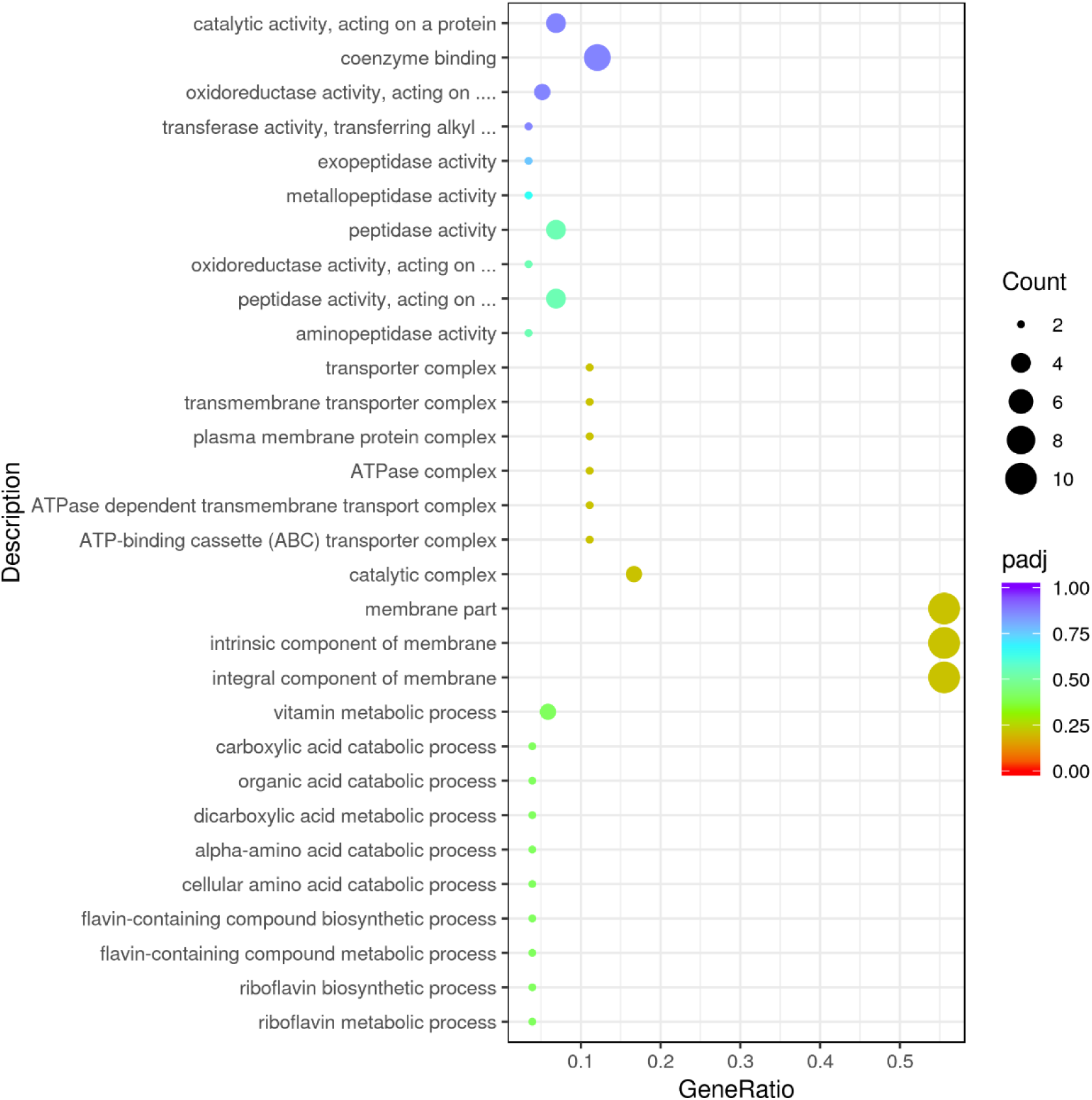
**Up-regulation GO enrichment (scatter plot) based on the transcriptomic analysis of PE degradation by *Ochrobactrum* sp. for 7 d.**

**Supplementary Fig. 62.**
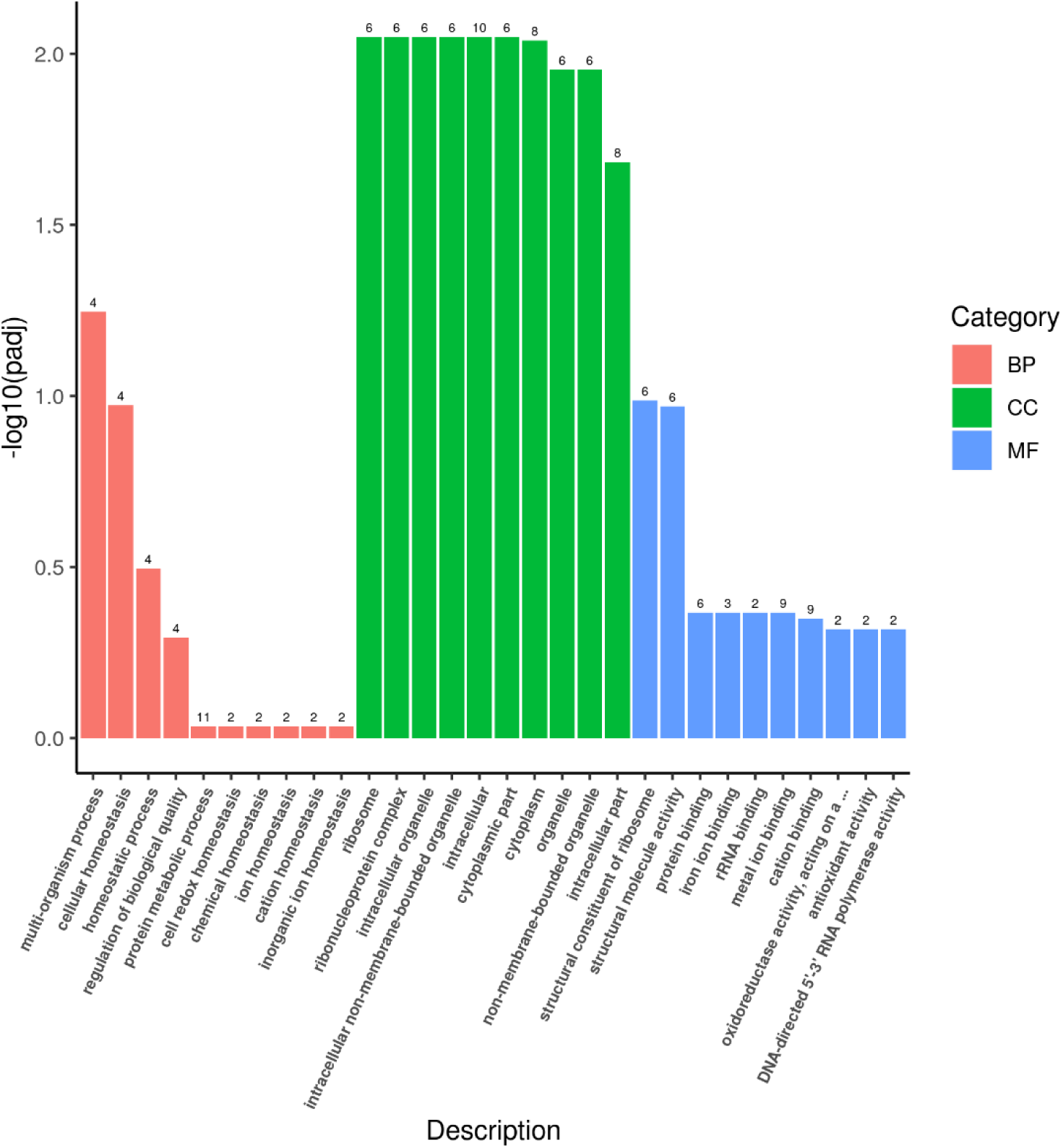
**Up-regulation Go enrichment (histogram) based on the transcriptomic analysis of PE degradation by *Ochrobactrum* sp. for 14 d.** The numbers above the column are corresponding genes number related to different pathways.

**Supplementary Fig. 63.**
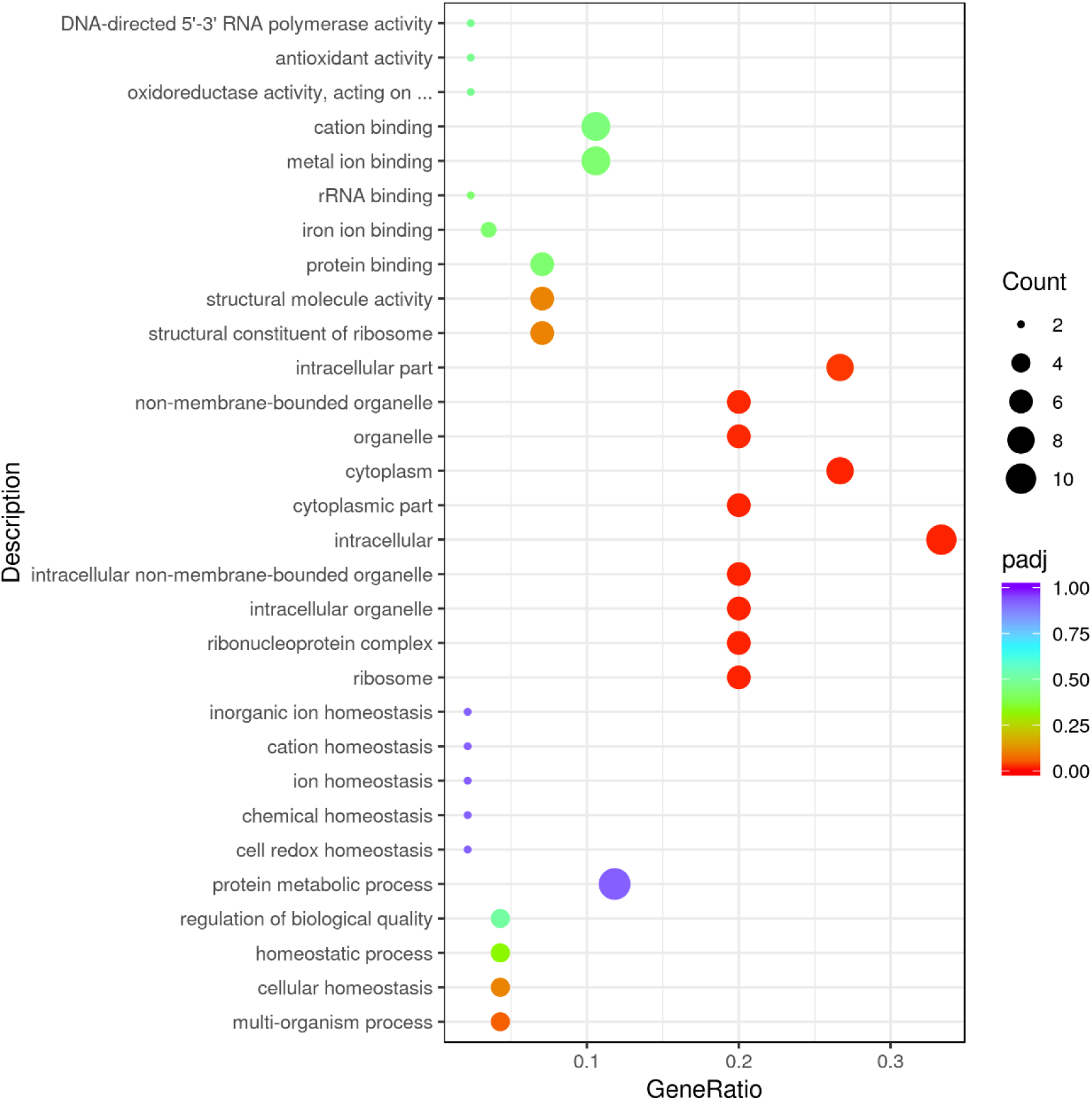
**Up-regulation GO enrichment (scatter plot) based on the transcriptomic analysis of PE degradation by *Ochrobactrum* sp. for 14 d.**

**Supplementary Table 1.** Absolute quantification analysis about 16S rRNA sequences of top 5 bacterial genera within the microbial community degrading plastics.

| genus | Total OTU<sup>*</sup> | E1 OTUsize<sup>#</sup> | E2 OTUsize<sup>#</sup> | E3 OTUsize<sup>#</sup> |
| --- | --- | --- | --- | --- |
| *Idiomarina* | 2456196232 | 1377935941 | 581578306 | 496681985 |
| *Marinobacter* | 1380072016 | 595840950 | 369029555 | 415201511 |
| *Exiguobacterium* | 886323611 | 424309595 | 265900937 | 196113079 |
| *Halomonas* | 122456072 | 60195881 | 36040715 | 26219476 |
| *Ochrobactrum* | 34634341 | 17227886 | 9263026 | 8143429 |

\*OTU: operational taxonomic unit. <sup>#</sup>Three biological replicates.

